# Structure and Function Relationships of Mucociliary Clearance in Human and Rat Airways

**DOI:** 10.1101/2023.12.24.572054

**Authors:** Doris Roth, Ayşe Tuğçe Şahin, Feng Ling, Niels Tepho, Christiana N. Senger, Erik J. Quiroz, Ben A. Calvert, Anne M. van der Does, Tankut G. Güney, Sarah Glasl, Annemarie van Schadewijk, Laura von Schledorn, Ruth Olmer, Eva Kanso, Janna C. Nawroth, Amy L. Ryan

## Abstract

Mucociliary clearance is a vital defense mechanism of the human airways, protecting against harmful particles and infections. When this process fails, it contributes to respiratory diseases like chronic obstructive pulmonary disease (COPD) and asthma. While advances in single-cell transcriptomics have revealed the complexity of airway composition, much of what we know about how airway structure impacts clearance relies on animal studies. This limits our ability to create accurate human-based models of airway diseases. Here we show that the airways in female rats and in humans exhibit species-specific differences in the distribution of ciliated and secretory cells as well as in ciliary beat, resulting in significantly higher clearance effectiveness in humans. We further reveal that standard lab-grown cultures exhibit lower clearance effectiveness compared to human airways, and we identify the underlying structural differences. By combining diverse experiments and physics-based modeling, we establish universal benchmarks to assess human airway function, interpret preclinical models, and better understand disease-specific impairments in mucociliary clearance.

## INTRODUCTION

Mucociliary clearance (MCC) is a critical mechanical barrier mechanism of the human airways^1–3^. In MCC, the beating of specialized multiciliated cells propel a layer of mucus across the epithelial surface, effectively trapping and removing inhaled particles and pathogens^4,5^. The mucus is produced by the submucosal glands and a variety of secretory cells interspersed among ciliated cells. These secretory cells include mucin-secreting goblet cells, club cells and their intermediate stages^6^. The composition of secretory cell types has a profound impact on the rheology and flowability of mucus^7–9^. Failure of MCC contributes to the debilitating pathophysiology of many respiratory diseases, including chronic obstructive pulmonary disease (COPD), primary ciliary dyskinesia, asthma, and cystic fibrosis^10,11^. However, our understanding of how diseases impair MCC remains limited by an incomplete knowledge of the mechanics that link secretory and ciliated cell organization to MCC in native human airway tissues^3^ since most *in vivo* and *ex vivo* data has come from animal models^12–14^. Obtaining data on live ciliary beat and clearance function in human tissue poses challenges and remains exceedingly rare in the literature^15,16^. Furthermore, the organization of secretory and ciliated cells at the epithelial surface is underexplored, as conventional histology typically reveals cross-sectional rather than surface-lining tissue architecture. While providing important insights about cellular heterogeneity, transcriptional profiling commonly does not resolve spatial organization^17,18^ and may not accurately reflect protein expression^19^.

The lack of such crucial data limits not only our understanding and capability to model human MCC but may also undermine correct diagnosis and early intervention in disease. For instance, a recent study revealed that while luminal surface ciliation in the mouse trachea is relatively low (approximately 40%), the specific distribution and orientation of ciliary beat ensure robust and efficient particle clearance^14^. However, the authors could only speculate whether these design principles directly translate to humans, where such quantitative insights could help refine treatments for conditions impairing ciliary beat, such as PCD^20^, or causing loss of ciliated cells, such as asthma^21^. The limited human-relevancy of animal studies has spurred the development of innovative *in vitro* human airway epithelial models^22–24^. Nevertheless, without quantitative benchmarks of MCC in humans, the ability of these models to accurately represent human physiology remains unproven. Furthermore, the proportions of ciliated and various secretory cell types are believed to vary along the airway tree within and between humans and rats^25^. It remains unclear whether this heterogeneity results in species-specific regional differences in MCC, which is pertinent for studying diseases that alter the regional proportions of airway epithelial cell types^26^.

To address these gaps, we have established a quantitative map of the distribution of ciliated, club, and goblet cells along the surface lining of the human and rat airway tree. We have evaluated species-specific differences in the associated particle clearance and utilized these data to develop quantitative metrics and physics-based computational models capturing key MCC characteristics in lower airway epithelia. Finally, we deploy our comprehensive quantitative framework to benchmark the structural and functional properties of a variety of *in vitro* human airway epithelial culture conditions against native human airway epithelia. The data presented significantly enhances our understanding of MCC mechanics and, by providing means of MCC quantification and quality control, will improve the translational potential of *in vitro* models for studying diseases and therapeutic interventions affecting MCC.

## RESULTS

### Tracheo-bronchial ciliation is higher in human airways compared to rat

Live ciliary beat and particle clearance^15,27^ was measured in freshly isolated airway epithelial tissue originating from the ventral wall of the respiratory tree branching generation (BG)0 through BG6 derived from 4 human whole lungs with no prior history of chronic lung disease, and BG0 through BG5 from healthy rat whole lungs (**Fig. 1a**; for detailed donor information and numbers see **Supplementary Tables 1 & 2**). The human BG6 samples were at, or below, 2 mm in diameter (**Supplementary Fig. 1**) and therefore constituted the transition to small airway morphology^28^. While such a definition of small airways does not exist for rodents, our analysis covered a comparable fraction of the conducting airways along the proximal-distal axis in both systems (7 BGs of 16 BGs in humans^29^, and 6 BGs of 14 BGs in rats^30^). Subsequently, we quantified the luminal (i.e., surface-lining) cell type composition in these samples and, to increase donor numbers, in additional fixed human bronchial rings. Where unknown, their BG was estimated based on their diameters (**Supplementary Table 1)** using a calibration curve **(Supplementary Fig. 1).** The percentage of cells at the mucosal surface expressing cell-specific markers was quantified using mucin 5AC (MUC5AC) to mark goblet cells, secretoglobin family 1A member (SCGB1A1) to label club cells, and acetylated alpha-tubulin (ATUB) to stain ciliated cells^4^ (**Fig. 1b-c**). Luminal cell boundaries were revealed by F-actin staining of the cytoskeleton, allowing us to segment the cell outlines using image processing and thereby determine total cell numbers in each field of view. By overlaying the signal of each marker onto the cell outlines, we computed the percentages of different cell types and their overlaps (See Methods and **Supplementary Figs. 2 & 3**). The results are summarized in **Fig. 1d**, for details see **Supplementary Tables 3 & 5.** Intriguingly, human airways exhibited a robustly high ciliated cell proportion, or cilia coverage, of 86 ± 9% (mean±STD) across all analyzed BGs, whereas cilia coverage in rat airways gradually increased along the airway tree, ranging from 49 ± 12% in trachea to 92 ± 2.3% in BG5. Of note, the trachea in rat exhibited a pattern of alternating lower (ca. 30-40%) and higher (ca. 45 - 60%) ciliation levels, corresponding to cartilage rings and the interspersed tracheal ligaments, respectively^31^. We did not observe such regular ciliation patterns in the human airways (**Supplementary Fig. 4)**. The dense luminal ciliation of the human airways contrasted with results obtained from classic histology or gene expression data, which, likely by including both luminal and subluminal cell populations in the analysis, have suggested a lower ciliation level of 30-50% in human trachea and increasing ciliation in higher BGs^25,32–35^ (**Supplementary Fig. 4**). Our data hence suggest that surface ciliation cannot be easily predicted from bulk tissue percentages. On the other hand, our analysis of secretory cell levels matched trends reported in these studies. In all human BGs examined, 10-20% of luminal cells were positive for either MUC5AC, SCGB1A1, or both. Within this secretory cell population, the proportion of MUC5AC+ SCGB1A1- (goblet) and MUC5AC-SCGB1A1+ (club) cells varied as a function of BG (**Fig. 1d**, inset). Goblet cells dominated the trachea-bronchial region whereas club cells became increasingly frequent in BG3 and higher. On average, 15 and 30% of the labeled secretory population were MUC5AC+ SCGB1A1+ (double-positive hybrid/transitionary) cells, consistent with transcriptional and histological studies^32,36,37^ (for additional staining see **Supplementary Fig. 6A & B**). A comparable analysis was inconclusive for the rat secretory cells due to the spatially variable, but overall low (<10%), abundance of cells positive for MUC5AC and/or SCGB1A1, which matches literature^25^. As noted by others^31^, there was spatial variability of secretory cell density (**Supplementary Fig. 6C)**. Since rat airways are thought to contain a high abundance of so-called “serous” secretory cells^25^ for which no protein markers have been reported, additional secretory cells could be present.

**Figure 1:**
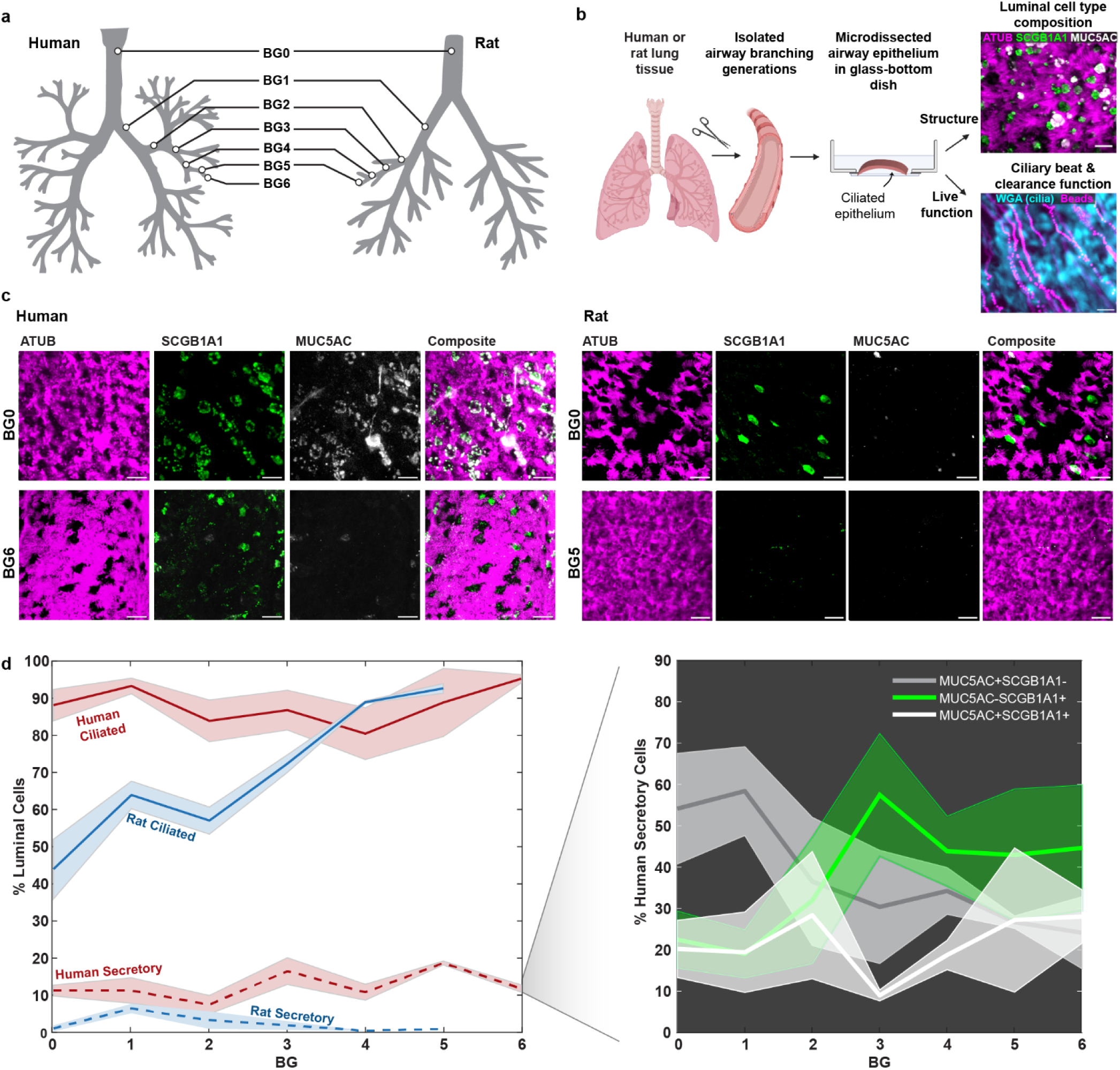
Luminal epithelial cell type composition differs along the human and rat airway tree. a, Airway branching generations in human and rat investigated in this study included BG0-6 in humans and BG0-5 in rats. b, Workflow for imaging luminal epithelial cell type composition and ciliary beat and clearance function in airway samples. Schematic: *Created in BioRender. Nawroth, J. (2025)* https://BioRender.com/a01i578). c, Example IF staining of cilia (ATUB, magenta) and secretory cells (SCGB1A1, green; MUC5AC, grey) in human and rat airway epithelium in BG0 (trachea) and BG5/6. Scale bar: 20 µm. d, Quantification of luminal cell proportions labeled with ATUB (ciliated cells) or with MUC5AC and/or SCGB1A1 (secretory cells) as a function of airway branching generation in human and rat airway epithelium. Inset: Percentage of human secretory cell population positive for only MUC5AC (grey), only for SCGB1A1 (green), or for both (white) as a function of branching generation. Solid line: mean, shaded region: SEM. Numbers of human donors (2-3 FOVs each): BG0, n=3; BG1, n=3 ; BG2, n=5 ; BG3, n=3 ; BG4, n=7 ; BG5, n=2 ; BG6, n=2 ; Numbers of rat donors (2-3 FOVs each): BG0, n=5 ; BG1, n=3 ; BG2, n= 2; BG4, n= 1; BG5, n=1. For full donor information see **Supplementary Tables 1 & 2**. Source data for panel d are provided as a Source Data file.

### Human airways achieve greater clearance per ciliary beat than rat airways

The differences in cilia coverage between species suggested differences in particle clearance, which we proceeded to explore by measuring live ciliary beat and fluorescent bead transport in the densely ciliated human airways (BG0 to BG6) compared to the more sparsely ciliated rat airways (BG0 and BG1). (**Fig. 2a-b, Supplementary Videos S1 & S2)**. These regions correspond to the trachea, main stem bronchi, and bronchial regions in humans and rats. Matching cell type composition analysis, we sampled multiple fields of views along the ventral airway tube. As the human explant tissues had been submerged in buffer for many hours prior to live recordings, the naturally occurring periciliary liquid and mucus layers were likely diluted or removed. Hence, in order to enable a direct comparison of ciliary beat and clearance between human explants and other samples, we chose to gently wash all apical surfaces to remove any build-up of mucus, and to then image the samples submerged in aqueous buffer. Ciliary beat frequency (CBF) recorded at room temperature was 2.6 ± 0.5 Hz in human tissue and 4.5 ± 1.3 Hz in rat tissue. Associated particle clearance speed reached 16.5 ± 8.2 µm s^−1^ in humans and 4.8 ± 2.0 µm s^−1^ in rats. Therefore, clearance speed was significantly higher in humans despite significantly lower CBF than in rat tissues. To assess this imbalance further, we derived the normalized “clearance speed per beat” (CPB) by dividing the average clearance speed by the average CBF for each field of view. CPB measures how far a particle is transported per ciliary beat cycle and has the units µm per beat. CPB can be considered a measure of stroke effectiveness and is independent of CBF. Our analysis revealed a significantly higher CPB in human tissue (6.2 ± 2.5 µm per beat) compared to rat tissue (1.1 ± 0.3 µm per beat) (**Fig. 2c**). We also investigated particle clearance directionality *D* as a function of distance *R*, defined as 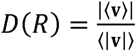, where 〈〉 indicates the mean, || indicates the magnitude, and v indicates flow velocity. *D(R)* ranges from 0 to 1, where a number near zero indicates highly convoluted flow and a number near 1 indicates straight and unidirectional flow, and it decays with increasing distance. Each trace was fitted with a decaying exponential, 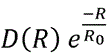, to estimate the correlation length *R_0_*. This analysis revealed a similar characteristic length scale of the correlated flow in human (*R_0_*= 23 ± 8 μm) and rat airways (*R_0_*= 29 ± 7 μm) (**Fig. 2d, left**). We also compared mean particle clearance directionality over multiple cell lengths at *R*=80 μm, which was significantly higher in human compared to rat tissues, indicating straighter transport (**Fig. 2d, right)**. All measurements were completed in tissues from healthy, adult lungs that were washed, mucus-free and submerged in aqueous saline buffer, suggesting that the differences in CPB and clearance directionality were due to species-specific properties of ciliary beat and organization^14^ rather than defective^38^ or immature ciliary beat^16^, or altered mucus properties^39,40^.

**Figure 2:**
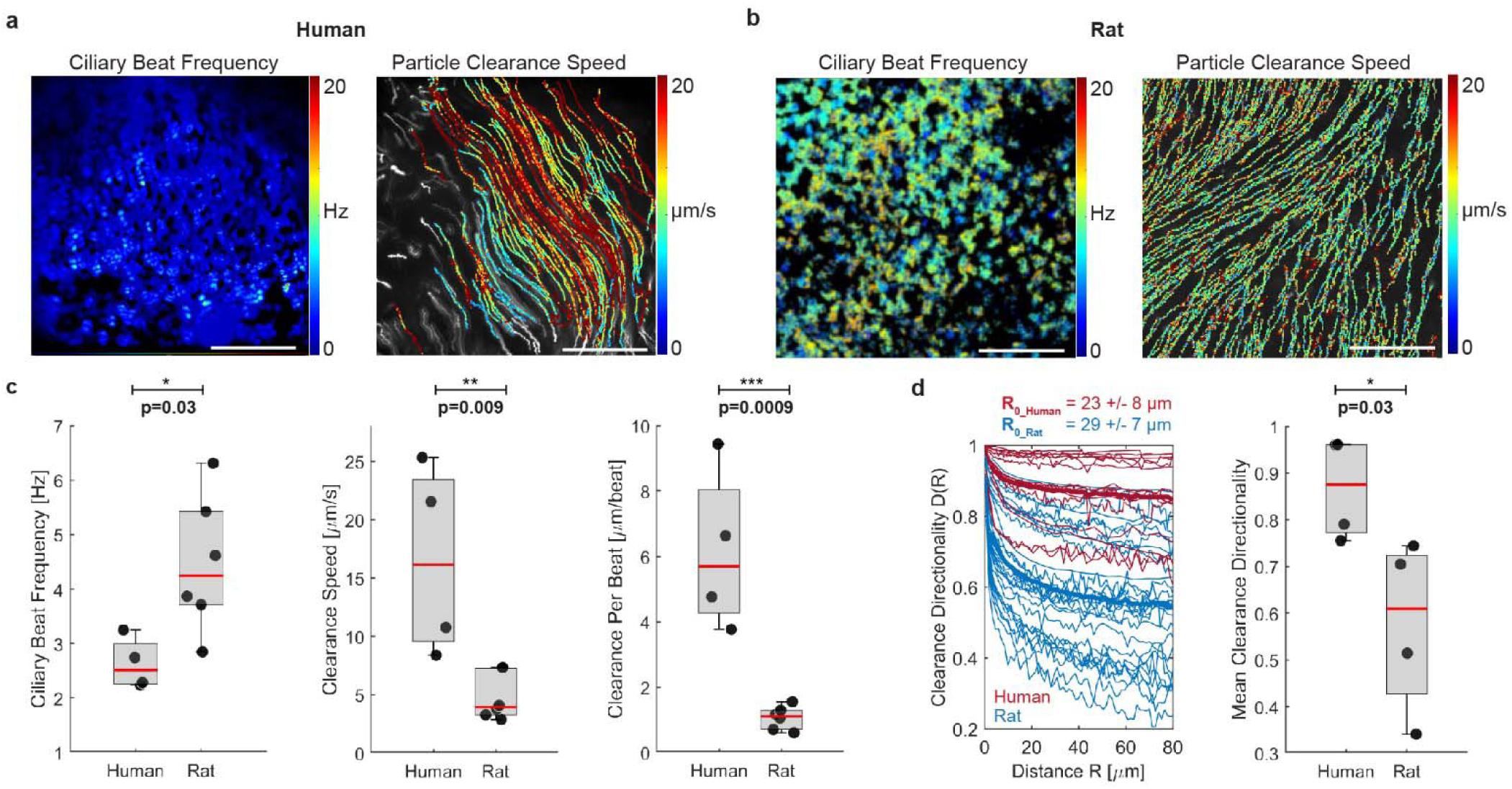
Ciliary beat and particle clearance function differ between human and rat airway. a, Representative measurement of ciliary beat frequency (CBF) and associated particle clearance trajectories and speed in a human airway epithelial sample (BG2). b, Same measurements in rat airway epithelial sample (BG1). Scalebar in a, b: 100 µm. c, Quantification of average CBF, particle clearance speed, and clearance per beat (CPB) in human airways BG0-6 and rat airways BG0-1. Number of human donors: n = 4. Number of rat donors: n=6. d, left: Clearance directionality as a function of distance in human (red) and rat (blue) airways. Thick lines are average curves. Right: Mean directionality over a flow distance of 80 µm. Number of human donors n=4; number of rat donors: n=4. Boxplots: Each solid dot is the mean value of one donor (1-3 BGs, 2-4 FOVs each); red line indicates median, bottom and top edges of the box indicate 25th and 75th percentiles, whiskers indicate minimum and maximum; significance was assessed with two-sided unpaired t-test. Source data for panels c and d are provided as a Source Data file.

To capture these species-to-species differences in ciliary activity, we measured multiple ciliary properties at the single cell and tissue level. On the cellular level, we assessed average ciliary beat orientation, i.e., the angle of the ciliary beat axis, as well as ciliary beat amplitude and cilia length (**Fig. 3a**). At the tissue-level, we analyzed the spatial distribution of ciliated cells using the spatial correlation function *C(R)* (see Methods) to find λ, the average gap distance between ciliated areas^14^ (**Fig. 3b)**. We computed the relative variability of λ using the crystalline order parameter (COP), defined as crystalline 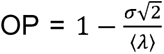 where σ is the standard deviation of λ between field of views^14^. We also determined the degree of alignment of ciliary beat in each field of view using the director-free orientational order parameter defined as ciliary beat 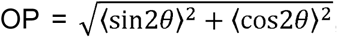, where θ are the measured ciliary beat angles across the field of view. The ciliary beat OP ranges from 0 to 1, where 0 indicates randomly distributed ciliary beat orientations, and 1 indicates spatial alignment of beat.

**Figure 3:**
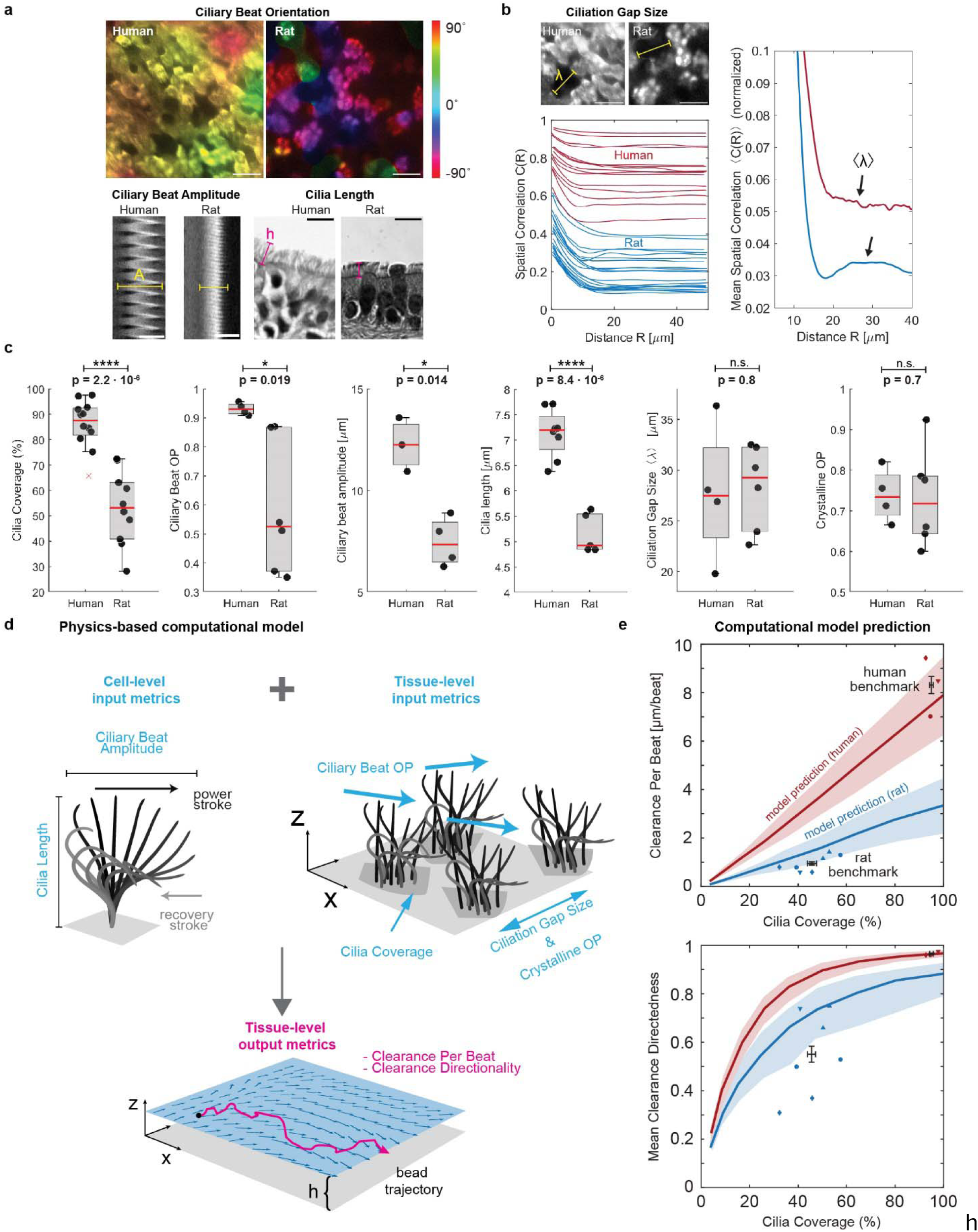
Quantitative analysis of ciliary beat parameters and their impact on clearance function. a, Cell-level analysis of ciliary beat orientation based on beat trajectories (scalebar: 20 µm), ciliary beat amplitude based on kymograph span (scalebar: 10 µm), and cilia length based on histology sections (scalebar: 10 µm). b, Tissue-level analysis of ciliation gap size λ using spatial correlation function *C(R)* of ciliated regions in multiple fields of view (left plot) where the first local maximum of the mean correlation curve reveals mean λ (right plot). Scalebar: 20 µm. c, Average cilia coverage, ciliary beat order, ciliary beat amplitude, cilia length, ciliation gap size and crystalline order parameter measured in human BG0-6 and rat BG0-1. Numbers of human donors from left to right: n= 11, 4, 3, 9, 4, 4. Numbers of rat donors from left to right: n= 9, 4, 4, 5, 7, 6. Boxplots: Each solid dot is the mean value of one donor (1-3 BGs, 2-4 FOVs each); red line indicates median, bottom and top edges of the box indicate 25th and 75th percentiles, whiskers indicate minimum and maximum except outliers, red crosses denote outliers (defined as exceeding +/–2.7 times standard deviation). Significance was assessed with two-sided unpaired t-test. Schematic of ciliary input metrics on cell- and tissue-level used to predict output metrics of tissue-level clearance using physics-based computational model. e, Predicted and measured clearance per beat and clearance directionality in human (red, BG0-6) and rat (blue, BG0-1). Solid line represents mean prediction, shaded area shows uncertainty based on spread of input metrics, and individual data points indicate measurements from human and rat airways. Different marker shapes indicate different donors (n=3 human donors; n=4 rat donors), illustrating presence of two different ciliation levels in trachea of same rat donor (e.g. blue circles). Black data points and error bars represent experimental human and rat benchmark (mean ± SEM of individual data points), indicating a reasonable match of the model predictions. Source data for panels b-d are provided as a Source Data file.

Using these metrics, we compared the human airways at BG0-6 with the rat airways at BG0-1. Shown by the proximal-distal cell type analysis (**Fig. 1d**), coverage with ciliated cells in these regions was significantly higher in the human airways (86.1 ± 9.2%) compared to rat airways (53.1 ± 14.4%) (**Fig. 3c**). We found that cilia length was significantly higher in human compared to rat tissue (human: 7.1 ± 0.5 µm; rat: 5.2 ± 0.4 µm), consistent with literature^16,41^. Ciliary beat OP was also significantly higher in human samples compared to rat samples (human: 0.9 ± 0.02; rat: 0.6 ± 0.2), as was ciliary beat amplitude (human: 12.2 ± 1.3 µm; rat: 7.5 ± 1.2 µm). Ciliation gap sizes were similar in human (λ = 27.8 ± 6.8 µm) and rat airways (λ = 28.3 ± 4.2 µm) and, notably, were thus comparable to the mean correlation length *R_0_* of the cilia-driven flow (23 µm and 29 µm, respectively). This is consistent with prior studies in mice showing that the spatial organization of ciliated cells imprints onto the emergent flow patterns^14^. Crystalline OP was also of similar magnitude between human and rat tissue.

To understand how the functional “output metrics” of clearance effectiveness, namely CPB and clearance directionality, emerge from structural “input metrics,” we developed a simple hydrodynamic model inspired by force singularity models of cilia^42–44^ to simulate particle clearance due to cilia submerged in aqueous liquid, similar to our experimental measurement conditions (**Fig. 3d, Supplementary Figs. 7 & 8, Supplementary Methods**). Here, we model each ciliated cell as a single regularized Stokeslet that scales with cilia beat amplitude, points horizontally in the effective stroke direction, and is located at one cilia length above a no-slip wall that represents the stationary cell surface. The position of ciliated cells and the orientation of the corresponding Stokeslets are chosen such that the simulated epithelium conforms to the desired input metrics such as cilia coverage and ciliary beat OP, patchiness, and crystalline order parameter. Next, the resulting fluid velocity field and tracer particles trajectories were computed to derive CPB and clearance directionality as a function of cilia coverage. To validate the model, we confirmed that it correctly predicted the most dependable clearance measurements in human and rat samples, i.e., from recordings with minimal sample warp where particles could be recorded right above the cilia layer, thereby minimizing distance dependent loss of speed (**Supplementary Figure 9).** In these human recordings, the mean cilia coverage of 95 ± 2.6% was associated with a mean clearance performance of CPB= 8.3 ± 1.2 µm per beat and a mean directionality value of 0.97 ± 0.01 (**Fig. 3e**, “human benchmark”). The model further predicted that CPB is linearly dependent on cilia coverage, while clearance directionality is a steeply rising function that exponentially converges to its maximum above a certain coverage fraction (**Fig. 3e**, red curves). We also computed these curves using the input parameters measured in rat airways. The predicted rat-specific curves (**Fig. 3e**, blue curves) fall below the human-specific curves because of the lower values in ciliary beat OP, amplitude, and cilia length in rats compared to humans (**Fig**. **3c**). This means that for identical cilia coverage, the maximal CPB and clearance directionality are lower in rats. Intriguingly, the experimentally determined mean benchmark value in rat airways with mean cilia coverage of 45.6 ± 8.7%, mean CPB of 0.96 ± 0.3 µm per beat, and mean directionality of 0.55 ± 0.2 (**Fig. 3b**, “rat benchmark”) match the range predicted by the model, suggesting that despite its simplicity, our model captures key structure-function relationships. The details of the human and rat benchmark data are listed in **Supplementary Table S6.**

### *In vitro* airway cultures rarely match human mucociliary properties

We next applied our structural and functional metrics to assess differentiated air-liquid interface (ALI) cultures of primary human airway epithelial cells. ALI cultures are typically used to study human airway diseases and hence there is great interest in establishing human airway-like, aka “organotypic”, phenotypes^45^. We hypothesized that we could generate different luminal epithelial cell type compositions in the same cell donor by using a variety of cell culture differentiation media^46–48^, thereby enabling us to compare these tissues to native human *ex vivo* tissues both in terms of structural organization and clearance function. After expanding the airway epithelial cells of multiple donors (n=4-7; **Supplementary Tables 1 & 2**) in a common medium, we differentiated them for 28 days at ALI in 5 commonly used cell culture media: BD^49^, mAir^47^, SAGM^TM^, PC, or PC-S (see Methods). The tissues differed dramatically in their proportions of luminal ciliated, club, goblet, and hybrid secretory cells depending on culture medium (**Fig. 4a; Supplementary Table 5;** see **Supplementary Figures 10 & 11** for staining in another donor). On average, BD, mAir, and SAGM-cultured tissues exhibited relatively low ciliation and contained a substantial proportion of unidentified luminal cells whereas PC and PC-S cultured tissues most closely resembled the average human *ex vivo* composition measured in BG0-6 (**Fig. 4b**).

**Figure 4:**
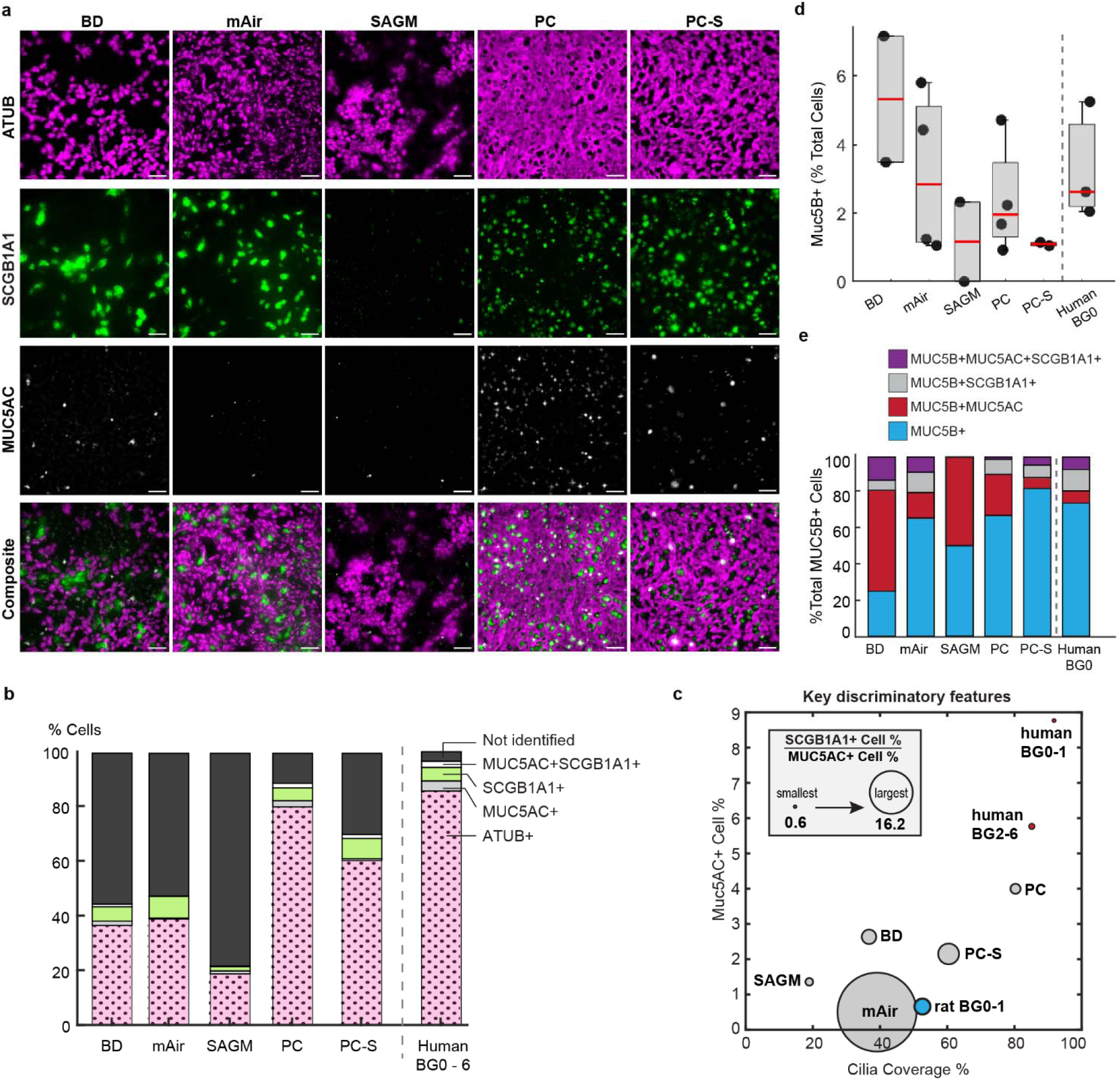
Cell culture media determine *in vitro* luminal cell type composition. a, Representative IF-images of primary human airway epithelial cultures from 1 donor grown in different differentiation media for 28 days at ALI and stained for cilia (ATUB, magenta) and secretory cell markers (SCGB1A1, green; MUC5AC, white). Scalebar, 40 µm. b, Average luminal cell type composition based on IF-staining (*in vitro*: N=4-7 donors, 2 inserts each with 3-8 FOVs each; *ex vivo*: 9 donors, 1-3 BGs, 2-3 FOVs each). c, Mapping of IF-staining data of *in vitro* and *ex vivo* samples onto three dimensions. *y-* axis: percentage of MUC5AC+ cells; x-axis: cilia coverage, i.e., percentage of ciliated (ATUB+) cells; circle diameter: ratio of SCGB1A1+ to MUC5AC+ cell percentages. d, Average percentage of MUC5B expressing cells and e, relative percentage of MUC5B expressing cells that also express either SCGB1A1 or MUC5AC from IF-stainings (*in vitro*: n=2 (SAGM, BD) to n=4 (PC, PC-S, mAir) donors, 1-2 inserts each with 1-6 FOVs each; *ex vivo*: n=3 donors, BG0, 2-4 FOVs each). Boxplot: Each dot represents mean of one donor; red line indicates median, bottom and top edges of the box indicate 25th and 75th percentiles, whiskers indicate minimum and maximum. Source data for panels b-e are provided as a Source Data file.

We next used the cellular composition data to organize the different *in vitro* conditions and visualize their relative similarity to different regions along the human airways tree in a physiologically meaningful way. We reasoned that key organizing metrics would include cilia coverage, which is a major determinant of clearance function (**Fig. 3c)**, percentage of MUC5AC+ cells since the presence of mucin 5AC strongly impacts mucus rheology and flow behavior^39^, and the ratio of SCGB1A1+ to MUC5AC+ cell proportions since this ratio changes along the healthy airway tree (**Fig. 1d)**^25,26^. Here, the definition of SCGB1A1+ and MUC5AC+ cells each includes hybrid MUC5AC+ SCGB1A1+ cells. Mapping the data onto these three dimensions showed that PC-generated cellular compositions were most like human *ex vivo* BG2-6 samples, whereas all other media conditions differed starkly from the human *ex vivo* phenotype (**Fig. 4c**). For further insight into the proximal-distal identity of the *in vitro* cultures, we also analyzed the proportions of MUC5B positive secretory cells, a major cell type present throughout the airway epithelium with highest frequency in the trachea and primary bronchi ^36^. We found that human trachea contained on average 3.3 ± 1.7% Muc5B positive luminal cells, which was matched by similar values across all large airway media conditions (BD: 5.3 ± 2.6%, mAir: 3.1 ± 2.4%, PC: 2.4 ± 1.6%) and lower values in small airway media (SAGM: 1.2 ± 1.6%, PC-S: 1.1 ± 0.1%) which is consistent with a small airway identity^36^. (**Fig. 4d**). As reported previously for both proximal and distal sites^36^, a sizable share of MUC5B positive cells also co-expressed MUC5AC and/or SCGB1A1 both in trachea explants and in all media conditions (**Fig. 4e).**

We proceeded by measuring ciliary beat and clearance function in washed and submerged *in vitro* cultures, to directly compare to the measurements in *ex vivo* samples. Whereas ciliary beat frequency was comparable in all media (**Supplementary Figure S13),** only PC-cultured epithelial tissues approached human *ex vivo*-like particle clearance function with mean CPB of 6.4 ± 2.0 µm s^−1^ and mean directionality of 0.81 ± 0.1 (**Fig. 5a, Supplementary Video 3**). In contrast, particle clearance in tissues differentiated in other media performed well below the human *ex vivo* benchmarks and more closely resembled rat *ex vivo* clearance. To understand the mechanistic underpinnings of these results, we assessed all ciliary input metrics (**Fig. 5b**, **Supplementary Table 4**). Cilia coverage, cilia length, beat amplitude, and ciliation gap size varied the most in response to different differentiation media. On average, BD and SAGM-cultured tissues had shorter cilia than human *ex vivo* samples whereas PC-S cultures had longer cilia, and mAir and PC cultures matched human cilia length (BD: 5.9 ± 0.9 µm; mAir: 6.7 ± 0.8 µm; SAGM: 4.8 ± 1.5 µm ; PC: 7.0 ± 0.4 µm; PC-S: 7.7 ± 0.9 µm; human: 7.1 ± 0.5 µm). Notably, only PC-cultured tissues reached human organotypic ciliary beat amplitudes whereas all other culture conditions fell short (BD: 9.3 ± 1.3 µm; mAir: 8.1 ± 1.4 µm; SAGM: 7.1 ± 0.5 µm; PC: 12.4 ± 1.5 µm; PC-S: 7.2 ± 0.6 µm; human: 12.2 ± 1.3 µm). Compared to human and rat airways, mean ciliation gap size λ was notably smaller in PC and PC-S cultures (BD: 24.5 ± 9.0 µm; mAir: 21.0 ± 6.4 µm; SAGM: 28.1 ± 5.2 µm; PC: 16.6 ± 5.9 µm; PC-S: 14.8 ± 3.7 µm; human: 27.8 ± 6.8 µm), which is consistent with the comparatively small cell size in PC and PC-S cultured epithelia^50^ (**Supplementary Figure 11).** To assess the functional impact of these differences, we entered the ciliary input metrics of each medium condition into our computational model to predict CPB and particle clearance directionality as a function of cilia coverage (**Fig. 5c, top**). Overlaying the measured CPB and directionality values demonstrated a good fit with the model predictions overall. We then replaced individual medium-specific cilia input parameters with the *ex vivo* values and computed the impact on CPB and directionality (**Fig. 5c, bottom**). These results suggest that our ciliary input metrics alone can be used to predict particle clearance function, explain its divergence from organotypic performance and estimate the impact that rescuing defective ciliary beat features would have on clearance.

**Figure 5:**
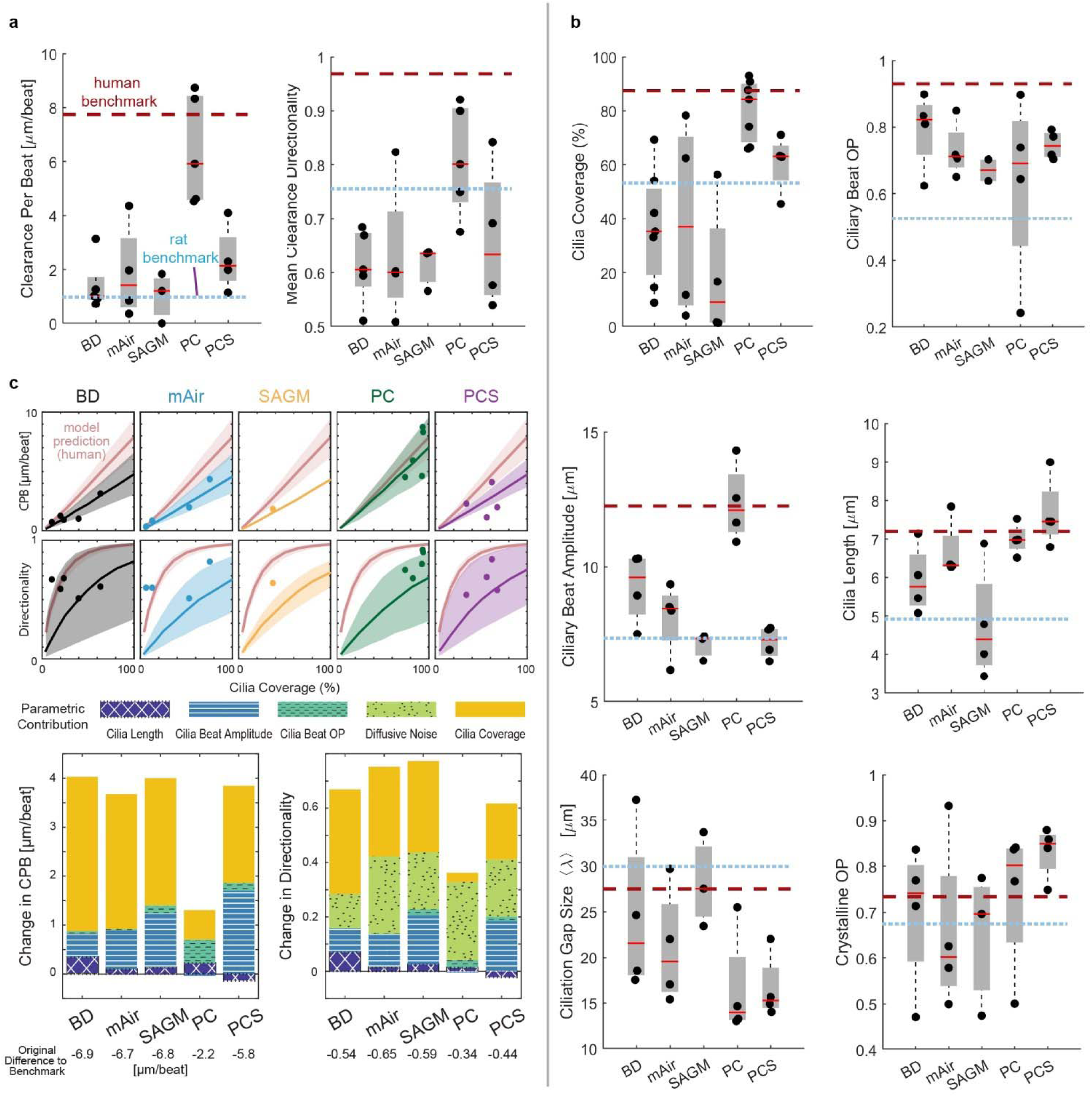
Cell culture media dramatically impact *in vitro* ciliary beat and clearance function. a, Quantitative analysis of particle CPB and directionality in primary human airway epithelial cultures grown in different differentiation media for 28 days at ALI compared to human and rat benchmark data. Number of donors (2-3 inserts each with 6 FOVs each) for BD, mAir, SAGM, PC, PC-S, respectively: n=5, 4, 4, 5, 4. b, Quantitative analysis of ciliary beat metrics in airway cells cultured and visualized as in (a). Number of donors (2-3 inserts each with 6 FOVs each) for BD, mAir, SAGM, PC, PC-S, respectively: Cilia Coverage, n=7, 4, 4, 7, 4; Ciliary Beat OP, n=4, 4, 2 (1 not measurable), 4, 4; Ciliary Beat Amplitude: n=4, 4, 3, 4, 4; Cilia Length, n= 4, 4, 4, 4, 4; Ciliation Gap, n=4, 4, 3, 4, 4; Crystalline OP, n=4, 4, 3, 4, 4. Boxplots in a and b: Each solid dot is the mean value of one donor (1-3 BGs, 2-4 FOVs each); red line indicates median, bottom and top edges of the box indicate 25th and 75th percentiles, whiskers indicate minimum and maximum. Dotted lines indicate average human and rat benchmark values. c, Top: Predicted CPB and clearance directionality in *in vitro* cultures compared to predicted human airway performance (red). Shaded regions indicate uncertainty based on spread of input metrics. Measured mean values per donor are overlaid (dots). Bottom: Predicted change in CPB and directionality of *in vitro* cultures in different media if an individual cilia input parameter is set to match *ex vivo* values. Source data for panel a-c are provided as a Source Data file.

An obvious limitation of our analysis is the washing step prior to recording, thus likely removing most of the mucus, which plays a vital role in trapping particles but also aligning flow^39^, and the underlying periciliary liquid, which has been shown to contain ATP^51^, a potent stimulator of ciliary beat frequency and amplitude^52,53^. To gauge the robustness of our results in physiological mucus conditions, we compared ciliary activity and particle clearance in PC and PC-S cultures with mucus^39^ and without mucus. We found that, in samples with similar cilia coverage, the presence of mucus did not lead to significant changes in average CPB, or directionality compared to washed samples, although beat amplitude was increased in mucus samples (+20.5 ± 14%) (**Supplementary Figure 14A to D**). As a positive control, we dosed the washed samples with 50 µM ATP^54^. After 2 minutes of incubation, ATP induced a slight increase in average CBF (+3.5 ± 6%) and a much greater increase in ciliary beat amplitude (+51 ± 25%), associated with a similar rise in clearance distance per beat (+75.7 ± 50%) compared to control (**Supplementary Figure 14A to D**). Our results match the reported ranges of ATP-mediated changes in CBF, beat amplitude, and CPB in healthy airway epithelial cells^52,53^. We then used our computational model to simulate the impact of the measured changes in ciliary beat amplitude on CPB and directionality, and we compared the predictions to experimentally determined values (**Supplementary Figure 14E and F**). Since the model explicitly accounts for ciliation levels, we were able to increase the data set shown in **Supplementary Figure 14C and D** by including samples varying in cilia coverages. The model curves correctly predicted the similar distribution of washed and mucus-containing samples, while capturing a slight increase in CPB and directionality in mucus samples associated with the higher beat amplitude. The model also recapitulated the notable increase in CPB due to ATP treatment. Taken together, these findings suggest that the presence of an intact mucus layer may indeed increase extracellular ATP levels and hence raise ciliary beat amplitude, thereby slightly increasing CPB and directionality; however, donor-to-donor variability was high, and, on average, there was little difference. Therefore, for modeling average healthy conditions, our structure-function model provides reasonable estimates in both absence and presence of mucus but additional parameters accounting for mucus properties are likely required to predict specific donor responses and disease conditions.

### Structural and functional maps enable phenotypic benchmarking

In *in vitro* airway cultures, the physiological development of directional particle clearance depends on the hydrodynamic interaction of dense ciliation with a mucus layer that is neither too high nor too low in viscosity; the long-range forces transmitted via flowing mucus aligns ciliary beat during differentiation, leading to long-range clearance^55,56^. Mucus viscosity and flowability in turn depend on the proportions and abundance of mucins, especially Muc5AC and Muc5B^9,39^. However, tools to assess this relationship remain limited, especially given the difficulty of measuring mucus rheology in miniscule *in vitro* samples^57^. We therefore evaluated whether ciliated and secretory cell type composition could be directly predictive of average clearance directionality in the human BG0-6 samples and the *in vitro* cultures. Indeed, a simple linear regression model resulted in a solid prediction of mean clearance directionality in human *in vitro* cultures using as inputs the percentage of ciliated cells and secretory cells (i.e. the sum of MUC5AC, SCGB1A1 and/or MUC5B positive cells) (**Fig. 6**, coefficient of determination R^2^=0.89). Other input combinations worsened the prediction, including using the percentage of ciliated cells as sole input, or removing it (R^2^=0.77 and R^2^=-1.34, respectively), or replacing the secretory cell percentage with only MUC5AC, MUC5B, or SCGB1A1 positive cell percentages (R^2^=0.56, R^2^=0.8, R^2^ =0.55), respectively). Hence, combining information on both ciliated and major secretory cell proportions was most predictive for clearance directionality.

**Figure 6:**
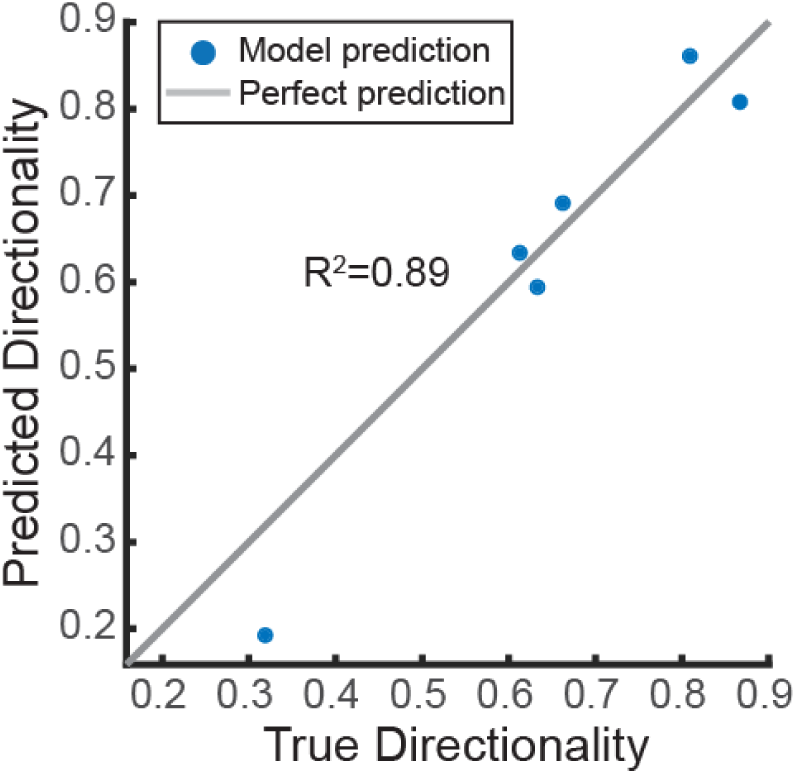
Clearance prediction from cell type composition. Linear regression model predicting average clearance directionality in human airway epithelia (*in vitro* and *ex vivo*) using as input the average values of cilia coverage and secretory cell percentage (incl. SCGB1A1+ , MUC5AC+ , and MUC5B+ cells). Source data are provided as Source Data file.

Collectively, our analysis has uncovered the following key findings: (1) structural ciliary metrics, particularly the extent of ciliated cell coverage, are mechanistic predictors of particle clearance function; (2) the selection of cell culture medium significantly influences ciliary metrics and consequently particle clearance function, and (3) the composition of luminal secretory and ciliated cell types may act as a statistical predictor of particle clearance directionality in airway epithelia.

Finally, to demonstrate the broader applicability of our analysis for tissue phenotyping, we leveraged the structural cell-type composition map (**Fig. 4c**) and the functional CPB-against-ciliation map (**Fig. 5c**) to assess the mucociliary machinery in additional culture conditions and animal models, where the data was derived from published literature or via proof-of-concept experiments (**Fig. 7a-b**). The systems evaluated included mature and developing mouse trachea, human airway epithelial cultures derived from human induced pluripotent stem cells, and primary human airway epithelial cultures subjected to asthma-like inflammatory conditions (interleukin-13 (IL-13) treated) or differentiated under mechanical stimulation (Organ-on-Chip models). We found that the pluripotent stem cells-derived airway epithelial tissue differentiated in PC at ALI (iALIs) as described earlier^58^ greatly increased cilia coverage between day 14 and day 35 at ALI; however, CPB remained much lower than expected from the extent of cilia coverage, suggesting immature ciliated cell function and organization (**Fig. 7a**, orange triangles). Secretory cell type composition in iALIs was dominated by SCGB1A1+ cells and showed relatively low levels of MUC5AC+ cells (**Fig. 7b**). The CPB of primary human airway epithelial cultures differentiated in BD was also below the human benchmark curve at day 14 of ALI when cultured in conventional static inserts (**Fig. 7a**, unfilled square) but approached the curve when cultured in continuously perfused Organ-on-Chips (**Fig. 7a**, inverted triangle). Further, after a prolonged culture time of 35 days, the static insert cultures increased ciliation and reached the human CPB benchmark curve, indicating maturation of ciliary beat (**Fig. 7a**, solid square). Luminal cell type composition in both insert and chip conditions reflected a lack of ciliation compared to the human benchmark; however, the Organ-on-Chip cultures reached nearly organotypic proportions of MUC5AC+ cells^49^ (**Fig. 7b**). The CPB of primary human airway epithelial cultures cultured in Vertex ALI medium, a popular medium choice for modeling cystic fibrosis and asthma due to the high proportion of secretory cells^59^, lay below the benchmark curve at day 35 of ALI. Cultures treated with IL-13 for the final 14 days lost cilia coverage and, despite ciliary beat, generated almost no flow at all (**Fig. 7a**, stars). The structural map reflects the high proportion of secretory cells expected from culture in Vertex ALI medium^59^, and IL-13 treatment creates the expected goblet-cell dominated Th2-like asthmatic phenotype^60^ (**Fig. 7b**). Ciliation and CPB in mouse trachea continue to increase after birth until, at approximately postnatal day (P) 15, they reach their mature performance at 40-45% ciliation^14,61^ (**Fig. 7a**, unfilled circle). The ciliation-dependent increase in CPB parallels the observed and predicted trends in the rat trachea-bronchial airways (**Fig. 7b**, filled star and blue model prediction), and hence it is possible that mouse and rat trachea share similar ciliary beat properties.

**Figure 7:**
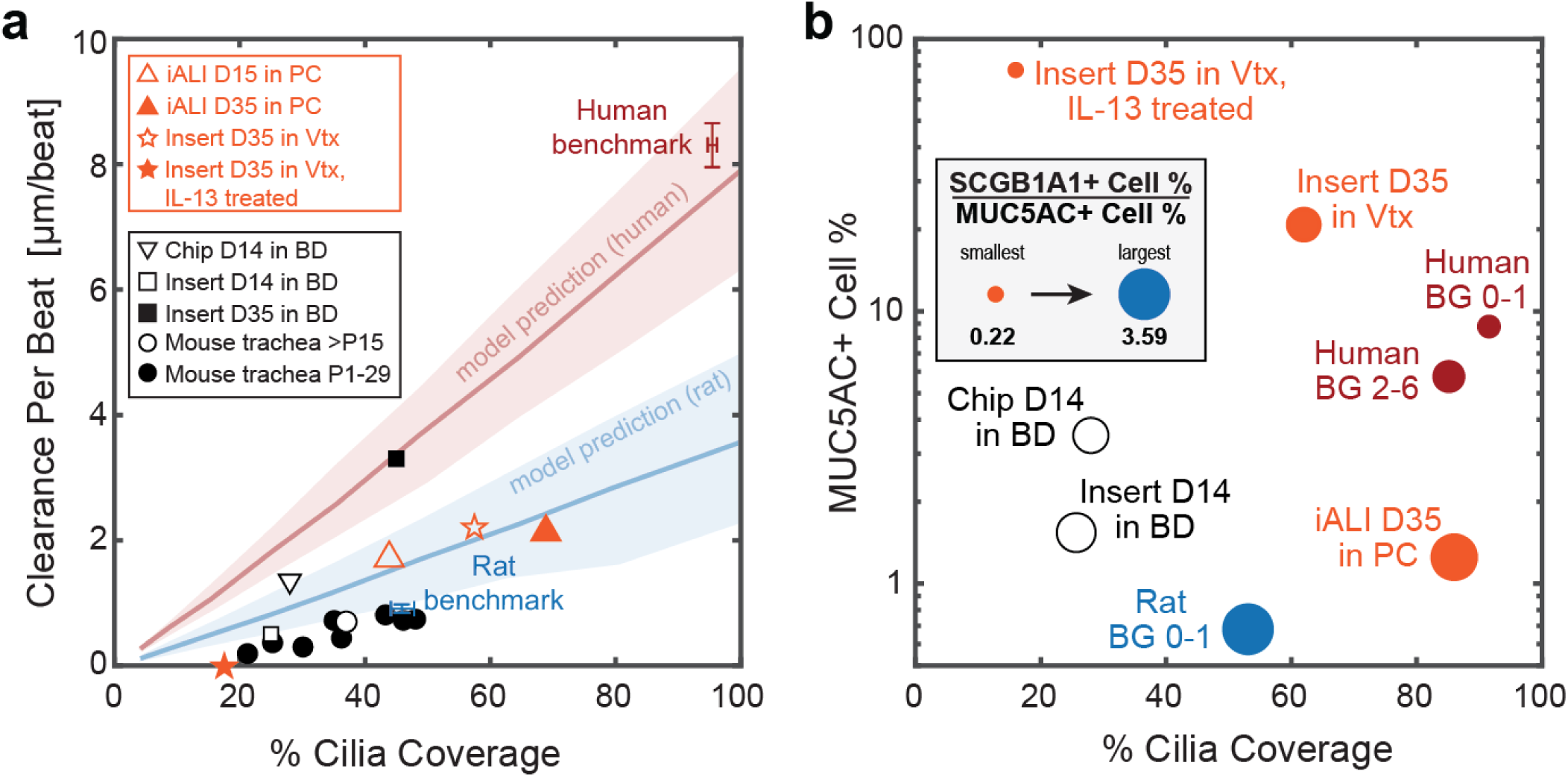
Structural and functional benchmarking of the mucociliary machinery. a, Clearance per beat map comparing different human in vitro and rodent ex vivo models to human and rat benchmark data. D, day at ALI; P, postnatal day; Vtx, Vertex ALI medium; iALI, human iPSC-derived differentiated airway epithelium. Orange markers indicate original data (iALI, Vtx and Vtx +IL13, proof-of-concept from N=1 donor each, 2-3 inserts, 3-6 FOVs each); black markers indicate data sourced from literature, see methods for details. Red line and shaded region: Human model predictions; blue line and shaded region: rat BG0-1 model predictions. Shaded regions indicate uncertainty based on spread of input metrics. b, Cellular composition map comparing different human *in vitro* models to human and rat benchmarks with *y-*axis: percentage of MUC5AC+ cells; *x-*axis: cilia coverage, i.e., percentage of ciliated (ATUB+) cells; circle diameter: ratio of SCGB1A1+ to MUC5AC+ cell percentages. Solid markers indicate original data (human BGs as Fig. 4.; iALI, Vtx and Vtx +IL13, proof-of-concept from N=1 donor each, 2-3 inserts, 3-6 FOVs each); black unfilled markers indicate data sourced from literature. Note that *y-*axis is in log-scale. Source data for panels a and b are provided as a Source Data file.

## DISUSSION

We developed a framework to predict how airway morphology influences particle clearance effectiveness and established benchmarks for comparing experimental airway models against the gold standard, the human airways. Specifically, we discovered that the human large airways exhibit a high degree of ciliation, akin to dogs and pigs^62,63^, while rat airways display a gradual increase starting from 45% ciliation in the trachea. These findings challenge the notion of conserved cilia coverage and clearance function in mouse and human airways^14^. Furthermore, mouse airways diverge from human airways in other relevant aspects, including near 100-fold smaller spatial dimensions, a different branching morphology, and the absence of mucus-producing goblet cells in healthy conditions^64^, similar to our findings in rats. Additionally, it was previously found that deposition of inhaled particle deposition differs between humans and obligatory nose breathers like rats and mice^65,66^, suggesting that in contrast to humans, in rats and mice the nasal respiratory tract serves as the main particle filter instead of the tracheobronchial tree. Indeed, a recent study showed that nasal ciliation in mice and associated CPB is multiple times higher than in the trachea and comparable to the values we measured in the human large airways^67^. However, the functional implications remain speculative, necessitating further investigation and comparative studies especially on the role of mucus.

Our study is first to evaluate culture conditions based on their ability to replicate human-typic characteristics of MCC. We developed structural and functional maps that leverage markers commonly measured, enabling us to visually compare different experimental models with human organotypic benchmarks. While previous studies have assessed the impact of cell culture media on *in vitro* differentiation, morphology and functional responses of respiratory epithelia^47,68,69^, they did not directly compare these metrics to native human airways and hence lacked organotypic benchmarks. Intriguingly, we demonstrated that the composition of secretory cell types may predict clearance directionality, consistent with the need for mucus during differentiation to establish long-range clearance^55^. This relationship provides impetus for future work on mucus micro-rheology and cilia-mucus interactions.

Our results suggest that commonly used culture media and protocols are insufficient to recreate the mucociliary performance of healthy airway epithelia in humans. We found that one of the major causes of diminished clearance *in vitro* is simply insufficient cilia coverage. Suppression of Notch signaling, which controls the balance between ciliated and secretory cells^70^, has been shown to increase ciliation *in vitro*^71^. Additionally, we found that many *in vitro* conditions were deficient in ciliary beat properties.

There are several potential strategies for improvement, such as stimulating Wnt signaling. Canonical Wnt signaling is essential for proliferation, migration and many other processes in airway epithelial development^72^, including specifically cilia biogenesis and beating^73^, whereas non-canonical Wnt signalling establishes planar cell polarity (PCP) and coordinated ciliary beat^74^, which is required for effective and directional clearance. Other possibilities include stimulation of nitric oxide generation, as nitric oxide also drives PCP signaling and furthermore increases ciliary beat frequency^75^. Faster beat in turn promotes cilia-driven flow and associated shear forces, which are known to stimulate alignment of ciliary beat and clearance^55^. Indeed, directly applying fluid shear forces using Organ-Chip devices accelerated mucociliary differentiation and polarized clearance^49,76^, indicating that targeting mechanosensory signaling might also be a powerful strategy for improving mucociliary clearance *in vitro*.

Our study does have limitations. The analysis of ciliary function and clearance in rat and human airways was performed on explanted tissue and not *in vivo*, and the possibility of artifacts must be considered in the interpretation of our findings. Analysis was also limited by the availability of healthy human airway samples (15 donors total, as detailed in **Supplementary Table 1**), some of which were extracted from peritumoral tissue. Many of the human donors were above the age of 60, raising the possibility of aging-related changes to MCC^77^. Despite these limitations, the robustly high ciliation levels provide confidence that this is a key characteristic of human large airways. While more studies are needed to also characterize MCC in the small airways, 85% luminal cilia coverage was reported in the human small airways^78^, suggesting continuously high ciliation levels throughout the conducting airways. We washed all samples prior to live imaging to enable direct comparison between buffer-stored explants and *in vitro* cultures, thereby lacking an intact periciliary liquid and mucus layer in our analysis. However, MCC recordings in unwashed mice trachea^79^ suggest a CPB of about 1-2 µm per beat (assuming a physiological CBF of 15 Hz^61^), comparable to the CPB measured in washed mouse trachea^14^. We also showed that our clearance metrics resulted in comparable average values between washed and unwashed conditions, but the high donor-to-donor variability indicates that further studies are warranted. We propose that in the case of airway disease, where mucus properties are often abnormal, comparing our metrics with and without mucus will help dissect the contribution of secretory compared to ciliary dysfunction. Other technical limitations are outlined in the **Supplementary methods.**

In conclusion, our comprehensive structure-function analysis of the mucociliary machinery along the human airway tree provides quantitative benchmarks and visualization tools to assess how human organotypic the mucociliary barrier of experimental models are. The analysis can also reveal the effects of maturation, treatments, and diseases on ciliary activity and resulting MCC. Our physics-based model linking cilia properties to particle clearance can aid in estimating the efficacy of treatments aimed at restoring MCC in humans.

## METHODS

### Ethical regulations

Our research complies with all relevant ethical regulations for the procurement and use of animal and human cells and tissues were followed. The details of these guidelines and specific protocols are listed in the next section on cell and tissue sourcing.

### Cell and tissue sourcing

#### Source of rat airways

Rat lungs were harvested from control animals that were freshly euthanized as part of other studies performed according to approved animal study protocols. The animals investigated at USC were 3-8 months-old female Wistar-Kyoto rats sourced from Charles River Laboratories, MA, USA, that were handled and euthanized (urethane injection) according to the approved Animal Study Protocols (IACUC 20263 and IACUC 20751). The animals investigated at Helmholtz/TUM were 2 months-old female Wistar rats from Charles River Laboratories, MA, USA, that were handled and euthanized (pentobabital injection) according to the approved Animal Study Protocol (TVA 02-17-177). Only female rats were used due to the availability of fresh tissues donated from other studies. Also see **Supplementary Table 1**.

#### Source of whole human lung, tracheobronchial rings, and cells at USC and University of Iowa

Human lung tissue from subjects with no prior history of chronic lung disease was obtained through the International Institute for the Advancement of Medicine (IIAM) with approval from the Institutional Review Board (IRB) of the University of Southern California (USC) (Protocol number: #HS-18-00273) and from the Center for Gene Therapy’s Cells and Tissue Core facility at the University of Iowa (tissues are obtained under IRB#199507432) and deidentified samples were provided to Dr. Ryan’s laboratory. Donor demographics are included in **Supplementary Table 1**. Human trachea-bronchial epithelial cells (HTBECs) were isolated^80^. Briefly, proximal airways including the trachea, main stem bronchi and 2 further branching generations of the cartilaginous airways were dissected into 1-4 cm^2^ sections and digested in of 0.1% Protease XIV (Sigma #P5147) and 0.001% DNase (Sigma #DN25) (%w/v) in DMEM/F12 (ThermoFisher #11330032) overnight at 4°C. Using a scalpel, epithelial tissue was gently scraped, collected in DMEM/F12, and centrifuged at 400 x g for 5 minutes. After red blood cell lysis in ACK lysis buffer (ThermoFisher #A1049201), epithelial cells were single cell dissociated by incubation with Accutase (Innovative Cell Technologies #AT104) at 37°C for 30 minutes. Cells were then seeded at a density of 30K cells cm^−2^ on PureCol (Advanced Biomatrix #5005) coated tissue culture dishes in airway epithelial growth media (Promocell #C-21160) and passaged at 80% confluence. At each passage cells were seeded at 5×10^3^ cells cm^−2^.

#### Source of human bronchial rings at LUMC

Bronchial rings were dissected from macroscopically normal lung tissue obtained from patients undergoing resection surgery for lung cancer at the Leiden University Medical Center, the Netherlands (**Supplementary Table S1**). Patients from which this lung tissue was derived were enrolled in the biobank via a no-objection system for coded anonymous further use of such tissue (www.coreon.org), not requiring written consent. All patient material used in this study was de-identified and approved for research use by the institutional medical ethical committee (BB22.006/AB/ab). No clinical data from the patients from which the tissues were derived for this study are available.

#### Source of cells at TUM/Helmholtz

Cells were sourced from commercial suppliers (**Supplementary Table 1**).

#### Source of iPSC-derived cultures

See section “Data for comparative CPB and cell type composition map.”

#### Source and generation of human primary airway epithelial cell cultures

Human primary small airway epithelial cells (hSAECs) and trachea-bronchial cells (hTBECs) were obtained from Lifeline Cell Technologies (USA) or via isolation from primary tissue obtained through the International Institute for the Advancement of Medicine (IIAM) with approval from the Institutional Review Board (IRB) of the University of Southern California (USC) (Protocol number: #HS-18-00273). (**Supplementary Table 1**). The first passage cells were expanded in collagen I coated T75 tissue culture flasks and dishes in standard bronchial epithelial cell medium (BEpiCM) (ScienCell) until ∼90% confluency. Expanded small airway and bronchial/tracheal cells were seeded on collagen IV (300 µg mL^−1^) coated 12-well 0.4 pore diameter PET Transwell membranes (Corning, 3460) at a density of 150K cells per insert (∼135K cells cm^−^²). The cells from each donor were cultured in BEpiCM supplemented with 1 nM EC23 (Tocris Bioscience) until confluent. Once the tissue was confluent, differentiation was induced by introducing air liquid interface (ALI) via removal of the apical medium (day 0 of ALI culture) and using one of five different differentiation medium in the basal compartment: 1. BD: bronchial epithelial base medium and Dulbecco’s modified eagle medium BEpiCM:DMEM (50:50) with addition of supplements and 50 nM EC23^49^; 2. PC: PneumaCult ALI (STEMCELL Technologies); 3. PC-S: PneumaCult ALI-S (STEMCELL Technologies); 4. SAGM: Small Airway Epithelial Growth Medium (Promocell, C-21170) containing 50 nM EC23; and 5. mAir: a 1:1 mix of Dulbecco’s modified eagle medium and Airway epithelial cell growth medium (AECGM, PromoCell, C-21160) with AECGM supplements and 50 nM EC23 (previously described in ^47^.) The apical surface was washed with phosphate-buffered saline (PBS, no Calcium and Magnesium) at 37 degrees Celsius twice a week to remove excess mucus. For bead transport measurements in samples containing mucus, 50 µL cm^−^² of the respective media was added to the apical surface every other day during differentiation to prevent mucus dehydration^55^. Cultures were differentiated until day 28 of ALI.

### Live ciliary beat and particle clearance recordings

#### Ex vivo samples

We freshly isolated airway epithelial tissue from respiratory tree branching generation (BG) 0 through BG6 from healthy transplant-rejected human whole lungs, and BG0 through BG5 from healthy rat whole lungs for live measurements. Samples were cut open along the airway tube to reveal the luminal epithelium, submerged in ice-cold HBSS buffer, mounted upside down in a glass bottom dish (ibidi), and gently flattened using a glass cover slip held down with silicone grease at the corners. All live recordings were performed at room temperature, which we found to prolong sample viability compared to higher temperatures. In some samples, ciliary beat was recorded with phase contrast at 100-200 frames per second (fps) using an inverted Leica microscope equipped with a 40x (NA 0.8) objective and a PCO Edge 4.2 high speed camera (Excelitas Technologies) operated with the micromanager plugin (ImageJ), or a Leica K3M camera operated with Leica software LAS-X. Cilia were also live-stained with fluorescent-dye conjugated wheat germ agglutinin (ThermoFisher; 20 minute incubation in dilution of 1:200)^15^, and 1-µm fluorescent tracer particles were added to the bath, such that live ciliary beat and particle clearance could be recorded in the same field of view for 10 seconds (ciliary beat: 15-33 fps; particle trajectories 8-33 fps) using epifluorescence imaging^49^. Two to four FOVs with visible CBF were recorded from each sample.

#### In vitro cultures

*In vitro* cultures were washed for 10 minutes with PBS and recorded at ALI using an inverted Zeiss microscope equipped with a 40x (NA 0.8) phase contrast objective and a temperature-controlled chamber that was preheated to 37°C (in contrast to the *ex vivo* recordings, see above). Movies of ciliary beat were taken at 140 fps using an Orca Flash 4.0 camera (Hamamatsu). To reveal beat kinematics of thicker cultures, samples were mounted upside down and cilia were live-stained with fluorescent-dye conjugated tomato lectin (IVISense™ Tomato Lectin 680, Perkin Elmer) by incubating the sample in a 0.25 µM dilution in PBS for 20 minutes. After rinsing, ciliary beat kinematics were recorded at 30 fps using epifluorescence imaging. Particle clearance was recorded for 10 seconds at 20-30 fps by adding 1-µm fluorescent tracer particles to the apical surface^49^. Video recordings were taken from 2 insert cultures per donor and condition, with at least 8 FOVs per sample.

To assess ciliary beat and clearance function in the presence of mucus, 50 µL cm^−^² of medium containing carboxylate-modified 1-µm fluorescent tracer particles (1:1000 dilution) were added to the PBS-washed apical surface two days prior to the recording, allowing mucus to accumulate and embed the beads in mucus. At day 30 of differentiation, ciliary beat activity and clearance were measured as described above (protocol based on Song *et al* (2022)^39^).

To test the impact of extracellular ATP on ciliary beat and resulting transport, 50 µL cm^−^² of medium containing 50 µM ATP plus 1-µm fluorescent tracer particles (1:1000 dilution) were added to the apical surface post PBS-wash. After 2 minutes of incubation, ciliary beat activity and particle clearance were recorded (protocol based on Lieb *et al* (2022)^54^https://doi.org/10.1113/jphysiol.2001.013222).

### Ciliary input metrics

#### Cilia length in ex vivo samples

Hematoxylin & eosin (H&E)-stained sections of human bronchial rings and rat airway trachea were imaged using a Zeiss Axioscope 7 fluorescence microscope and a 40x oil DIC objective (NA 1.4). Cilia length data were measured manually using the freehand line tool in the image processing software Fiji ImageJ (Version 1.54m)^81^. Only cilia with visible starting and end point were measured. For each rat donor, on average 15 FOVs were analyzed and on average 10 cilia were measured in each FOV. For each human donor, on average 4 FOVs were analyzed and on average 30 cilia were measured in each FOV.

#### Cilia length in in vitro samples

The cell layer was dissociated by incubating the cultures for 20 min in prewarmed Accutase (Invitrogen, 00-4555-56) in a conical tube. After 20 min warm culture medium was added at a ratio of 3:1 and the cells were centrifuged at 210× g for 7 min. The resulting pellet was resuspended in 500 µL 4% paraformaldehyde and 10 µL of this suspension was placed onto a glass bottom imaging dish (ibidi,81218-200) and covered with a glass coverslip. Cilia length was measured as above. For each donor, 30 FOVs were analyzed and on average 10 cilia were measured in each FOV.

#### Ciliary beat frequency (CBF)

CBF was measured by applying Fourier spectral analysis to each cilia-associated pixel recorded in high-speed videos^22^.

#### Ciliary beat order

We determined ciliary beat orientations using either the ImageJ plugin directional analysis or manual tracing of the beat axis from all ciliated cells in at least 3 FOVs, each spanning approximately 200 µm by 200 µm. This analysis yields a list of beat angles across each field of view. We derived the director-free ciliary beat order parameter (OP) for each FOV from the angle distribution as follows: 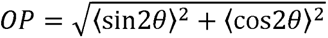, where 〈〉 indicates the mean and θ indicates the measured angles. To compute the average value per donor and condition, we averaged the ciliary beat OP over all FOVs.

#### Ciliary beat amplitude

We measured ciliary beat amplitude by manually tracing the span of the ciliary beat using kymographs of videorecordings^82^ in at least 10 ciliated cells each in 3 FOVs.

#### Cilia coverage, ciliation gap size, and crystalline order

These metrics were measured from the standard deviation of pixel intensity across all video frames, revealing the spatial distribution of motile cilia. When these data were unavailable, cilia coverage revealed by IF-staining was used instead. Images were thresholded and binarized to reveal the ciliation pattern, and ciliation gap size and crystalline order were estimated as shown previously^14^. Briefly, the 2-point correlation function *C(R)* was used to measure the probability that two pixels at a certain distance from each other both have a binary intensity level of “1”, i.e., are part of a ciliated cell. For an image with pixel dimensions *m* × *n*, the 2-point correlation function is defined as

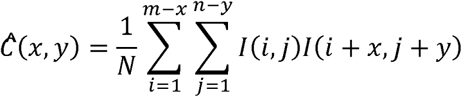

where *N=(m-x)(n-y).* From this function, *C(R)* is derived by only allowing for coordinate pairs (*x,y*) that are part of a circle perimeter of radius *R* (rounded to integer pixel coordinates) and then averaging over the number of pixels in the circle perimeter. The resulting function *C(R)* is oscillating and its first local maximum reveals λ, the average spacing of two ciliated patches^14^. The crystalline order parameter (COP) describes the degree of variability of the patchiness between multiple FOVs and is derived from the average of λ and its standard deviation std(λ) across FOVs as follows: 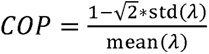. Cilia coverage, defined here as percentage of luminal cells that are ciliated, was determined as part of the cell type composition analysis discussed in the associated methods section.

### Particle clearance output metrics

The displacement and trajectories of fluorescent tracers driven by ciliary beat was automatically measured using the open source ImageJ Trackmate plugin^83^. From these data, two metrics were calculated.

#### Particle clearance directionality

Particle clearance directionality *D(R)* was defined in the Eulerian framework. We derived a Eulerian vector field from the particle trajectories by averaging the velocity components *(u,v)* at each image coordinate *(x,y)* over all trajectories passing through *(x,y)* at any time during the recorded video, i.e., 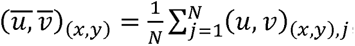, where *N* is the total number of trajectories passing through *(x,y)*. This procedure creates a temporally averaged flow vector field, which is useful when tracer particles density is low at any given moment in time. *D(R)* was defined as the magnitude of with side length *R*, i.e., *D*(*R*) = |<v)|<|v|), where 〈〉 indicates the mean and || indicates the magnitude of the average flow vector divided by the average magnitude of all flow vectors within a square window each flow vector *v*. After normalization to mitigate offset, each trace *D(R)* was fitted with a decaying exponential, 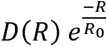, to reveal the correlation length *R*_0_. To remove ill-fitted curves, we only accepted fits with coefficient of determination above 0.8. We used bootstrapping to estimate the statistics of *R*_0_ and account for the limited sample size. Since *D(R)* decays with window size *R*, we defined its mean value <*D_R_*=80>by averaging *D(R)* at *R*=80 µm to assess transport directionality over multiple cell length.

#### Clearance distance per beat

Clearance per beat (CPB) was defined as the mean speed of particle clearance (in units of µm per second) divided by the ciliary beat frequency (beats per second), resulting in units of µm per beat. This is a measure of the efficacy of each ciliary beat cycle in driving particle clearance. When both speed and CBF data were available for the same FOV, CPB was computed directly from the ratio. When speed and CBF data were recorded separately, their mean values over multiple fields of views were used to compute one single value of CPB from their ratios.

In *ex vivo* samples, the surface topography was often distorted due to elastic recoil after cutting the cartilage rings, leading to contorted tracer trajectories and variable distance between tracer particle and cilia, which impacts apparent clearance speeds (**Supplementary Fig. 6**). To establish benchmarks and to validate the physics-based model, we used the clearance measurements from human and rat samples that we trusted the most, i.e., measurements from entirely flat airway sections where the particles were visibly touching the cilia. In humans, benchmarks were derived from 3 donors, one recording each, in H44 (BG6), H47 (BG2), and H2924 (BG0). In rats, benchmarks were derived from 4 animals (1 or 2 recordings each), in R43USC (BG0 and BG1), R42USC (BG0), TUMR56 (BG0) and TUMR55 (BG0). **Supplementary Videos 1 & 2** show one example per species.

### IF staining and imaging

#### Human and rat airway sections

Airway rings were fixed using a 4% paraformaldehyde (PFA) solution for 3 to 24h depending on tissue thickness, washed with PBS, and stored in PBS at 4 °C until staining. Prior to staining, the diameters of the rings were measured using a ruler. For staining, sections were cut from both dorsal and ventral sides of the airway rings, to capture potential spatial variability along the perimeter. Nearly level sections were cut along the long axis of the tube to minimize warping of the epithelium. Samples were placed into a 96-well plate for staining.

#### Human airway epithelial cell cultures

After differentiation at ALI, the primary human airway epithelial cultures and iPSC-derived cultures were fixed using incubation with 4% PFA solution for 30 min at RT, then washed again three times with PBS and stored in PBS at 4°C until staining.

#### IF staining

Samples were blocked and permeabilized using 0.25% (v/v) Triton-X 100 in PBS with 3% BSA for 60 min at RT, then incubated overnight at 4°C with primary antibodies (**Supplementary Table S8**) diluted in the Triton/BSA buffer The samples were rinsed three times for 5 min with PBS before incubation with secondary antibodies diluted in Triton/BSA buffer for 1 h at 37 °C, followed by a triple 5 min wash with PBS. Then, as applicable, the samples were incubated with directly conjugated primary antibodies (**Supplementary Table S8**) and F-actin stain phalloidin 555 (Invitrogen, A30106) or phalloidin 405 (Invitrogen, A30104) in Triton/BSA buffer for 1 hour at 37 °C, followed by a triple 5 min wash with PBS. The samples were stored at 4 °C until mounting. For mounting, *ex vivo* samples were placed into glass-bottom imaging dishes (ibidi, 81218-200) and covered with SlowFade™ Glass Antifade Mountant (Invitrogen, S36917). A round coverslip lined with silicone grease was used to push down and flatten the sections. The *in vitro* samples were mounted by removing the cell culture membrane from the insert using a scalpel and placing the membrane onto a glass slide with the cells facing upwards. The membranes were coated with a drop of ProLong™ Diamond Antifade Mountant (Invitrogen, P36965) and covered with a round number 1.5 glass coverslip.

#### IF imaging

Human bronchial ring sections were imaged with a 40× water objective (NA 0.80) using a Leica DMi8 microscope equipped with an Andor Dragonfly 200 spinning disk confocal or using a Zeiss LSM 700 confocal microscope. From every stained ring section, 3 to 8 FOVs with a size of 2048 x 2048 pixels at 6.6 pixels per µm resolution were recorded. Rat samples, other human airway sections, and *in vitro* cultures were imaged using a Leica confocal scanning microscope or a Zeiss Axioscope 7 fluorescence microscope equipped with a 40x oil DIC objective (NA 1.4). Six FOVs with a size of 2048 x 2048 pixels at 6-7 pixels per µm resolution were recorded per sample.

### Cell type composition literature survey

We reviewed human data from standard histology^25,34,35^ and RNAseq studies^32,33^, as well as rat data from standard histology^25^ (**Supplementary Fig. 4**).

### Cell type composition analysis

We employed semi-automated image analysis to quantify luminal cell type composition from IF images containing 4 channels, either labeling F-Actin, MUC5AC, SCGB1A1 and acetylated α-tubulin (ATUB), or F-actin, MUC5AC, MUC5B and SCGB1A1 (**Supplementary Fig. S2**). Using ImageJ Fiji^81^, raw image data was converted to 16 or 8-bit tiff images using maximum projection for stacks. As needed, images were cropped to remove edge artifacts and the subtract background function was applied with a rolling ball radius of 1000 pixels. The Fiji plugin Advanced Trainable Weka Segmentation with all default and the Laplacian, Derivatives and Structure training features^84^ was used to segment cell outlines from the F-actin mesh. The channels for the different cell markers (MUC5AC, SCGB1A1, ATUB and/or MUC5B, dependent on the staining combination) were filtered and optimized in contrast. After these preprocessing steps, cell type composition analysis was performed by overlaying the markers with the cellular outlines using CellProfiler™(Versions 4.0.7 to 4.2.5)^85^, providing total cell number and proportions of ATUB+, MUC5AC+, SCGB1A+, MUC5AC+, and/or MUC5B+ cells, dependent on the staining combination, as well as overlaps of secretory markers, indicating double- or triple positive cells. Image sets with low signal-to-noise ratio were excluded from the analysis.

### Airway histology and imaging

#### Rat trachea

Tracheas were obtained from wildtype Wistar rats and fixed in 2% PFA at 4°C overnight on an orbital shaker. After washing, samples were embedded into paraffin wax in a lateral (dorsoventral) orientation. Traditional H&E staining was performed and microscopical images at 400x magnification were taken of all sections, with at least 6 FOVs per trachea.

#### Human bronchial rings

Bronchial rings were obtained from tumor-free resected lung tissue at the Leiden University Medical Center. The rings were fixed using 4% PFA solution for 24 hours after which the rings were transferred to PBS and stored at 4°C until paraffin embedding at the Department of Pathology at the Leiden University Medical Center. Bronchial ring sections were deparaffinized in xylene and dehydrated in an ethanol gradient. Traditional H&E staining was performed and microscopical images at 400x magnification were taken of all sections, with at least 3 FOVs per sample. *Diameter measurements.* Multiple inner diameters were measured and averaged in each cross-section of rat and human airway rings.

### Statistical analysis

For each metric, the average of the entire FOV was determined, and by averaging this value across all FOVs taken from all samples of the same condition, a single value for each metric was established for each donor and condition. See **Supplementary Table S2** for donor numbers for each condition and measurement. Where noted in the figures, statistical analysis of differences between the mean values was performed using the two-sided, unpaired t-test. Unless noted differently in legend, boxplots consist of the following elements: center line, median; box limits, upper and lower quartiles; whiskers, extreme data points. Where error bars are present, they represent standard deviation (STD) or standard error of the mean (SEM), as noted in caption.

### Linear regression model

The model was created in MATLAB (Mathworks; Version 2023b) using the regression learner application.

### Data for comparative CPB and cell type composition map

*Mouse tracheal CPB.* CPB in developing mouse trachea between postnatal days (P) 0 and 29 (grey crosses in Fig.6b) in were derived from Toskala *et al*^86^. CPB in mature mouse trachea at P15 (black cross in Fig.6b) was derived from Ramirez-San Juan *et al*^14^.

#### Airway-on-Chip and insert cultures of primary human airway epithelial cells in BD

CPB and cell type composition were derived from our previous study^49^.

#### IL-13 treatment of primary human airway epithelial cultures in Vertex ALI medium

Cells (N=1 donor) were differentiated in Vertex ALI medium^59^ for 21 days at ALI. A chronic airway inflammatory phenotype was induced by treatment with 100 ng mL^−1^ of IL-13 (Invitrogen, A42525) for 14 days. Analysis of CPB and cell type composition was conducted at day 35 ALI as described above.

#### Generation and analysis of iPSC-derived respiratory epithelial cultures (iALI cultures)

Differentiation of human iPSCs (hiPSCs) towards respiratory epithelium was performed as described previously^58^. Briefly, hiPSCs from N=1 donor were differentiated to definitive endoderm by using the STEMdiff^TM^ Definitive Endoderm Kit (STEMCELL Tech., Vancouver, BC, Canada). Subsequently, cells were dissociated and replated for anterior foregut induction by supplementing basis medium with 10 µM SB431542 (provided by the Institute of Organic Chemistry, Leibniz University, Hannover, Germany), and 3 µM Dorsomorphin (Sigma Aldrich, Saint Louis, MO, USA) for 24 hours, followed by supplementation with 2 µM IWP2 (Tocris, Bristol, UK) and 10 µM SB431542 for another 24 h. For lung lineage specification, basis medium was supplemented with 10 ng mL^−1^ BMP4 (R&D Systems, Minneapolis, MN, USA), 10 ng mL^−1^ FGF10 (R&D Systems, Minneapolis, MN, USA), and 3 µM Chir99021 (provided by the Institute of Organic Chemistry, Leibniz University, Hannover, Germany) until day 14 of differentiation. NKX2.1 positive lung progenitor cells were enriched by sorting for the cell surface marker carboxypeptidase M (CPM) (FUJIFILM Wako, Cat# 014-27501). To mature lung progenitor cells to ciliated respiratory epithelium, enriched cultures were seeded onto transwells (Greiner Bio-One, Frickenhausen, Germany) and expanded in small airway epithelial cell growth medium (SAECGM; PromoCell, Heidelberg, Germany) supplemented with 1% penicillin/streptomycin (Gibco, Billings, MT, USA), 1 µM A83-01 (Tocris, Bristol, UK), 0.2 µM DMH-1 (Tocris, Bristol, UK) and 5 µM Y-27632 (Tocris, Bristol, UK) for four days. Afterwards, medium was switched to PneumaCult^TM^-ALI medium (STEMCELLTech., Vancouver, BC, Canada) and cells were differentiated in air-liquid interface conditions for 28 days before analysis. Analysis of CPB and cell type composition was conducted as described above for primary human airway epithelial cultures.

### Physics-based model

#### Cell-level ciliary input parameters

We model the averaged forces generated by all cilia of a multiciliated cell as a single force monopole, located at one cilia length above a no-slip cell surface at z=0, in a semi-infinite domain, based on custom MATLAB (Mathworks; Version 2023b) implementation of the regularized Stokeslet algorithm^87^. The regularized Stokeslet’s strength is proportional to ciliary beat amplitude, with its direction corresponding to power stroke direction (**Supplementary Fig. 7A**). While this approach cannot resolve the flow and coordination of individual cilia as in single-cilia models^42,88–91^, it is straightforward to implement, suitable for large number of ciliated cells, and directly takes into account of the wall-screening effects due to finite cilia length comparing to the slip-boundary-velocity approach^14^. We did not consider the effects of double confinement or mucus film geometry discussed in Ramirez-San Juan *et al.* ^14^ because (i) we intend to compare tracer particle motions recorded at the cilia tip and sufficiently far from other confinement boundaries such as coverslip and air-liquid interface; (ii) experimentally, we chose appropriately-sized field-of-views such that no recirculation effects are observable; (iii) all functional measurements were done in washed samples without the presence of mucus.

#### Tissue-level ciliary input parameters

We derive the ciliated cell distribution based on a cilia coverage percentage and a crystalline order parameter as defined previously^14^, where the order parameter relies on the distribution of wavelength λ between each ciliated patch (**Supplementary Fig. 7A**). Here the mean and standard deviation of λ is determined based on structural measurements. Then ciliated patches are generated based on Gaussian displacement of cells that follow regular crystalline patterns, similar to the procedure reported in Ramirez-San Juan *et al* ^14^. Each ciliated cell assumes a beat direction angle θ sampled from a Von Mises distribution, where its mean is set to be 0 (beating towards *x-*axis) and its second moment, or the orientation order parameter, is determined based on the measured cilia beat order. In the **Supplementary Methods**, we provide an exact description of how both cilia- and tissue-level parameters are implemented *in silico*.

#### Model output

The model uses a rectangular grid of 51 x 51 cells, doubly periodic in both *x-* and *y-*axis, where *x-*axis is defined as the clearance direction. Every cell is assumed to have a diameter of 10 µm, with its center slightly perturbed away from the exact grid points in Monte-Carlo simulations; for visual reference, cell boundaries are drawn based on the Voronoi diagram of the perturbed cell center points (**Supplementary Fig. SB**). We implement the periodic boundary conditions by truncating hydrodynamic interactions further than one periodic image away (>250 µm) from any given point of interest. This introduces only a small error in the flow velocity because the no-slip surface at *z=*0 causes a quadratic decay of hydrodynamic interactions. We derive ciliary flow characteristics from the trajectories of simulated tracer particles injected near cilia tip. Tracers are subject to both cilia-driven flow and random fluctuations. All flow experiments were done using 1 µm diameter particles suspended in buffer after the mucus was removed. For predictions associated with *ex vivo* measurements, we assume the random fluctuation is due to only the thermal diffusivity of water at room temperature ( 20°C; *D=*0.4 µm^2^ s^−1^). For predictions related to *in vitro* experiments, we accounted for additional noise, possibly due to immature ciliary beat, and used effective diffusivity scales estimated using particle tracks (**Supplementary Methods; Supplementary Fig. 15**). Time evolution of 500 initially uniformly distributed particles are computed for 500 beat periods, following the Langevin equation d**r**/d*t* = **v**(**r**) + (2*D*)^0.5^η(*t*), where **r** is the particle position, **v**(**r**) the cilia-driven flow, and η(*t*) a standard Wiener process (white noise). Equations are numerically integrated with a Euler-Maruyama scheme, using periodic boundaries in both *x-* and *y*-directions. The main quantitative output of our simulation is the clearance per beat (CPB) and clearance directionality (**Supplementary Fig. 7A**).

#### Example flow pattern

To illustrate how tissue-level parameters affect our model output, we present example case studies (**Supplementary Fig. 7B**). The crystalline order parameter changes how ciliated cells are distributed; however, it does not strongly impact the characteristics of the tracer trajectories. Lowering cilia coverage or orientation order parameters reduces the clearance distance and directionality. In **Supplementary Fig. 8**, we present quantitative results of CPB, and clearance directionality change with (1) cilia length, (2) cilia beat amplitude, (3) cilia beat order and (4) patch heterogeneity.

## DATA AVAILABILITY

The raw measurements and consolidated summary data, as well as example microscopy footage, generated in this study have been deposited in the Figshare database under https://doi.org/10.6084/m9.figshare.24989700. Source data for each experimental figure panel are provided with this paper.

## CODE AVAILABILITY

The custom MATLAB script used to generate the model prediction results for human/rat benchmark (Fig. 3e), in vitro culture (Fig. 5c), and parametric studies (Supplementary Figs. 7, 8, 9C, 14E and F, 15A) has been deposited in a Zenodo repository (DOI: 10.5281/zenodo.14684411). Detailed descriptions of the underlying mathematical algorithm are provided in the supplementary information and within the same Zenodo repository. The code is released under MIT license and is freely accessible without restrictions.

## ACKNOWLEDGEMENTS

This study was funded by (1) the National Institute of Health: National Heart, Lung, and Blood Institute (NIH:NHLBI R01 HL152633, A.L.R., J.N., and E.K.), (2) The Hastings Foundation (A.L.R.), (3) the European Research Council (ERC: ERC-STG 950219, J.N.), (4) the European Molecular Biology Organization (EMBO: Scientific Exchange Grant 9101; D.R.), (5) Technical University Munich (TUM: Internationalization Grant, D.R.) and (6) German Center for Lung research (DZL: 82DZL002A1, R.O.). We also want to thank Sandra Sühnel (excision of rat lungs at TUM) and Jackie Mao (excision of rat lungs at USC).

## AUTHOR CONTRIBUTIONS

D.R. and T.S. contributed equally to the work. J.N., A.L.R. and E.K. conceptualized the project. J.N. and A.L.R. contributed equally to the work. J.N., A.L.R. and E.K. provided funding for this study. J.N., D.R., T.S., C.N.S, E.J.Q., B.A.C., A.D., T.G., N.T., S.G., A.S., L.S., and R.O. performed and oversaw the experiments, and D.R., T.S., and J.N. performed the data analysis. F.L., E.K. and J.N. formulated the theoretical model. J.N., F.L. and A.L.R. wrote the paper and all authors reviewed and edited the paper. Correspondence and requests for materials should be addressed to J.N. or A.L.R.

