## Supplemental Information for "Structure and Function Relationships of Mucociliary Clearance in Human and Rat Airways"

| COMMERCIAL HUMAN CELLS | | | | | | | |
|---|---|---|---|---|---|---|---|
| Internal ID | Supplier | Catalog number | Cell Type | Lot number | Biological Sex | Age | Cause of Death |
| HSA8938 | Lifeline Cell Technologies | FC-0016 | HSAEC | 08938 | Female | 56 y.o. | CVA/Stroke |
| HB6288 | | FC-0035 | HBTEC | 06288 | Female | 70 y.o. | CVA/2nd ICH |
| HB7783 | | FC-0035 | HBTEC | 07783 | Male | 12 y.o. | SIGSW |
| HB8265 | | FC-0035 | HBTEC | 08265 | Female | 18 y.o. | Head Trauma |
| HB9439 | | FC-0035 | HBTEC | 09439 | Male | 48 y.o. | CVA/Stroke |

| HUMAN TISSUE SAMPLES | | | | | | |
|---|---|---|---|---|---|---|
| Internal ID | Site of sample analysis | Tissue Type | Cell Type Isolated | Biological Sex | Age | Cause of Death Or Tissue Collection |
| H44 | USC | Whole Lung | HBTEC | Female | 73 y.o. | Cause of Death: Head Trauma |
| H47 | USC | Whole Lung | HBTEC | Male | 75 y.o. | Cause of Death: Stroke |
| H48 | USC | Whole Lung | HBTEC | Male | 63 y.o. | Cause of Death: Anoxia, Cerebrovascular accident |
| 2124 | University of Iowa | Trachea Ring | | Male | 65 y.o. | Cause of Death: Cardiac Arrest |
| 2624 | University of Iowa | Trachea Ring | | Male | 62 y.o. | Cause of Death: CVA/Stroke |
| 2924 | University of Iowa | Trachae Ring | | Female | 63 y.o. | Cause of Death: CVA/Stroke |
| BR640<br>BR641<br>BR643<br>BR644<br>BR646<br>BR647<br>BR648<br>BR650<br>BR652 | LUMC | Bronchial Rings | Not Applicable | Data Not Available | | Cause of Tissue Collection: Resection for lung cancer treatment |

| RAT TISSUE SAMPLES | | | |
|---|---|---|---|
| Internal ID | Site of sample analysis | Breed | Biological Sex |
| R#USC | USC | Wistar-Kyoto rats (Charles River) | F |
| R#TUM | Helmholtz/TUM | Wistar rats (Charles River) | F |

**Supplemental Table 1: Human and rat donor information**

#### Number (N) of donors per condition and measurement

| In vitro condition - human | | Cell type composition | Muc5B analysis | Clearance per beat | Clearance directionality | Ciliary Beat Orientational Order Parameter | Cilia Length | Beat Amplitude | Patchiness | Crystalline Order Parameter (COP) |
|---|---|---|---|---|---|---|---|---|---|---|
| BD | | 6 | 4 | 4 | 4 | 3 | 3 | 3 | 3 | 3 |
| PC | | 6 | 4 | 4 | 4 | 3 | 3 | 3 | 3 | 3 |
| PCS | | 3 | 4 | 3 | 3 | 3 | 3 | 3 | 3 | 3 |
| SAGM | | 3 | 4 | 3 | 3 | 3 | 3 | 3 | 3 | 3 |
| mAir | | 3 | 4 | 3 | 3 | 3 | 3 | 3 | 3 | 3 |
| L MEDIA- TOTAL UNIQUE DONORS | | 6 | 4 | 4 | 4 | 3 | 3 | 3 | 3 | 3 |
| Ex vivo BG-human | | Cell type composition | Muc5B analysis | Clearance per beat | Clearance directionality | Ciliary Beat Orientational Order Parameter | Cilia Length | Beat Amplitude | Patchiness | Crystalline Order Parameter (COP) |
| BG0 | | 3 | 3 | 2 | 3 | 3 | | 3 | 3 | 3 |
| BG1 | | 3 | | 1 | 1 | 1 | | 1 | 1 | |
| BG2 | | 5 | | 1 | 1 | 1 | 2 | 1 | 1 | 1 |
| BG3 | | 3 | | | | | | | | |
| BG4 | | 7 | | | | | 5 | | | |
| BG5 | | 2 | | | | | | | | |
| BG6 | | 2 | | 1 | 1 | 1 | | | 1 | 1 |
| BG0-6 TOTAL UNIQUE DONORS | | 11 | 3 | 4 | 4 | 4 | 7 | 3 | 4 | 4 |
| Ex vivo BG-rat | Ciliation Only | Cell type composition | | Clearance per beat | Clearance directionality | Ciliary Beat Orientational Order Parameter | Cilia Length | Beat Amplitude | Patchiness | Crystalline Order Parameter (COP) |
| BG0 | 8 | 5 | | 4 | 4 | 4 | 5 | 4 | 6 | 6 |
| BG1 | 4 | 3 | | 1 | 1 | | | | 1 | |
| BG2 | 3 | 2 | | | | | | | | |
| BG3 | 4 | - | | | | | | | | |
| BG4 | 3 | 1 | | | | | | | | |
| BG5 | 3 | 1 | | | | | | | | |
| BG0-1 TOTAL UNIQUE DONORS | 7 | 4 | | 4 | 4 | 4 | 5 | 4 | 6 | 6 |
| BG0-5 TOTAL UNIQUE DONORS | 12 | 5 | | 4 | 2 | 4 | 5 | 2 | 4 | 4 |

**Supplemental Table 2: Detailed numbers of donors per condition and measurement.**

| Species | Donor | BG | M/E | Diameter [mm] | % Ciliated Cells | % Secretory Cells | of which | % Muc5AC+ CC16- | % Muc5AC- CC16+ | % Muc5AC+ CC16+ | | Clearance Speed [µm/s] | Mean Directionality (at R=80 µm) | CBF [Hz] | CBD [µm/beat] | Ciliary Beat OP | Lambda [µm] | STD Lambda [µm] | Cilia Length [µm] | Ciliary Beat Amplitude [µm] | Crystalline OP |
|---|---|---|---|---|---|---|---|---|---|---|---|---|---|---|---|---|---|---|---|---|---|
| HUMAN | H44 | 0 | M | 17 | 86.93528384 | 9.083542607 | | 74.73930195 | 11.50641026 | 13.7542878 | | | | | | | | | | | |
| HUMAN | H44 | 1 | M | 12 | 92.82029307 | 11.7826538 | | 76.14602711 | 9.344273186 | 14.50969971 | | | | | | | | | | | |
| HUMAN | H44 | 2 | M | | 90.44539802 | 2.713351102 | | 25.70874862 | 57.84606866 | 16.44518272 | | | | | | | | | | | |
| HUMAN | H44 | 3 | M | 5.7 | | | | | | | | | | | | | | | | | |
| HUMAN | H44 | 4 | M | 4.4 | 94.66456551 | 8.948423205 | | 7.851851852 | 88.14814815 | 4 | | | | | | | | | | | |
| HUMAN | H44 | 5 | M | 2.5 | | | | | | | | | | | | | | | | | |
| HUMAN | H44 | 6 | M | 2.2 | 94.61045043 | 10.80658299 | | 16.37366983 | 61.06018119 | 22.56614898 | | 21.55126908 | 0.9612 | 3.25 | 6.631159716 | 0.9081 | 26.89 | 4.6536 | | | 0.755255328 |
| HUMAN | H44 | 7 | M | 1.9 | | | | | | | | | | | | | | | | | |
| HUMAN | H47 | 0 | M | 17.5 | 96.64619245 | 14.05901562 | | 29.46217699 | 35.42200374 | 35.11581927 | | 6.010020068 | 0.6289 | 1.3 | 4.62309236 | 0.9509 | 25.8 | 7.353910524 | | 13.44738599 | 0.596899225 |
| HUMAN | H47 | 1 | M | | 97.82553101 | 16.9213766 | | 39.28716303 | 21.23391231 | 39.47892466 | | | | | | | | | | | |
| HUMAN | H47 | 2 | M | 10 | 97.77611402 | 10.8801182 | | 31.31920904 | 30.27966102 | 38.40112994 | | 15.52900872 | 0.881525 | 3.16771425 | 4.90227574 | 0.925725 | 13.8 | 0.707106781 | | 11.03569444 | 0.927536232 |
| HUMAN | H47 | 3 | M | 5 | 97.58096735 | 12.88014879 | | 20.81392162 | 68.78596876 | 10.40010962 | | | | | | | | | | | |
| HUMAN | H47 | 5 | M | 2.4 | 98.52855164 | 19.21864498 | | 26.22076485 | 28.00252845 | 45.7767067 | | | | | | | | | | | |
| HUMAN | H47 | 6 | M | 2 | 96.87439044 | 12.75792107 | | 33.87987013 | 30.5275974 | 35.59253247 | | | | | | | | | | | |
| HUMAN | H48 | 0 | M | | 82.15675179 | 10.52139447 | | 61.87739464 | 23.42389411 | 14.69871125 | | 8.178527591 | 0.782 | 1.9 | 4.304488206 | 0.9177 | 31.3 | 6.646803743 | | | 0.699680511 |
| HUMAN | H48 | 1 | M | | 90.6670409 | 5.015616885 | | 63.47321429 | 29.39285714 | 7.133928571 | | 8.610547065 | 0.79895 | 2.666666667 | 3.228955149 | 0.9214 | 24.8 | | | 13.57946667 | |
| HUMAN | H48 | 2 | M | | 77.86173549 | 3.927507385 | | 98.91754518 | 0 | 1.082454819 | | | | | | | | | | | |
| HUMAN | H48 | 3 | M | | 79.51133452 | 23.76539246 | | 58.75576037 | 29.36507937 | 11.87916027 | | | | | | | | | | | |
| HUMAN | H48 | 4 | M | | 41.07579151 | 13.53222878 | | 50.49135686 | 26.64574947 | 22.86289368 | | | | | | | | | | | |
| HUMAN | H48 | 5 | M | | 80.06022787 | 18.17158735 | | 29.21568627 | 60.11832319 | 10.66599053 | | | | | | | | | | | |
| HUMAN | BR640 | 2 | E | 7.8 | 65.66316283 | 15.5261798 | | 17.88617886 | 76.76790919 | 5.34591195 | | | | | | | | | 7.059074074 | | |
| HUMAN | BR641 | 2 | E | 6.8 | 89.83050847 | 3.95480226 | | 14.28571429 | 0 | 85.71428571 | | | | | | | | | 6.565833333 | | |
| HUMAN | BR643 | 3 | E | 5.7 | 84.59164292 | 12.85018993 | | 14.6780592 | 77.9829112 | 7.3390296 | | | | | | | | | | | |
| HUMAN | BR644 | 4 | E | 3 | 83.1150064 | 9.663558282 | | 40.81724711 | 23.66804139 | 35.5147115 | | | | | | | | | 7.167927083 | | |
| HUMAN | BR646 | 4 | E | 4 | 80.4 | 20.4 | | 50.98039216 | 29.41176471 | 19.60784314 | | | | | | | | | 7.70532 | | |
| HUMAN | BR648 | 4 | E | 4 | 97.13790113 | 1.56114484 | | 38.88888889 | 44.44444444 | 16.66666667 | | | | | | | | | 7.227533333 | | |
| HUMAN | BR650 | 4 | E | 4 | 85.24034694 | 9.014140103 | | 31.36752137 | 45.77018044 | 22.8622982 | | | | | | | | | 7.235893939 | | |
| HUMAN | BR652 | 4 | E | 5 | 90.12546146 | 12.25411774 | | 26.77655678 | 56.79399966 | 16.42944358 | | | | | | | | | 7.712513514 | | |
| HUMAN | H2924 | 0 | M | 14 | 92.7 | | | | | | | 25.33333333 | 0.960739604 | 2.743333333 | 9.439889179 | 0.9558 | 36.33333333 | 4.618802154 | | 10.93333333 | 0.820220936 |
| RAT | USCR1 | 1 | M | | | | | | | | | | | | | | | | | | |
| RAT | USCR2 | | M | 3.4 | | | | | | | | | | | | | | | | | |
| RAT | USCR2 | 1 | M | 1.9 | | | | | | | | | | | | | | | | | |
| RAT | USCR2 | 2 | M | 1.2 | 90 | | | | | | | 35.13261462 | | 6.744791667 | 5.20885097 | | | | | | |
| RAT | USCR2 | 3 | M | 0.6 | | | | | | | | | | | | | | | | | |
| RAT | USCR2 | 4 | M | 0.4 | | | | | | | | | | | | | | | | | |
| RAT | USCR3 | 0 | M | | 90 | | | | | | | 25.21354548 | | 7.6171875 | 3.310085971 | | | | | | |
| RAT | USCR10 | 0 | M | | 49.85286618 | 2.570204207 | | 36.47727273 | 62.95454545 | 0.568181818 | | | | | | | | | | | |
| RAT | USCR10 | 1 | M | | 60.44862045 | 8.342046467 | | 43.16908582 | 54.60911637 | 2.22179781 | | | | | | | | | 6.382526316 | | |
| RAT | USCR10 | 2 | M | | 53.6121948 | 0.857791764 | | 66.05392157 | 33.94607843 | 0 | | | | | | | | | | | |
| RAT | USCR20 | 0 | M | 3.4 | 54.39345305 | 0.321942303 | | 0 | 100 | 0 | | | | | | | | | | | |
| RAT | USCR20 | 1 | M | 2.2 | 71.71178932 | 4.01386496 | | 27.75862069 | 71.37931034 | 0.862068966 | | | | | | | | | | | |
| RAT | USCR20 | 2 | M | 1.6 | | | | | | | | | | | | | | | | | |
| RAT | USCR20 | 3 | M | 0.6 | | | | | | | | | | | | | | | | | |
| RAT | USCR20 | 4 | M | 0.26 | | | | | | | | | | | | | | | | | |
| RAT | USCR30 | 0 | M | | 28.18927851 | 0.033200531 | | 0 | 100 | 0 | | | | | | | | | | | |
| RAT | USCR41 | 1 | M | | 60.53920418 | 6.702123922 | | 4.880952381 | 90.77561328 | 4.343434343 | | | | | | | | | | | |
| RAT | USCR41 | 2 | M | | 60.97560976 | 5.563282337 | | 0 | 96.96969697 | 3.03030303 | | | | | | | | | | | |
| RAT | USCR42 | 0 | M | | 40.83066667 | | | -- | -- | -- | | 4.088044537 | 0.7441 | 6.310992 | 0.60374953 | 0.5114 | 22.65 | 5.444722215 | | 8.890050752 | 0.66004415 |
| RAT | USCR43 | 0 | M | | 50.31 | | | -- | -- | -- | | 6.027563044 | 0.66275 | 4.609979 | 1.160014715 | 0.529114286 | 19.9 | 3.798683983 | | 7.985935484 | 0.730042693 |
| RAT | USCR43 | 1 | M | | 52.88 | | | -- | -- | -- | | 8.659411402 | 0.74665 | 6.2335475 | 1.427284384 | 0.55 | 28 | | | | |
| RAT | USCR44 | 4 | M | | 89.7482297 | 0.31171305 | | 50 | 50 | 0 | | | | | | | | | | | |
| RAT | USCR44 | 5 | M | | 91.27831163 | 0.870972854 | | 51.92307692 | 48.07692308 | 0 | | | | | | | | | | | |
| RAT | USCR45 | 0 | M | 3.5 | | | | | | | | | | | | | | | | | |
| RAT | USCR45 | 1 | M | 2.2 | | | | | | | | | | | | | | | | | |
| RAT | USCR45 | 2 | M | 1.7 | | | | | | | | | | | | | | | | | |
| RAT | USCR45 | 3 | M | 1 | | | | | | | | | | | | | | | | | |
| RAT | USCR45 | 4 | M | 0.45 | | | | | | | | | | | | | | | | | |
| RAT | USCR45 | 5 | M | 0.2 | | | | | | | | | | | | | | | | | |
| RAT | TUMR01 | 0 | M | | 35.192 | | | -- | -- | -- | | 3.285405793 | | 2.845629667 | 1.168630255 | 0.869866667 | 28.25 | 1.522333735 | | | 0.923790973 |
| RAT | TUMR01 | 3 | M | | 67.03802393 | | | -- | -- | -- | | | | | | | | | | | |
| RAT | TUMR01 | 4 | M | | 89.71195916 | | | -- | -- | -- | | | | | | | | | | | |
| RAT | TUMR01 | 5 | M | | 95.63409563 | | | -- | -- | -- | | | | | | | | | | | |
| RAT | TUMR02 | 0 | M | | 36.13666667 | | | -- | -- | -- | | 7.3 | | 4.6171405 | 1.553250969 | 0.8678 | 32.5 | 9.192388155 | | | 0.6 |
| RAT | TUMR02 | 4 | M | | 88.68125701 | | | -- | -- | -- | | | | | | | | | | | |
| RAT | TUMR02 | 5 | M | | 92.44537154 | | | -- | -- | -- | | | | | | | | | | | |
| RAT | TUMR03 | 3 | M | | 70.5504303 | | | -- | -- | -- | | | | | | | | | | | |
| RAT | TUMR04 | 3 | M | | 76.32028864 | | | -- | -- | -- | | | | | | | | | | | |
| RAT | TUMR05 | 3 | M | | 76.99918824 | | | -- | -- | -- | | | | | | | | | | | |
| RAT | TUMR56 | 0 | M | 57.5 | | | | | | | | 4.2 | 0.53 | 3.325 | 1.306917087 | 0.60025 | 28.5 | 7.778174593 | | 6.45 | 0.614035088 |
| RAT | TUMR56 | 0 | M | 39.33333333 | | | | | | | | 3.371683333 | 0.498333333 | 4.103333333 | 0.800370651 | 0.14345 | 36 | 8.485281374 | | 6.05 | 0.666666667 |
| RAT | TUMR55 | 0 | M | 45.8 | | | | | | | | 2.62072 | 0.37462 | 4.648 | 0.595148396 | 0.4499 | 30 | | | 6.9 | |
| RAT | TUMR55 | 0 | M | 32.33333333 | | | | | | | | 3.143333333 | 0.306633333 | 3.0875 | 0.807418677 | 0.2509 | 30.5 | 9.192388155 | | 6.5 | 0.573770492 |

**Supplemental Table 3: Detailed *ex vivo* data on cell type composition and clearance function.** Shown are mean values. Donor names are internal IDs. M/E: Airway branching generation was either measured/known (M) or estimated (E) based on trend in Fig. S1. STD: Standard deviation. Missing values indicate unavailability of data.

| Species | Donor | Cell Type | Conditon | % Ciliated Cells | % Secretory Cells | of which | % Muc5AC+ CC16- | % Muc5AC- CC16+ | % Muc5AC+ CC16+ | | Motile Cilia Density | Clearance Speed [µm/s] | Mean Directionality (over 80 µm) | CBF [Hz] | CBD [µm/beat] | Ciliary Beat OP | Lambda [µm] | STD Lambda [µm] | Crystalline OP | Cilia Length [µm] | Ciliary Beat Amplitude [µm] |
|---|---|---|---|---|---|---|---|---|---|---|---|---|---|---|---|---|---|---|---|---|---|
| HUMAN | H44 | HBEC | BD-28DALI | 54.11 | 12.47 | | 19.91 | 76.63 | 3.46 | | | | | | | | | | | | |
| HUMAN | H44 | HBEC | PC-28DALI | 92.93 | 8.53 | | 19.08 | 57.82 | 23.09 | | | | | | | | | | | | |
| HUMAN | H47 | HBEC | BD-28DALI | 39.06 | 13.14 | | 23.49 | 53.15 | 23.36 | | 39.70 | 10.69 | 0.51 | 10.65 | 1.01 | | | | | | |
| HUMAN | H47 | HBEC | PC-28DALI | 74.12 | 16.04 | | 53.11 | 19.72 | 27.17 | | | 28.83 | 0.75 | 6.33 | 4.52 | | | | | | |
| HUMAN | H48 | HBEC | BD-28DALI | 33.04 | 13.88 | | 23.89 | 52.39 | 23.72 | | 73.20 | | | | | | | | | | |
| HUMAN | H48 | HBEC | PC-28DALI | 65.88 | 4.40 | | 53.79 | 37.93 | 8.28 | | | | | | | | | | | | |
| HUMAN | HB7783 | HBEC | BD-28DALI | 35.26 | 2.39 | | 3.61 | 77.50 | 2.22 | | 20.00 | 18.34 | 0.59 | 14.65 | 1.25 | 0.62 | 17.50 | 3.55 | 0.71 | 5.46 | 10.28 |
| HUMAN | HB7783 | HBEC | mAir-28DALI | 62.38 | 5.96 | | 1.19 | 98.46 | 0.00 | | 53.00 | 32.00 | 0.51 | 16.26 | 1.97 | 0.72 | 15.38 | 4.07 | 0.63 | 6.28 | 9.34 |
| HUMAN | HB7783 | HBEC | PC-28DALI | 87.69 | 5.79 | | 13.87 | 69.73 | 16.39 | | 92.38 | 66.38 | 0.80 | 14.39 | 4.61 | 0.74 | 14.63 | 1.69 | 0.84 | 6.51 | 12.55 |
| HUMAN | HB7783 | HBEC | PCS-28DALI | 71.04 | 4.09 | | 9.90 | 41.82 | 48.28 | | 67.40 | 32.27 | 0.58 | 16.26 | 1.98 | 0.72 | 14.88 | 2.64 | 0.75 | 7.46 | 6.92 |
| HUMAN | HB7783 | HBEC | SAGM-28DALI | 16.61 | 0.23 | | 25.00 | 41.67 | 16.67 | | 5.40 | 0.00 | NAN | 11.58 | NAN | 0.64 | 27.50 | 10.23 | 0.47 | 4.78 | 6.51 |
| HUMAN | HB8265 | HBEC | BD-28DALI | 8.85 | 2.16 | | 11.72 | 88.28 | 0.00 | | 23.62 | 14.94 | 0.68 | 16.20 | 0.92 | 0.83 | 24.63 | 9.21 | 0.47 | 5.08 | 10.30 |
| HUMAN | HB8265 | HBEC | mAir-28DALI | 11.70 | 9.71 | | 3.40 | 96.14 | 0.45 | | 12.80 | 15.31 | 0.60 | 18.09 | 0.85 | 0.71 | 22.00 | 6.56 | 0.58 | 6.34 | 8.51 |
| HUMAN | HB8265 | HBEC | PC-28DALI | 84.22 | 7.87 | | 14.22 | 77.08 | 8.71 | | 93.69 | 135.62 | 0.90 | 16.29 | 8.32 | 0.64 | 13.00 | 2.14 | 0.77 | 7.52 | 11.64 |
| HUMAN | HB8265 | HBEC | PCS-28DALI | 63.18 | 9.10 | | 14.02 | 58.31 | 27.67 | | 56.00 | 16.78 | 0.69 | 14.80 | 1.13 | 0.79 | 15.63 | 1.77 | 0.84 | 7.44 | 7.66 |
| HUMAN | HB8265 | HBEC | SAGM-28DALI | 1.53 | 3.45 | | 39.71 | 59.86 | 0.43 | | 6.48 | 0.00 | NAN | 0.00 | NAN | 0.00 | 33.67 | 7.23 | 0.70 | 4.01 | 7.32 |
| HUMAN | HSA8938 | HSAEC | BD-28DALI | 69.27 | 3.76 | | 40.73 | 57.92 | 1.35 | | 63.48 | 45.78 | 0.61 | 14.63 | 3.13 | 0.81 | 18.50 | 2.14 | 0.84 | 7.14 | 7.50 |
| HUMAN | HSA8938 | HSAEC | mAir-28DALI | 78.30 | 4.40 | | 15.98 | 76.51 | 7.50 | | 75.44 | 72.48 | 0.82 | 16.66 | 4.35 | 0.85 | 17.00 | 0.82 | 0.93 | 7.84 | 6.16 |
| HUMAN | HSA8938 | HSAEC | PC-28DALI | 90.78 | 6.82 | | 22.39 | 46.02 | 31.59 | | 93.24 | 139.64 | 0.92 | 15.98 | 8.74 | 0.90 | 13.25 | 1.49 | 0.84 | 6.97 | 14.30 |
| HUMAN | HSA8938 | HSAEC | PCS-28DALI | 62.82 | 5.06 | | 2.53 | 91.11 | 6.37 | | 61.57 | 59.74 | 0.84 | 14.60 | 4.09 | 0.70 | 14.00 | 1.20 | 0.88 | 8.99 | 7.73 |
| HUMAN | HSA8938 | HSAEC | SAGM-28DALI | 56.33 | 0.38 | | 65.76 | 13.03 | 4.55 | | 37.21 | 26.85 | 0.64 | 14.64 | 1.83 | 0.70 | 23.43 | 3.74 | 0.77 | 6.88 | 7.42 |
| HUMAN | HB6288 | HBEC | BD-28DALI | 14.46 | 7.04 | | 11.76 | 87.10 | 1.14 | | 10.60 | 7.16 | 0.67 | 14.79 | 0.72 | 0.90 | 37.25 | 6.08 | 0.77 | 6.06 | 8.93 |
| HUMAN | HB6288 | HBEC | mAir-28DALI | 3.98 | 13.87 | | 2.54 | 96.28 | 1.18 | | 6.20 | 5.26 | 0.60 | 17.77 | 0.35 | 0.65 | 29.67 | 10.50 | 0.50 | 6.30 | 8.36 |
| HUMAN | HB6288 | HBEC | PC-28DALI | 66.47 | 11.46 | | 1.71 | 86.77 | 11.52 | | 82.60 | 15.47 | 0.68 | 13.95 | 5.92 | 0.24 | 25.48 | 9.00 | 0.50 | 6.99 | 10.92 |
| HUMAN | HB6288 | HBEC | PCS-28DALI | 45.43 | 19.93 | | 4.07 | 90.12 | 5.81 | | 33.80 | 10.96 | 0.54 | 14.79 | 2.28 | 0.77 | 22.02 | 2.20 | 0.86 | 6.79 | 6.49 |
| HUMAN | HB6288 | HBEC | SAGM-28DALI | 1.30 | 7.71 | | 36.32 | 52.40 | 11.28 | | 0.00 | 0.00 | NAN | NAN | NAN | NAN | NAN | NAN | NAN | 3.44 | NAN |

**Supplemental Table 4: Detailed *in vitro* data on cell type composition and clearance function.** Shown are mean values. Donor names are internal IDs or supplier IDs. HBEC: Primary human bronchial epithelial cells; HSAEC: Primary human small airway epithelial cells. STD: Standard deviation.

Total Cell Type Composition

| | Number of donors | % Ciliated Cells | SEM | % Secretory Cells | SEM | % Not Identified | SEM |
|---|---|---|---|---|---|---|---|
| BD | 4 | 36.7 | 8.0 | 7.8 | 8.3 | 47.6 | 9.2 |
| mAir | 4 | 39.1 | 18.4 | 8.5 | 2.1 | 36.7 | 19.6 |
| SAGM | 4 | 80.3 | 4.3 | 8.7 | 1.5 | 56.1 | 18.0 |
| PC | 4 | 60.6 | 5.4 | 9.5 | 3.6 | 6.4 | 5.0 |
| PC-S | 4 | 18.9 | 13.0 | 2.9 | 1.8 | 11.5 | 6.9 |
| HUMAN BG0-6 | 11* | 85.5 | 2.6 | 11.0 | 1.5 | 3.5 | 2.2 |
| RAT BG0-5 | 12** | 63.1 | 4.1774 | 2.6 | 0.6 | 38.0 | 5.5 |

Secretory Cell Type Composition

| | % Muc5AC+ CC16- | SEM | % Muc5AC- CC16+ | SEM | % Muc5AC+ CC16+ | SEM |
|---|---|---|---|---|---|---|
| BD | 19.1 | 4.5 | 71.5 | 5.5 | 7.9 | 17.4 |
| mAir | 5.8 | 3.4 | 91.8 | 5.1 | 2.3 | 1.8 |
| SAGM | 25.5 | 7.6 | 56.4 | 8.9 | 18.1 | 3.5 |
| PC | 7.6 | 2.7 | 70.3 | 12.2 | 22.0 | 10.1 |
| PC-S | 41.7 | 8.6 | 41.7 | 10.3 | 8.2 | 3.6 |
| HUMAN BG0-6 | 87.1 | 4.3 | 11.6 | 4.6 | 39.2 | 1.1 |
| RAT BG0-5 | 28.0 | 8.4 | 70.9 | 8.2 | 0.1 | 0.50 |

**Supplemental Table 5: Summary comparison table of luminal cell type composition.** *Number of donors per BG: BG0: 3; BG1:3; BG2:5; BG3:3; BG4:7; BG5:2; BG6:2 .**Number of donors per BG: BG0: 7 (4 only ciliation); BG1: 3 (1 only ciliation); BG2: 2; BG3: 5 (5 only ciliation); BG4: 3 (2 only ciliation); BG5: 3 (2 only ciliation)

| Species | Donor | BG | % Ciliation | CPB [µm/beat] | Directionality R=80µm | CBF [Hz] | MCC_Speed [µm/s] | Beat OP | Lambda [µm] | STD_Lambda [µm] | Ciliary Beat Amplitude [µm] |
|---|---|---|---|---|---|---|---|---|---|---|---|
| RAT | TUMR55 | 0 | 32.33 | 0.81 | 0.31 | 3.09 | 3.14 | 0.25 | 30.50 | 9.19 | 6.50 |
| RAT | TUMR56 | 0 | 39.33 | 0.80 | 0.50 | 4.10 | 3.37 | 0.14 | 36.00 | 8.49 | 6.05 |
| RAT | USCR42 | 0 | 40.83 | 0.60 | 0.74 | 6.31 | 4.09 | 0.51 | 22.65 | 5.44 | 8.89 |
| RAT | TUMR55 | 0 | 45.80 | 0.60 | 0.37 | 4.65 | 2.62 | 0.45 | 30.00 | 0.00 | 6.90 |
| RAT | USCR43 | 0 | 50.31 | 1.16 | 0.66 | 4.61 | 6.03 | 0.53 | 19.90 | 3.80 | 7.99 |
| RAT | USCR43 | 1 | 52.88 | 1.43 | 0.75 | 6.23 | 8.66 | 0.55 | 28.00 | | |
| RAT | TUMR56 | 0 | 57.50 | 1.31 | 0.53 | 3.33 | 4.20 | 0.60 | 28.50 | 7.78 | 6.45 |
| HUMAN | H2924 | 0 | 92.70 | 9.44 | 0.96 | 2.74 | 25.33 | 0.96 | 36.33 | 4.62 | 10.93 |
| HUMAN | H44 | 7 | 94.60 | 7.03 | 0.96 | 3.50 | 24.60 | 0.82 | | | |
| HUMAN | H47 | 2 | 97.80 | 8.50 | 0.98 | 2.68 | 22.81 | 0.87 | | | |
| HUMAN MEAN | | | 95.03 | 8.32 | 0.97 | 2.98 | 24.25 | 0.88 | 36.33 | 4.62 | 10.93 |
| HUMAN STD | | | 2.58 | 1.21 | 0.01 | 0.46 | 1.30 | 0.07 | | | |
| RAT MEAN | | | 45.57 | 0.96 | 0.55 | 4.62 | 4.59 | 0.43 | 27.94 | 5.78 | 7.13 |
| RAT STD | | | 8.71 | 0.34 | 0.17 | 1.28 | 2.10 | 0.17 | 5.30 | 3.48 | 1.09 |

**Supplemental Table 6: Benchmark data for model calibration and validation**

| | Total % Secretory cells | SEM | % Muc5B only | SEM | % Muc5AC only | SEM | % SCGB1A1 only | SEM | % Muc5B & Muc5AC | SEM | %SCGB1A1 & Muc5AC | SEM | %SCGB1A1 & Muc5B | SEM |
|---|---|---|---|---|---|---|---|---|---|---|---|---|---|---|
| BD (n= 2 donors) | 28.86 | 11.70 | 4.87 | 1.87 | 40.17 | 16.64 | 28.35 | 16.33 | 10.54 | 1.75 | 12.50 | 1.01 | 1.13 | 0.96 |
| mAir (n= 4 donors) | 25.27 | 5.42 | 6.65 | 1.17 | 13.46 | 5.29 | 68.60 | 2.67 | 2.00 | 0.97 | 6.77 | 1.05 | 1.32 | 0.41 |
| SAGM (n= 2 donors) | 4.81 | 0.97 | 15.64 | 15.64 | 19.79 | 19.79 | 50.00 | 50.00 | 14.57 | 14.5 7 | 0.00 | 0.00 | 0.00 | 0.00 |
| PC (n= 4 donors) | 17.18 | 2.78 | 9.48 | 3.52 | 30.19 | 5.85 | 48.09 | 4.02 | 3.49 | 2.43 | 7.65 | 3.26 | 0.90 | 0.14 |
| PC-S (n= 4 donors) | 14.50 | 3.16 | 6.33 | 1.76 | 22.40 | 2.79 | 58.76 | 3.33 | 1.50 | 0.64 | 9.53 | 3.31 | 0.60 | 0.39 |
| HUMAN BG0 (n= 3 donors) | 13.93 | 1.56 | 18.83 | 7.33 | 29.22 | 5.36 | 38.10 | 7.04 | 1.81 | 0.85 | 8.05 | 2.18 | 2.52 | 0.70 |

**Supplemental Table 7: Summary comparison table of luminal secretory cells including Muc5B**

| Antibody | Supplier | Catalog # | Lot # | Species | Dilution |
|---|---|---|---|---|---|
| Anti-Mucin 5AC | abcam | ab3649 | 1016221-3 | mouse | 1:150 |
| Anti-MUC5B | abcam | ab77995 | 1063495-1 | Mouse | 1:100 |
| Anti-Uteroglobin/SCGB1A1 | Proteintech | 26909-1-AP | 00109398 | rabbit | 1:100 |
| Anti-Acetylated α-tubulin 647 (6-11B-1) | Santa Cruz | sc-23950 | I0921 | mouse | 1:200 |
| Anti-Mucin 5AC 594 | Abcam | ab218363 | GR3446395-2 | rabbit | 1:200 |
| Anti-mouse 568 | Invitrogen | A10037 | | donkey | 1:500 |
| Anti-rabbit 647 | Invitrogen | A21244 | 2433883 | Goat | 1:500 |
| Anti-rabbit 488 | Invitrogen | A11008 | | goat | 1:500 |
| Anti-mouse 405 | Invitrogen | A31553 | 2491371 | goat | 1:500 |
| Anti-mouse 488 | Invitrogen | A32766 | YA367353 | donkey | 1:500 |
| Anti-rabbit 488 | Invitrogen | A21441 | 2387456 | chicken | 1:500 |

**Supplemental Table 8: Antibody Information**

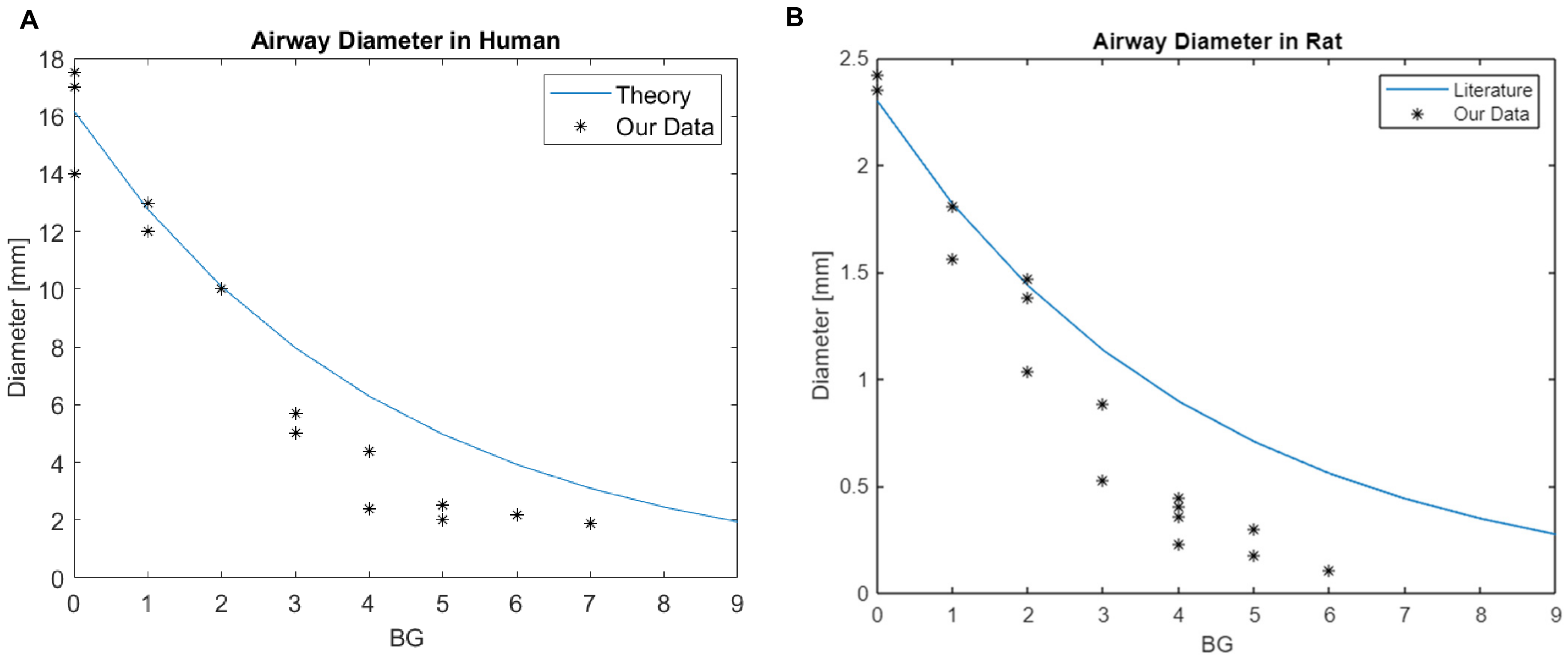

**Supplemental Figure 1: Airway diameters along airway tree in human and rat.**

**A** Human airway diameters of different branching generations (BG) measured in this study roughly follow the theoretical estimates assuming optimality of air conductance. Theoretical model based on [1].

**B** Rat airway diameters measured in this study match estimates published elsewhere. Source data for panels A and B are provided as a Source Data file. Data curve generated from data in [2].

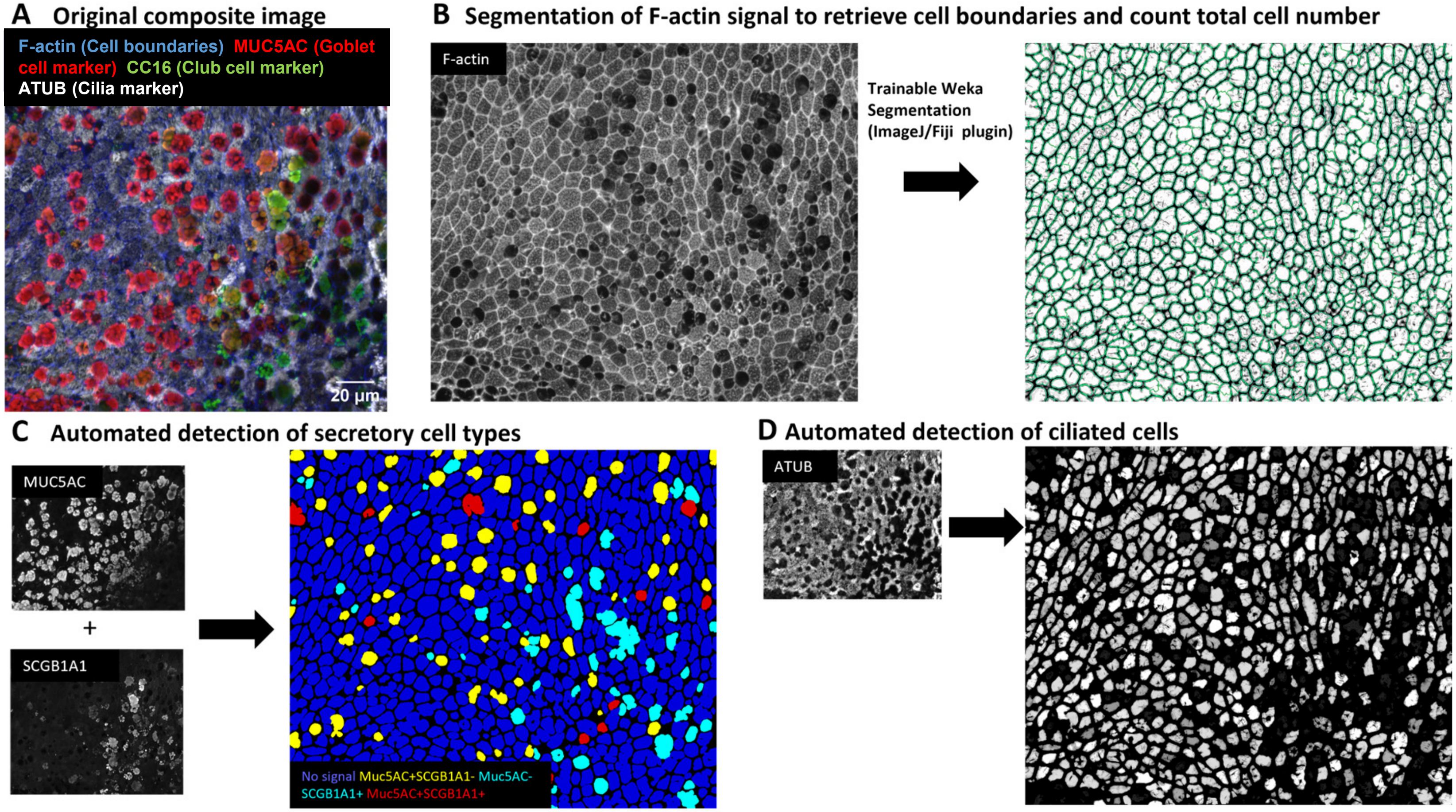

**E Compute cell type percentages. e.g., % ciliated = 100 * # ciliated cells / total cell count**

**Supplemental Figure 2: Automated cell type analysis**

**A** IF-stained luminal surface of human airway epithelium.

**B** Phalloidin staining of field of view in (A) reveals cellular boundaries that are segmented using machine learning, yielding cellular objects and total cell count.

**C and D** Automated detection of secretory and ciliated cell types based on spatial overlay of IF markers and cellular objects detected in (B).

**E** From results in B-D, luminal cell type percentages are computed.

**A**

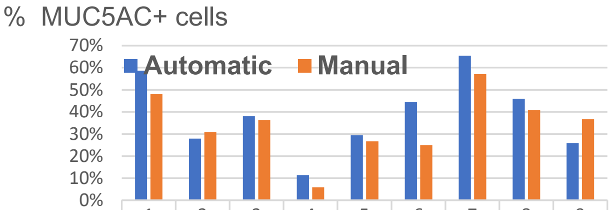

| | 1 | 2 | 3 | 4 | 5 | 6 | 7 | 8 | 9 |
|---|---|---|---|---|---|---|---|---|---|
| Automatic | 59% | 28% | 38% | 11% | 29% | 44% | 65% | 46% | 26% |
| Manual | 48% | 31% | 36% | 6% | 27% | 25% | 57% | 41% | 37% |

**Image number**

Median difference: 5%

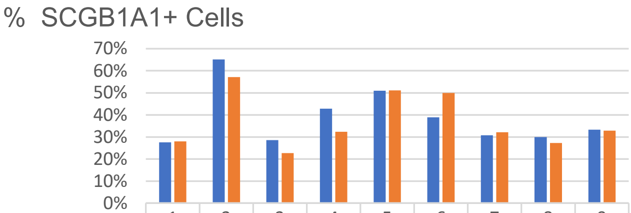

| | 1 | 2 | 3 | 4 | 5 | 6 | 7 | 8 | 9 |
|---|---|---|---|---|---|---|---|---|---|
| Automatic | 28% | 65% | 29% | 43% | 51% | 39% | 31% | 30% | 33% |
| Manual | 28% | 57% | 23% | 32% | 51% | 50% | 32% | 27% | 33% |

**Image number**

Median difference: 6%

**B**

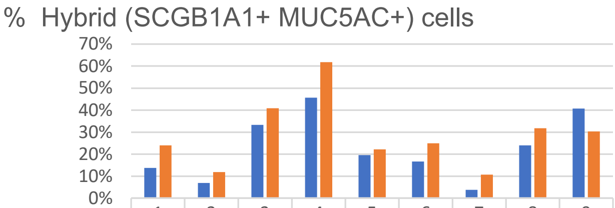

| | 1 | 2 | 3 | 4 | 5 | 6 | 7 | 8 | 9 |
|---|---|---|---|---|---|---|---|---|---|
| Automatic | 14% | 7% | 33% | 46% | 20% | 17% | 4% | 24% | 41% |
| Manual | 24% | 12% | 41% | 62% | 22% | 25% | 11% | 32% | 30% |

**Image number**

Median difference: 15%
(compounded error)

**Supplemental Figure 3: Validation of automated cell type analysis**
**A** Comparison of manual and automated cell type analysis in ten different field of views shows an error of the automated procedure below 10% for MUC5AC and SCGB1A1 positive cells, **B** This error compounds to 15% for double-labeled hybrid cells. Source data for panels A and B are provided as a Source Data file.

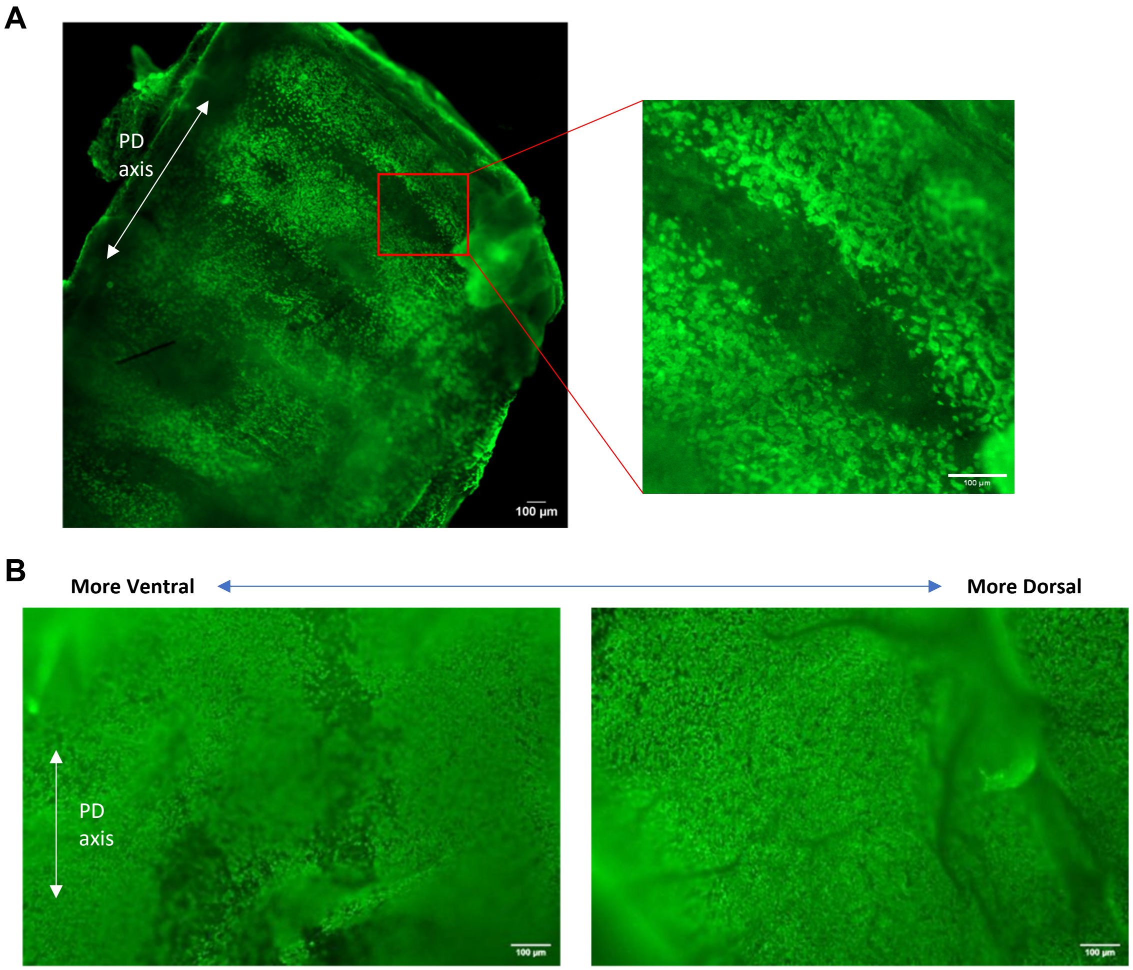

**Supplemental Figure 4: Ciliation patterns in trachea revealed by live staining with fluorescent WGA A** Ventral wall of rat trachea showing regular stripe pattern of densely and sparsely ciliated rings. **B** Two fields of view along ventral wall of same human trachea showing dense ciliation throughout. While there were occasionally patches of lower ciliation, no obvious spatial patterns were discernible. PD, proximal-distal. Scalebars: 100µm.

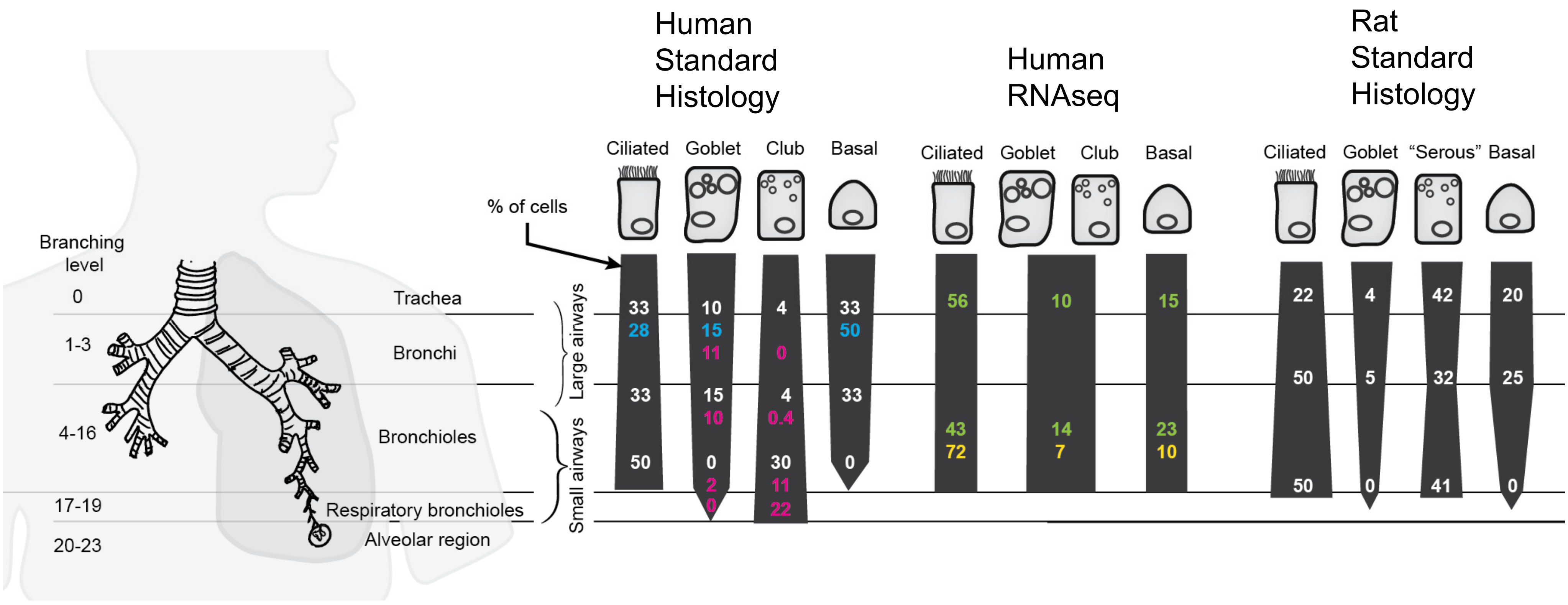

**Supplemental Figure 5: Literature survey of airway epithelial cellular composition.**
The numbers denote the average percentages, and the colors indicate the publication source: white, [3]; pink, [4]; blue, [5]; yellow, [6]; green: [7]. Reference [3-5] are standard histology results. Reference [6,7] are scRNAseq results. If the sources are in consensus, the width of the dark-grey column indicates the relative abundance of the cell type at different levels of the airway tree.

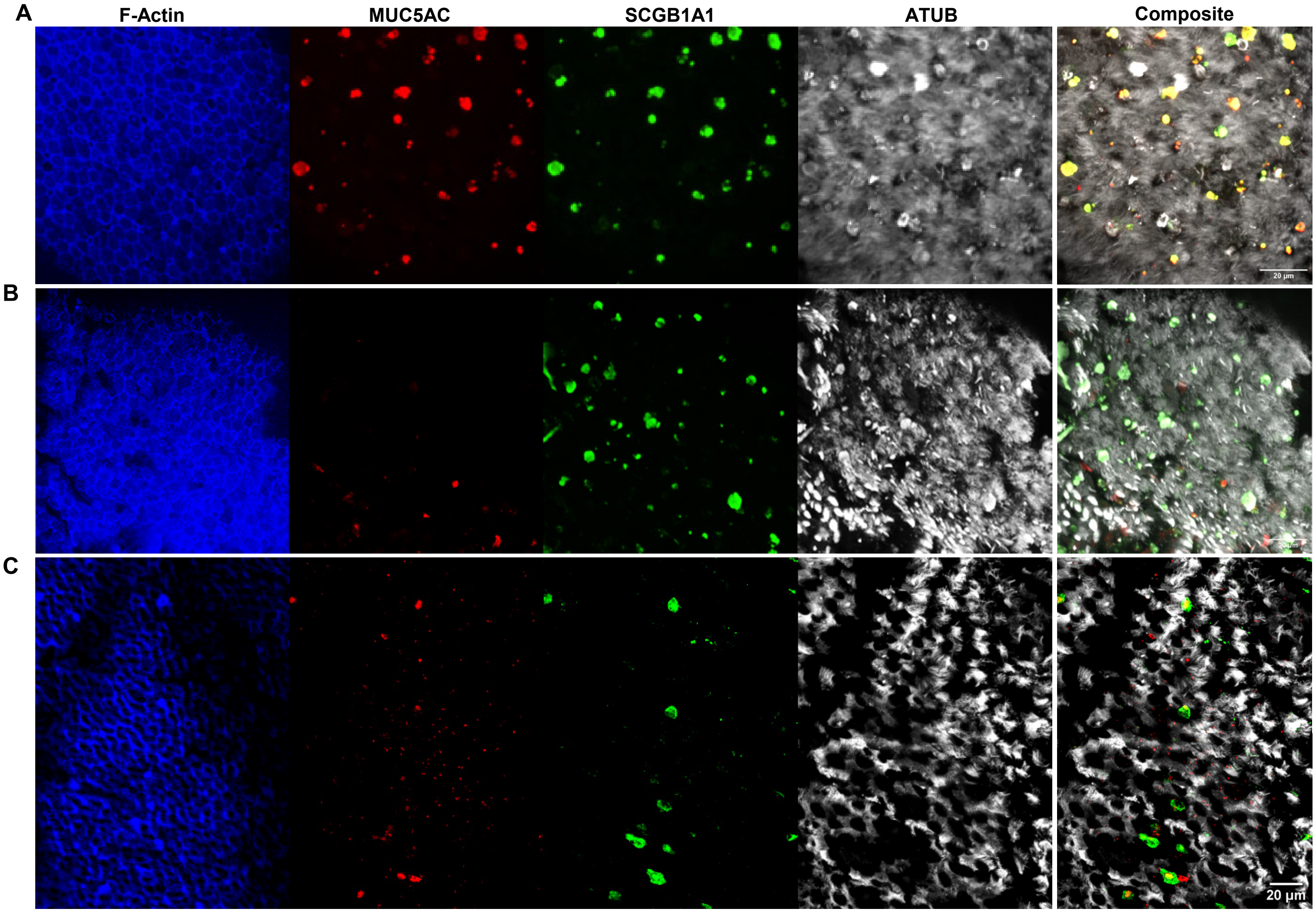

**Supplemental Figure 6: Example images of luminal cell type composition in human and rat airways.**
**A** Human BG4 tissue with high proportion of hybrid secretory cells (MUC5AC+ SCGB1A1+).
**B** Human BG4 tissue with low proportion of hybrid secretory cells (MUC5AC+ SCGB1A1+).
**C** Rat BG0 tissue showing variable and overall sparser levels of ciliated and secretory cells than human tissues.

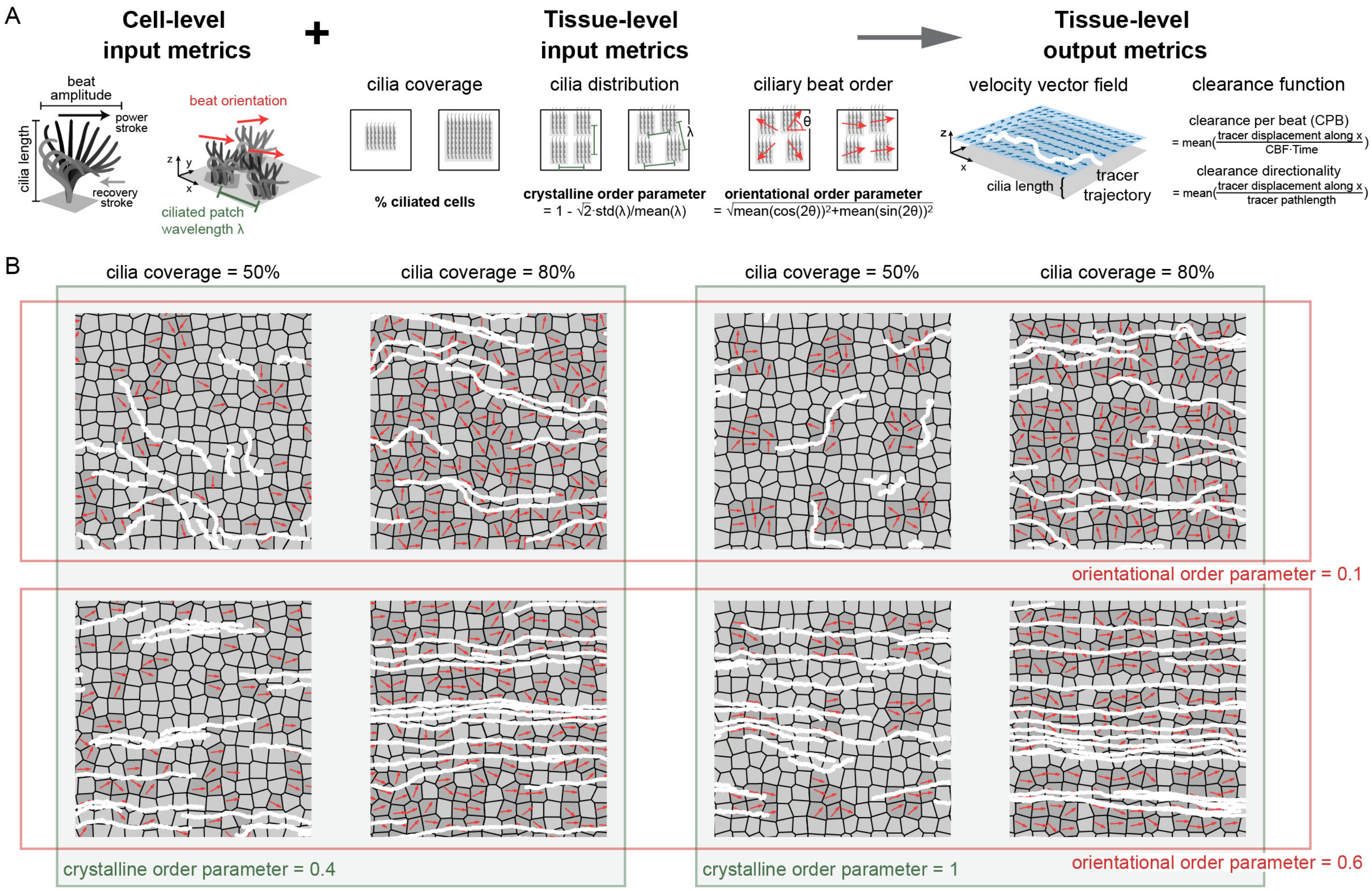

**Supplemental Figure 7: Ciliary input metrics and clearance output metrics of computational model.**
**A** Visualization and definitions of cell and tissue-level ciliary input metrics and tissue-level clearance function in terms of CPB and clearance directionality.
**B** Example predictions of particle clearance trajectories (white) given different combinations of ciliary input metrics. Ciliated cells are shown in dark grey with beat direction indicated by red arrow. Shorter trajectories indicate lower CPB and curved trajectories indicate lower clearance directionality.

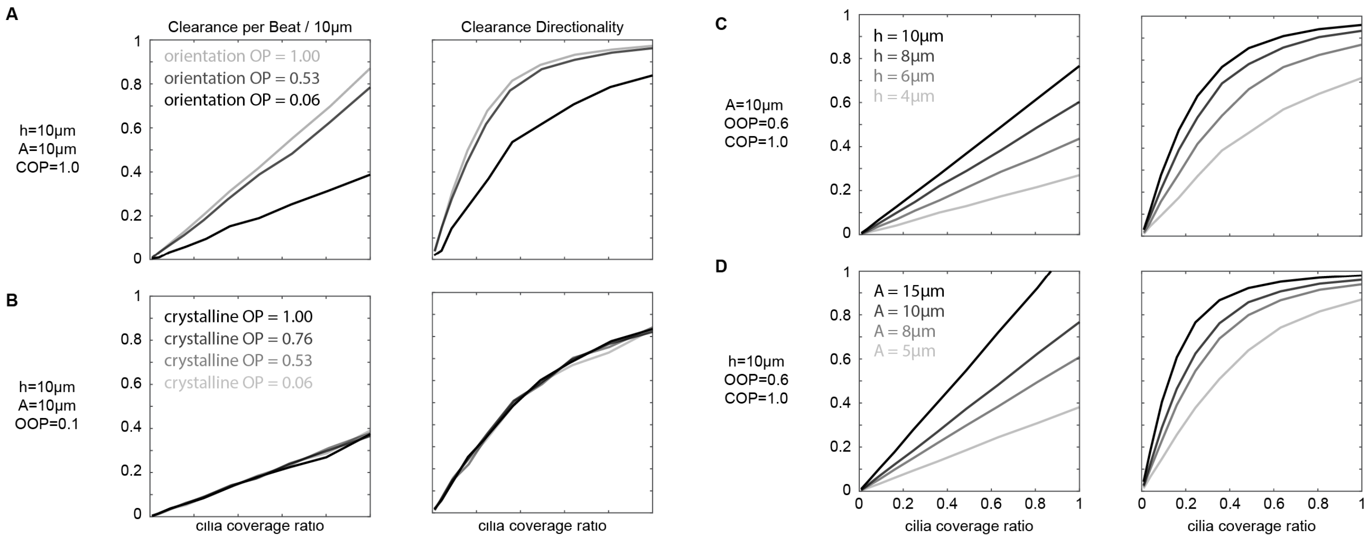

**Supplemental Figure 8: Predicted change of CPB and clearance directionality for different cilia- and tissue-level input parameters.** We tested how dimensionless CPB and clearance directionality change in response to changes in OOP, ciliary order parameter; COP, crystalline order parameter; h, cilia length; and A, ciliary beat amplitude, while keeping the other parameters fixed as noted in figure. **A,** impact of different ciliary order parameters, **B,** impact of different crystalline order parameters, **C,** impact of different cilia lengths, **D,** impact of different ciliary beat amplitudes.For all panels, we use assume unit drag coefficient and regularization parameter at 0.1.

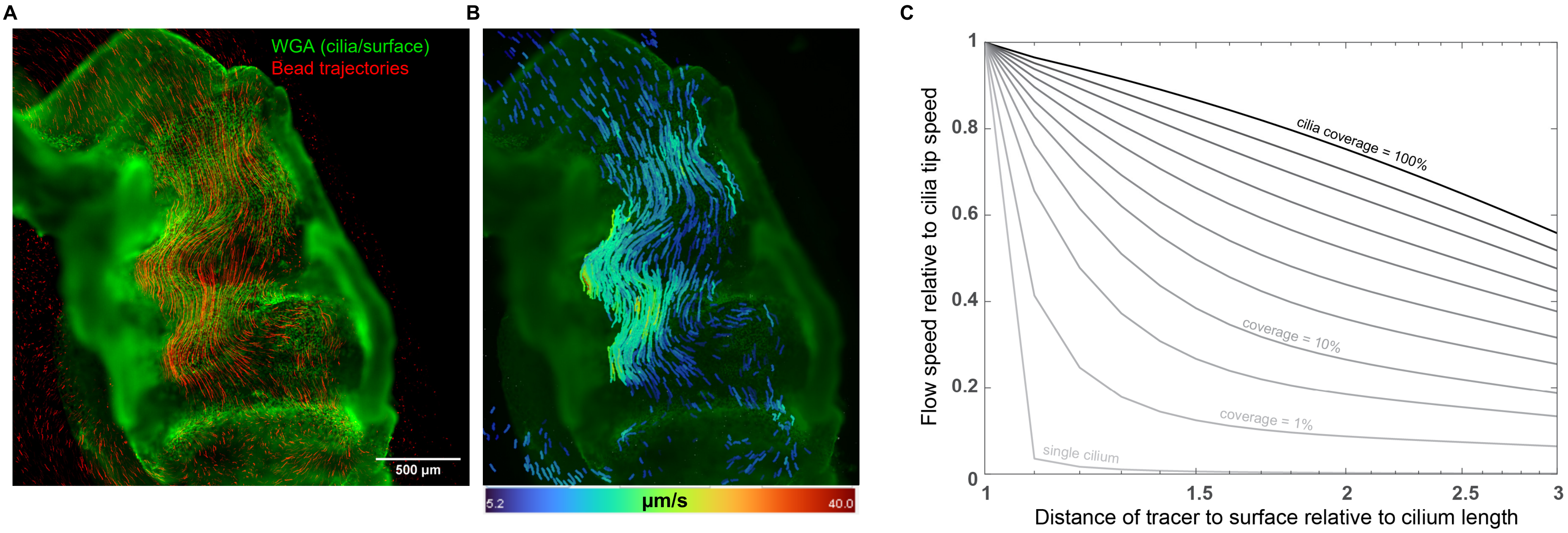

**Supplemental Figure 9: Surface curvature reduces accuracy of clearance measurements.**
**A** Example of live bead clearance (red trajectories) in longitudinal section of rat bronchial tube. The ciliated surface (green) is warped due to elastic recoil and cartilage rings. Such warp means that the recorded bead trajectories become contorted and may reflect different distances from the ciliated surfaces.
**B** Variable flow speed measured from tracer displacement shows that speed is highest where the tracers pass directly over a fold, i.e. are closest to the surface.
**C** Computational data showing that increasing distance of tracer from surface dramatically lowers the relative flow speed reached by tracers. This effect becomes more pronounced with decreasing cilia coverage. For a typical rat bronchus with ca. 50% ciliation and a cilia length of 5 µm, a tracer that is 10 µm away from the cilia tip experiences at most 60% of the maximal flow speed near the tip.

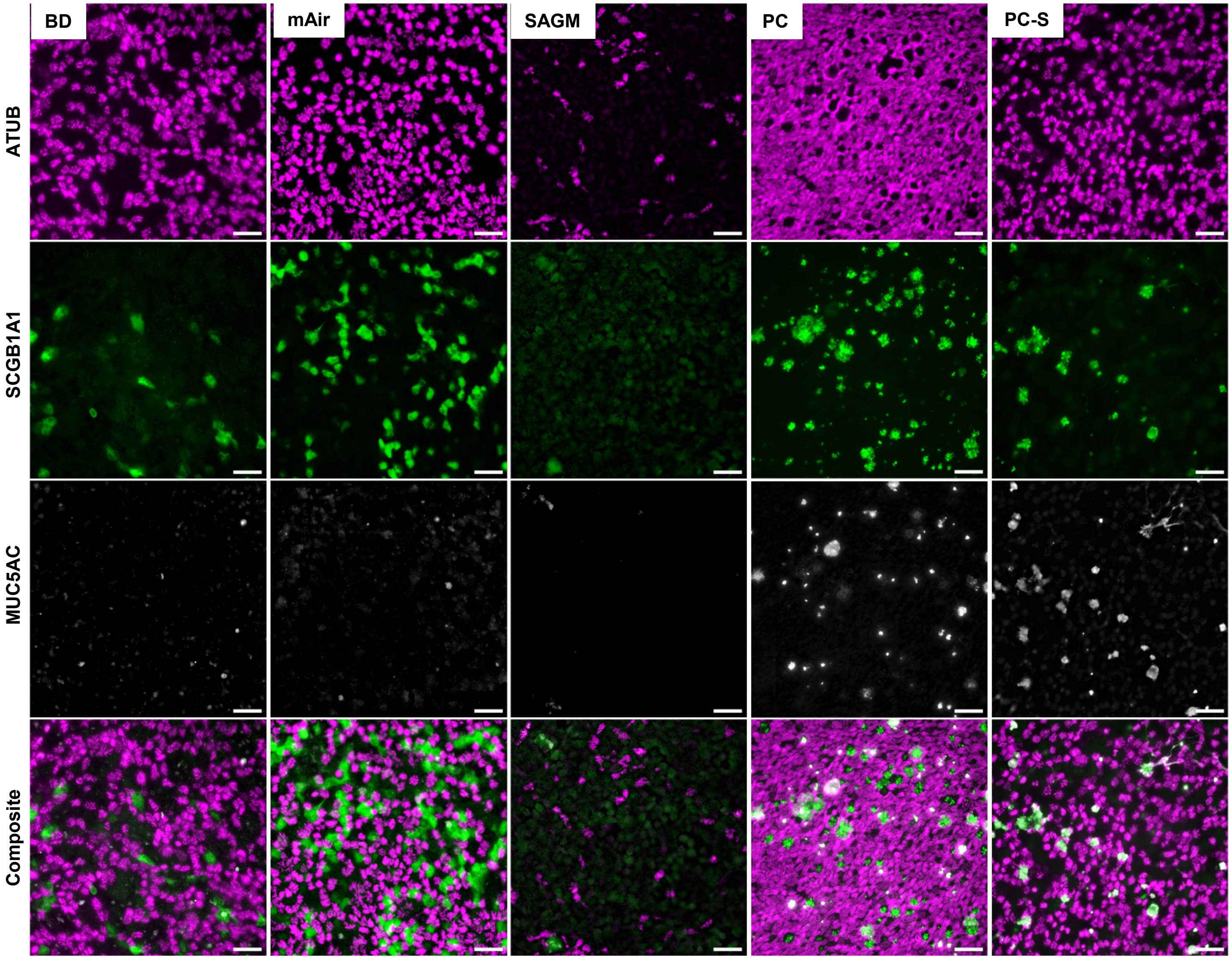

**Supplemental Figure 10: Cell culture media impacts cell type composition in differentiated airway epithelial cultures.** Same conditions as in Figure 4a using a different cell donor. Representative images from day 28 of differentiation at ALI. Scalebar, 40 µm.

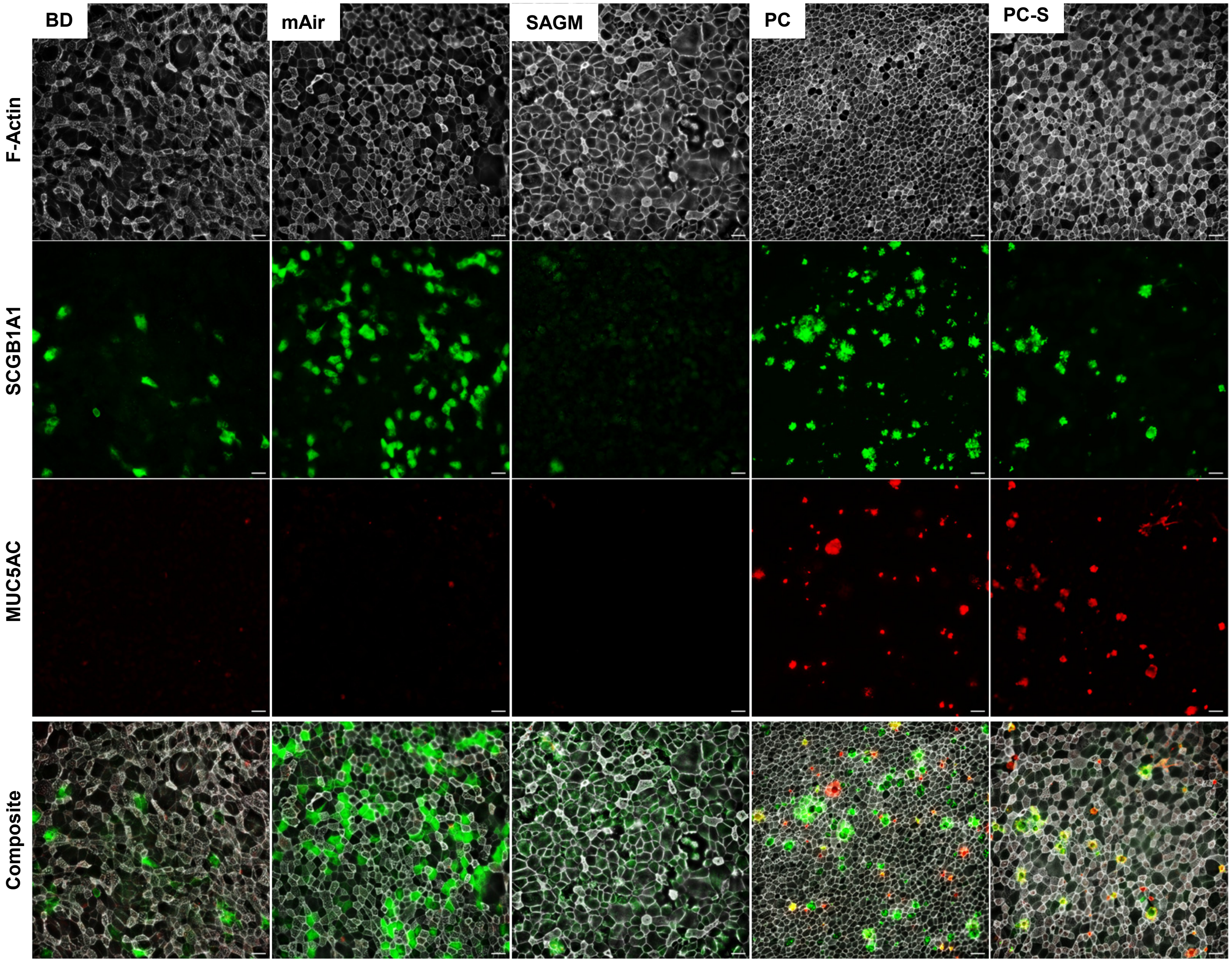

**Supplemental Figure 11: Cell culture media impacts cellular size and shape.** Luminal cell outlines used for quantifying relative cell proportions are visualized by phalloidin staining of F-actin. Same fields of view as in Figure S9. Scalebar, 20 μm.

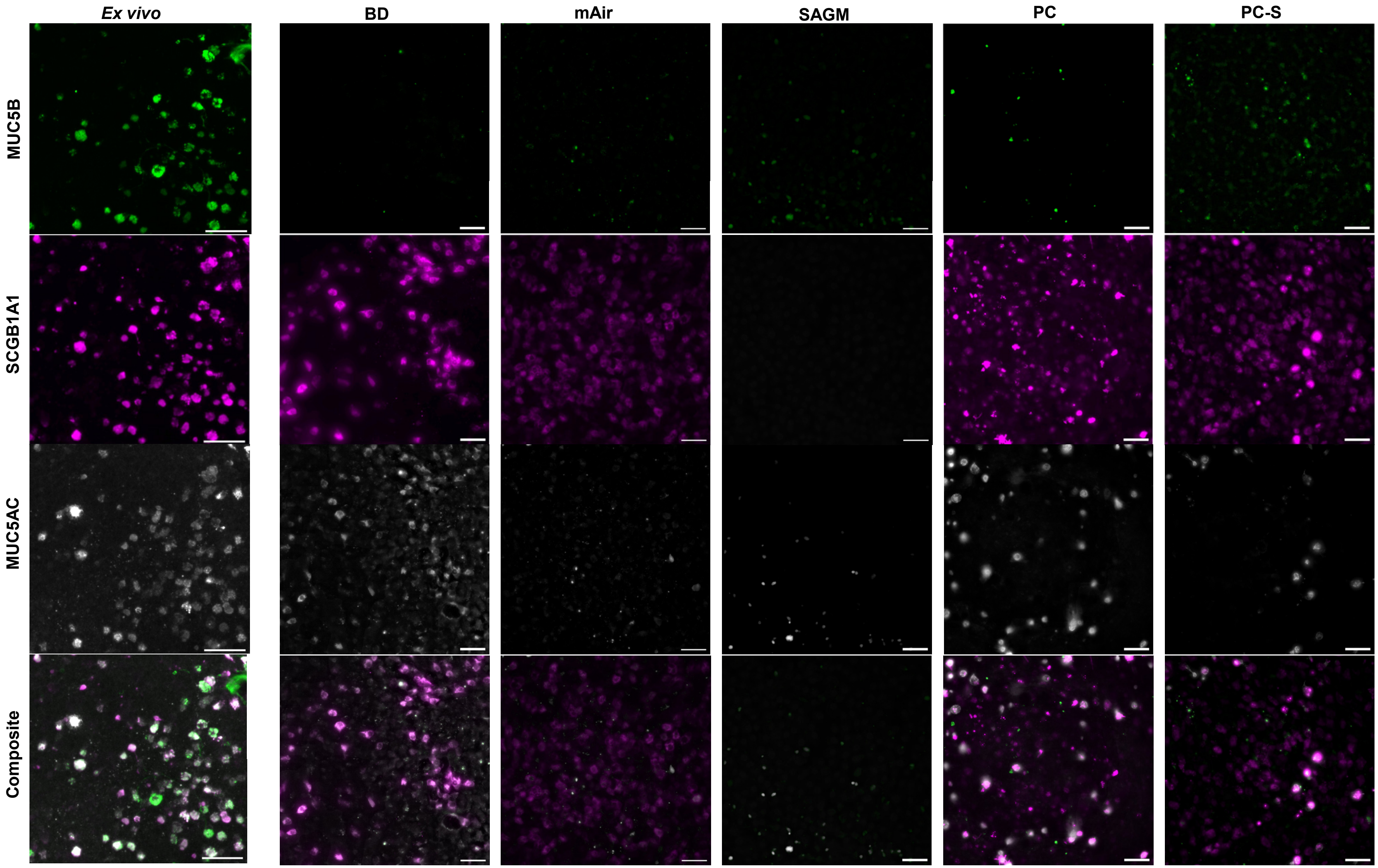

**Supplemental Figure 12: Secretory cell type composition *ex vivo* and *in vitro*.** IF staining of secretory cell markers MUCAC, MUC5B and SCGB1A1. *Ex vivo*: Example images from 1 donor (2924) taken from the ventral wall ; *In vitro*: Example images from 1 donor (6288) at day 28 of differentiation at ALI. Scalebar, 40 μm.

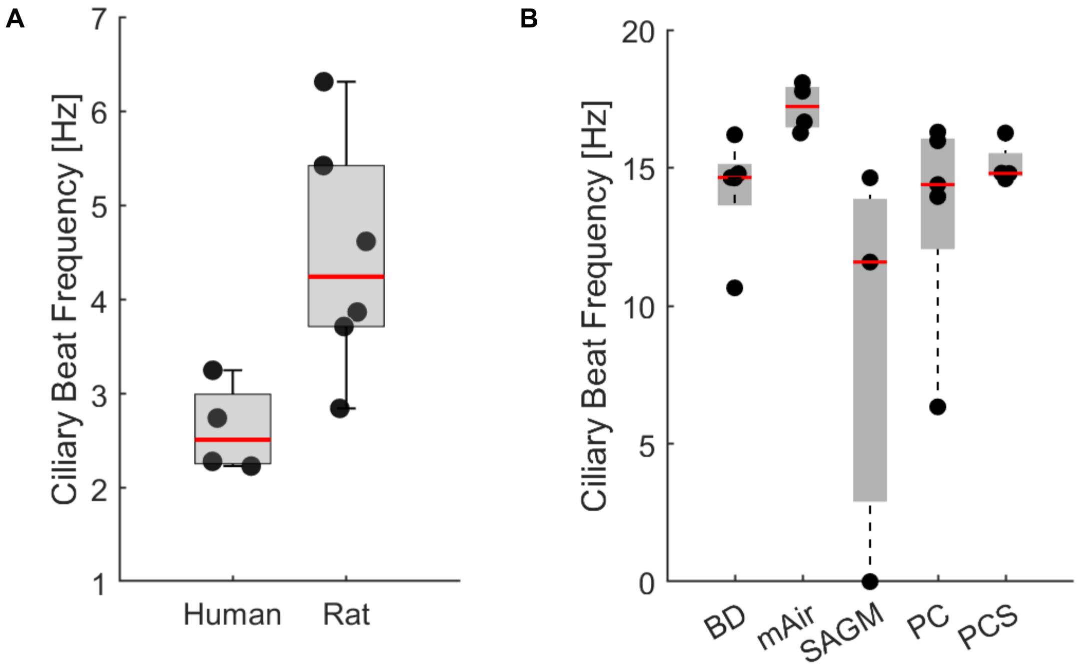

**Supplemental Figure 13: Ciliary beat frequency comparison.** Mean ciliary beat frequencies of **A** *ex vivo* airway samples from rat (BG0-1, N=6) and human (BG0-6, N=4) stored at 4°C and recorded at 18°C and **B** in vitro human airway epithelial cultures maintained and recorded at 37°C (N=4-7; some donors did not differentiate in SAGM). Solid dots correspond to mean values of individual donors. On each boxplot, the central mark indicates the median, and the bottom and top edges of the box indicate the 25th and 75th percentiles, respectively. The whiskers extend to the most extreme data points not considered outliers (ca. +/–2.7 times standard deviation). Source data for panels A and B are provided as a Source Data file.

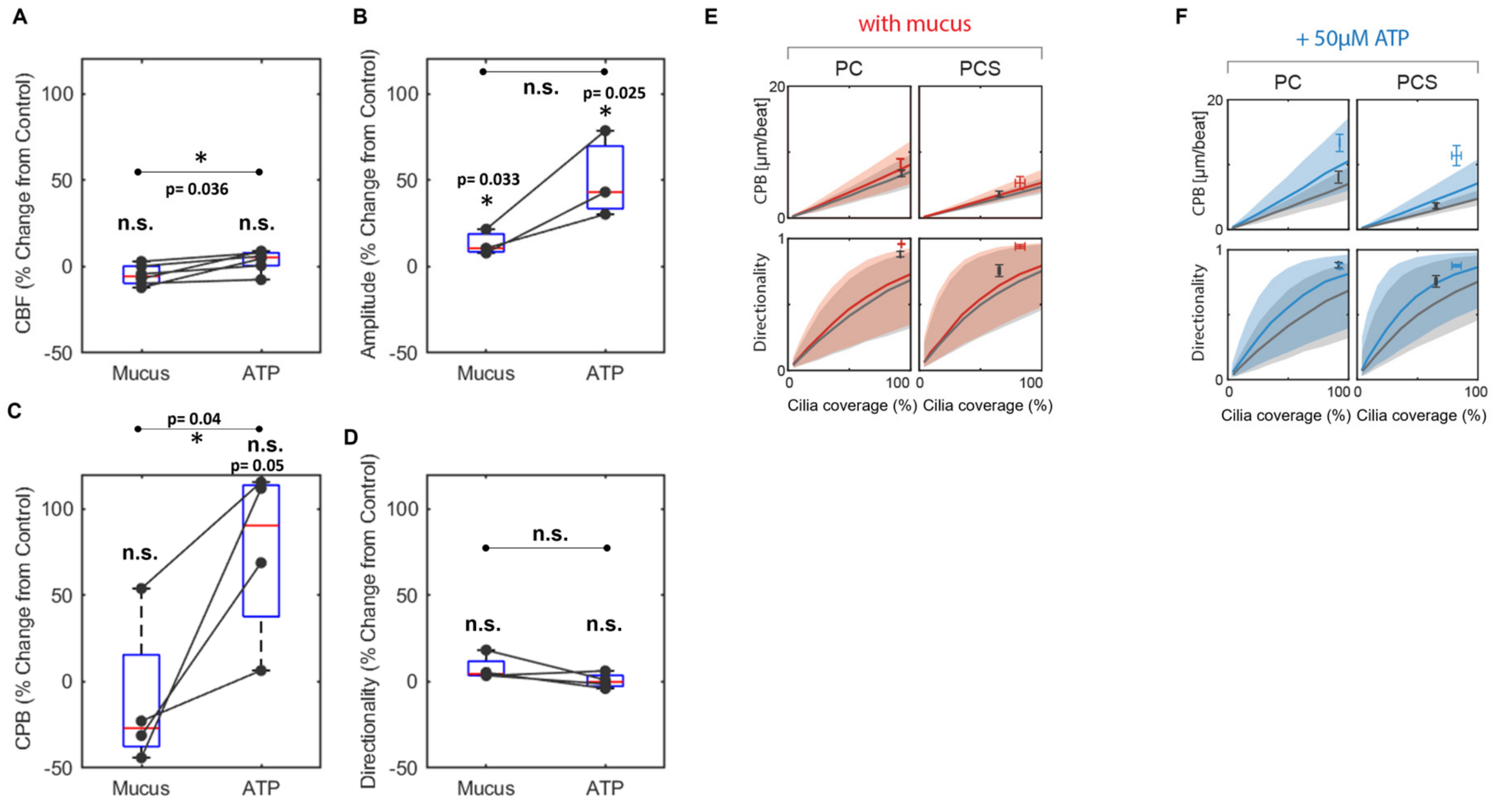

**Supplemental Figure 14: Impact of mucus and ATP on ciliary beat and MCC.** Compared to control (washed samples), the percent change of multiple readouts was measured in unwashed (Mucus) and washed and ATP-treated (ATP) samples: **A** CBF, **B** Ciliary beat amplitude, **C** Clearance per beat, and **D** Clearance directionality. Dots represent averages values recorded in different donors (N=3 total) in either PC or PC-S medium. Averages were formed over 3 inserts with 1-6 FOVs each. Lines connect measurements performed in same donor and medium. Statistical significance is indicated for unpaired t-test between Mucus and control, ATP and control, and between Mucus and ATP. For C and D, only samples with comparable cilia coverage were included as both metrics are sensitive to cilia coverage. On each boxplot, the central mark indicates the median, and the bottom and top edges of the box indicate the 25th and 75th percentiles, respectively. The whiskers extend to the most extreme data points not considered outliers (ca. +/–2.7 times standard deviation).Source data for panels A-D are provided as a Source Data file.
**E** Plot of CPB and clearance directionality against cilia coverage measured in control (black) and mucus-containing (red) samples in PC or PC-S medium (mean±SEM; N=3 per medium) , compared to model predictions that take into account the changes in ciliary beat amplitude in mucus conditions. Solid lines: average prediction; shaded area: area of uncertainty due to variability in input parameters. **F** Plot of CPB and clearance directionality against cilia coverage measured in control (black) and ATP-stimulated (blue) in PC or PC-S medium (mean±SEM; N=3 per medium), compared to model predictions that take into account the changes in ciliary beat amplitude in ATP conditions. Solid lines: average prediction; shaded area: area of uncertainty due to variability in input parameters.

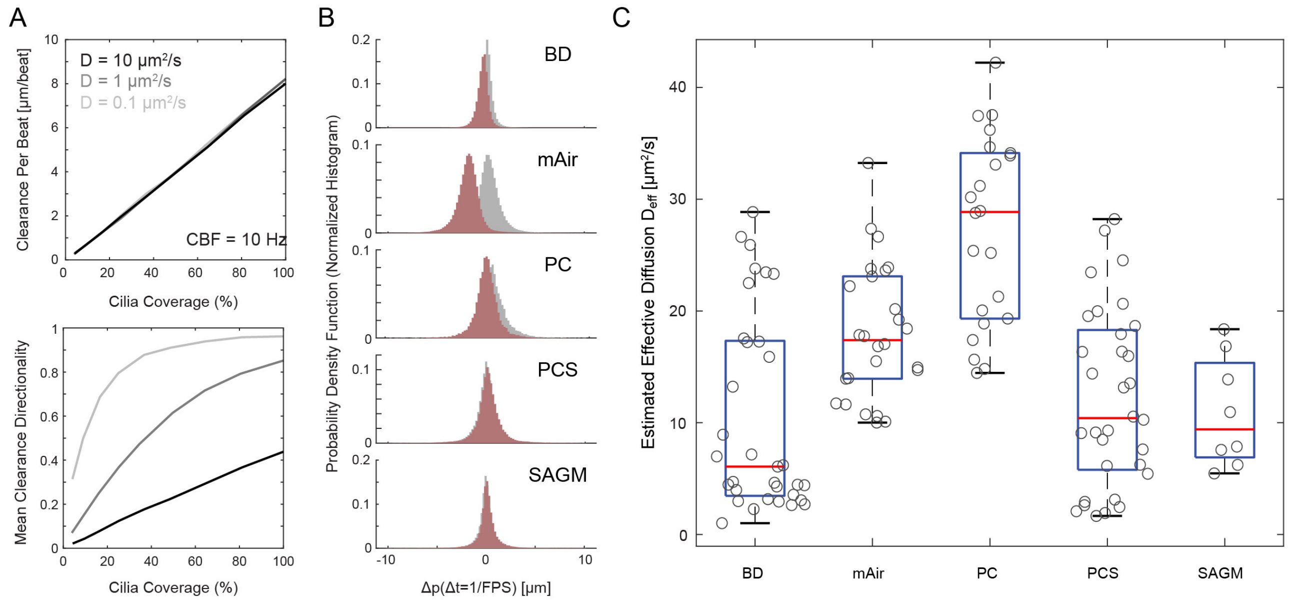

**Supplemental Figure 15: Estimated effective diffusion constant for in vitro cultures.**

**A**. Different values of diffusion constant only changes mean clearance directionality without significant impact on clearance per beat. Here we fix cilia height and beat amplitude to be 10 μm, λ= 30 μm m, CBF = 10 Hz, OOP = 0.6, and COP = 0.53.

**B**. Representative particle displacement distributions at the minimum possible time lag Δt = 1/FPS, where FPS are the frames per seconds of the associated video recording. Here, we verify that locally, the recorded particle trajectories have a displacement per time step that is distributed normally. An effective diffusion coefficient can thus be calculated based on the variance of this distribution. Grey, displacement in x-coordinate; red, displacement in y-coordinate.

**C**. All estimated effective diffusion constants $D_{eff}$ grouped by different culture medium conditions. Here, each circle represents results from a single field-of-view. On each box, the central mark indicates the median, and the bottom and top edges of the box indicate the 25th and 75th percentiles, respectively. The whiskers extend to the most extreme data points not considered outliers (ca. +/–2.7 times standard deviation).

Source data for panels B and C are provided as a Source Data file.

.

### Supplementary Discussion

### 1 Physics-based model

#### 1.1 Cell-level ciliary input parameters.

The fundamental solutions of Stokes' equation are frequently used to model fluid dynamics generated by cilia [8–14]. Computationally, the force generated by a cilium is modeled by either distributed point forces along the cilium, or by a single, coarse-grained point force that represent the net force of a cilium at a given time instance during its beat cycle (e.g., the rotor or rower model of cilia). As the goal of our study is to map the structure-function relationship for ciliated surfaces that contain hundreds to thousands of cells, we coarse-grain the motion of the hundreds of cilia per multiciliated cell in both space and time into a single point force. This approach assumes that (i) mucociliary clearance is primarily performed by the 'envelope' formed by the power strokes of dense patches of straight cilia, and (ii) the ciliary beat pattern and their state of coordination remain unchanged during our time-scale of interest. While this approach cannot resolve the flow and coordination of individual cilia in models presented elsewhere [10, 13–16], it is straightforward to implement, suitable for large number of ciliated cells, and directly takes into account of the wall-screening effects due to finite cilia length comparing to the slip-boundary-velocity approach [17]. Under these assumptions, the linearity of Stokes' flow (Reynolds number ≪ 1) implies that we can approximate the net effect of all cilium of a cell as a single static effective force that sits at $z = h$, one cilia length $h$ above the no-slip cell surface, and points in the direction of the average power stroke direction of the cell, assumed to be along the $x-$axis without loss of generality. The force magnitude will be the product of the ciliary beat amplitude $A$, ciliary beat frequency $f$, and a drag coefficient $\zeta$; See schematics shown in Supplementary Fig. 7A.

We do not consider the double confinement or mucus film effects discussed in [17], since we intend to compare tracer particle motions recorded near the distal tip of cilia and sufficiently far from other confinement boundaries such as coverslip and air-liquid interface, i.e., the height of the liquid film $H \gg h$. Experimentally, we also perform several washes to remove mucus from the cell surface before measurements of cilia driven flow and we choose appropriately sized field-of-views such that no recirculation effects are observable.

Numerically, we compute the flow field produced by the effective point forces using the regularized Stokeslet algorithm [18]. This solves the forced Stokes' equation

$$\begin{aligned} \mu\Delta\mathbf{v} - \nabla p + \zeta A f \delta_\epsilon(\mathbf{x} - h\hat{\mathbf{z}})\hat{\mathbf{x}} = 0, \\ \nabla \cdot \mathbf{v} = 0, \end{aligned} \tag{1}$$

with the no-slip boundary condition $\mathbf{v}(z = 0) = 0$. Here, $\mu$ is the viscosity, $\mathbf{v}(\mathbf{x})$ the velocity of the fluid at point $\mathbf{x}$, $p$ the pressure, and $\delta_\epsilon(\mathbf{x} - h\hat{\mathbf{z}})$ a blob function centered around $[0, 0, h]^\mathsf{T}$ with its spread determined by the regularization parameter $\epsilon$ and integrates to unity [18]. As $\zeta$ and $\epsilon$ are coarse-grained, effective parameters that cannot be measured, we need to determine their size based on physical insights and then validate our choices by confirming that our choices reproduce the structure-function relationship revealed by the experimental data. The regularization parameter $\epsilon$ can be interpreted as the effective hydrodynamic radius of the cilia bundle, which should be larger than the radius of a single cilium ($\sim 200$ nm) but smaller than the size of a single ciliated cell ($\sim 10$ µm). For simplicity, we set $\epsilon = 1$ µm. This implies that the induced flow field is accurate at the scale of cell sizes without introducing spurious singular peaks that would be smeared out by the asynchronous motion of individual cilium in reality. We also confirmed by numerical experiments that using different values of $\epsilon$ that satisfy our order of magnitude limit do not dramatically change the results of model output. Since we measure ciliary beat amplitude $A$ as the maximum distance the tip of the cilia travel during one beat period, the maximum speed induced by cilium motion should be of the order $Af$. Thus, we set $\zeta = \mu$ to ensure that $\max(|\mathbf{v}|) \sim O(Af)$ given our choice of $\epsilon$. Albeit based on simplifying estimates, our chosen $\zeta$ and $\epsilon$ produces output metrics that closely match the *ex vivo* human benchmarks (Fig. 3C of main text). Since different $\zeta$ will introduce a simple linear scaling of induced velocity, this match also means that our choices can be seen as the correct fitting parameters for our experimentally observations.

This leaves the cell-level ciliary input parameter to be (1) cilia length $h$, (2) ciliary beat amplitude $A$. Throughout our computation, we fix fluid viscosity $\mu$ to be that of water, and non-dimensionalize time and length using cilia beat frequency $f$ and the assumed size of a ciliated cell (10 µm).

#### 1.2 Tissue-level ciliary input parameters

We derive the ciliated cell distribution based on a cilia coverage fraction $C \in (0, 1)$ and a crystalline order parameter as defined in [17], where the crystalline order parameter (crystalline OP) is computed based on the distribution of ciliation gap size $\lambda$ between each ciliated patch as $1 - \sqrt{2} \cdot \text{std}(\lambda)/\text{avg}(\lambda)$; see Supplementary Fig. 7A. Here the cilia coverage (% ciliated cells), the average and standard deviation (std) of $\lambda$ can be mapped one-to-one with the structural measurements. The periodic ciliated patches are generated following our previously reported algorithm [14, SI section 6], except that the rotor heterogeneity is used to express the crystalline disorder of ciliated cells. Specifically, we compute the ciliated cell positions by designing a Boolean function of $(x, y)$ that returns true if and only if a ciliated cell is present at that location. We start from a 1D triangular-wave function $\mathcal{T}(x)$, parameterized to vary periodically from $-1$ to $1$ with a period of 1, and apex at $x = 0.5$ in each wavelength

$$\mathcal{T}(x) = \begin{cases} 2 \bmod (x+1, 2) - 1 & \bmod (x+1, 2) \leq 1 \\ 1 - 2(\bmod(x+1, 2) - 1) & \bmod (x+1, 2) > 1 \end{cases}, \tag{2}$$

The piecewise linearity of the triangular-wave allows us to obtain an arbitrary 1D coverage ratio $L_C$ by setting a function to return true if and only if $\mathcal{T}(x)$ is above $1 - 2L_C$. We generalize this one-dimensional construction to two-dimensions by combining two 1D functions using the unit norm separable function $f(x, y) = \min(f(x), f(y))$. Namely, we get

$$\mathcal{T}_{2\text{D}}(x, y) = \min \left[\mathcal{T}\left(x/\text{avg}(\lambda) + \eta_x\right), \mathcal{T}\left(y/\text{avg}(\lambda) + \eta_y\right)\right], \tag{3}$$

where we re-parameterize $x$ and $y$ by the average ciliation gap size avg($\lambda$) and the ciliated cells are displaced by a Gaussian noise $\eta_x, \eta_y \sim \mathcal{N}(0, \text{std}(\lambda))$. To ensure that the area fraction of the periodic domain covered by cilia is $C$, we set $L_C = \sqrt{C}$ and test if $\mathcal{T}_{2\text{D}}(x, y)$ is above $1 - 2L_C$ just like in the one-dimensional case.

The beat direction angle $\theta_i$ of each ciliated cell $i$ are sampled from a wrapped normal distribution with a zero mean (cilia beat globally towards the positive $x-$axis direction on average). The variance of this distribution is specified by the ciliary beat order parameter (ciliary OP; orientation order parameter or the second circular moment of the wrapped normal distribution), and the covariance is determined based on the relative distance between each ciliated cell. In other words, $\theta_i$ is drawn from $\bmod(\mathcal{N}(0, M_{ij}), 2\pi)$, where the covariance matrix $M_{ij} = \sigma \exp(-d_{ij}/R_o)$. Here $\sigma$ is the variance associated to the given ciliary beat OP, $d_{ij}$ represents the distance between cell $i$ and $j$, and $R_o$ is the characteristic decoherence length for each angle. The numerical value of $\sigma$ is calculated by inverting the expression for ciliary beat OP $\exp(-2\sigma^2)$. We set $R_o$ to be equal to the ciliation gap size $\lambda$ based on the observation that ciliated cell in a same patch tend to have highly correlated effective stroke angles; see [17] and our own data (Fig. 2d, left).

Together with the patch heterogeneity parameters, we have the following tissue-level input parameters (1) % ciliated cells, (2) patch wavelength, (3) crystalline order parameter, and (4) orientation order parameter.

#### 1.3 Model output

We simulate ciliary flow on top of a square grid of 51 x 51 cells each 10 µm in size, doubly periodic in both $x-$ and $y-$axis, where $x-$axis is defined as the clearance direction. Cell centers are slightly perturbed from the exact grid points in Monte-Carlo simulations; for visual reference, cell boundaries are drawn based on the Voronoi diagram of the perturbed cell centers (Supplementary Supplementary Fig. 7B). Theoretically, we need to compute the fluid velocity by summing the hydrodynamic interaction of all force monopoles spanning a doubly-infinite image system that tiles the entire 2D plane [19, 20]. This doubly-infinite sum is conditionally-convergent because of the quadratic decay of the Blake-Oseen tensor in semi-infinite domains [21, 22]. Since our interests lie in the coarse-grained ciliary flow characteristics and, in reality, the ciliated tissue do not extend to infinity, we relax the requirement of computing an infinite lattice by considering only the hydrodynamic interactions within one periodic image domain. Based on our chosen grid size, this means that we will be truncating the influence of ciliated cell more than 250 µm away from any given point of interest. The quadratic decay implies that this introduces only a small error in the computed flow velocity.

We compute our ciliary flow characteristics of interest by running Brownian dynamics simulations for particles injected near the distal tips of cilia. Tracers are subject to cilia-driven flow due to forces associated to each ciliated cell and random fluctuations governed by a diffusivity constant $D$; see §1.5. Time evolution of $500$ particles are computed for $500$ beat periods, following the Langevin equation

$$\frac{\mathrm{d}\mathbf{r}}{\mathrm{d}t} = \mathbf{v}(\mathbf{r}) + \sqrt{2D}\eta(t), \tag{4}$$

where $\mathbf{r}$ is the position of a simulated particle, $\mathbf{v}(\mathbf{r})$ the cilia-driven flow, and $\eta(t)$ a vector of standard Wiener process defined by the correlation functions $\langle \eta_i(t) \rangle = 0$ and $\langle \eta_i(t)\eta_j(t') \rangle = \delta(t-t')\delta_{ij}$, i.e., the added noise is not correlated between different time steps and $x$-, $y$- components. We initialize the particles uniformly in $x-$ and $y-$ directions but fix their height at the tip of cilia ($z = h$). Equations are numerically integrated with an Euler-Maruyama scheme with one time-step equal to 5% of the beat period. This value is used to ensure the computed trajectories achieve convergence. The periodic boundaries in both $x-$ and $y-$directions is enforced by always considering the hydrodynamic influence within the fundamental domain centered around the particle of interest.

The quantitative output of our simulation is clearance per beat (CPB), i.e., the average distance that particles translate in the $x-$direction per ciliary beat period, and clearance directionality, i.e., the ratio between the magnitude of the average flow field vector $\mathbf{v}$ and the average of the magnitude of flow field vector $\mathbf{v}$ for boxes of side length equal to 80 µm; also see Supplementary Fig. 7A. To obtain the associated flow vector field $\mathbf{v}$, we follow the same particle trajectory velocimetry process used to post-process experimentally measured tracks, where we average the velocity of each simulated Brownian track into a rectangular Eulerian grid. In simulation we fix the grid resolution at $4$ points per µm, similar to the experimental pixel density.

#### 1.4 Controlled variations

To visualize how tissue-level parameters affect our model output, we present case studies where either cilia coverage, crystalline order parameter, and/or orientation order parameter are at exaggerated values in Supplementary Supplementary Fig. 7B. Here we cropped the field of view to be centered around a 15 x 15 cell section for legibility. While crystalline order parameter significantly changes how ciliated cells are distributed (darker cells with labeled cilia beat direction), it does not significantly impact the characteristics of the tracer trajectories. Both lower cilia coverage and orientation order parameters significantly reduce the clearance distance and directionality of the tissue.

In Supplementary Fig. 8, we examine how CPB and clearance directionality changes quantitatively for different (A) cilia length, (B) cilia beat amplitude, (C) cilia beat order parameter and (D) crystalline order parameter as functions of cilia coverage. In general, CPB scales linearly with respect to % ciliated cells, while directionality measures shows an exponential plateau as cilia coverage approach 100%. This linear increase of CPB with respect to cilia coverage is because our hydrodynamic model follows the Stokes' flow regime, and without contribution of boundary terms, net transport effects are simply additive due to the linearity of the Stokes' equation. This contrasts with the findings of [17], where local recirculation is possible as only a very thin film of fluid is assumed to be transported on top of the tissue. Tracers that move due to isotropic Brownian motion will exhibit zero directionality in the infinite-time limit, since Brownian motion does not have a preferred direction of travel. In contrast, a point directly on top of a cilium should in principle exhibit very high directionality, close to 1 in the infinite-time limit. On top of unciliated area, tracer directionality will be strongly influenced by nearby ciliated cells as the hydrodynamic effect of a single ciliated spot (Stokeslet) decays slowly at a finite rate. Therefore, as long as the tissue surface contains enough ciliated spots such that the streamline connects, the mean clearance directionality will be far away from 0 throughout. This slow-decay effect causes clearance directionality to increase steeply with cilia coverage increase initially, and plateau at full ciliation to a fixed value, which is determined by factors like cilia beat order parameter or beat amplitude. For biologically relevant ranges of cilia length (4 to 10 µm) and ciliary beat amplitude (5 to 15 µm), CPB scales roughly linearly. The clearance directionality seems to have a sublinear growth with respect to cilia length, and increases linearly with respect to beat amplitude. If the orientation order parameter drops below certain values (cilia are significantly misaligned), we see that CPB and directionality are severely reduced. Changing the crystalline order parameter has little to no effect on either clearance measures; this is because they are primarily measuring the characteristics of flow averaged over the entire field of view and are not sensitive to local variation. Thus we opt to use only the mean patch wavelength value and crystalline order parameter when trying to predict clearance metric from measured input parameters (Fig. 3e and 5c). In Supplementary Fig. 9, we show that if we change the distance of the tracer particles to the cilia tips, the relative flow speed dramatically decreases. This effect becomes less pronounced when cilia coverage is high since nearby ciliated patches slow down the decay of hydrodynamic interaction [23].

#### 1.5 Effective diffusivity

Since the size of tracer particles are fixed to be $1$ µm in diameter (both experimentally and in simulation), and we ignore the effects of viscosity variation (experimentally mucus was removed from samples via multiple washes), the baseline thermal diffusivity $D$ of the simulated particles in water can be easily calculated using the operating temperature $T$ via the formula $D = k_B T/(6\pi\mu r)$ where $\mu$ is the viscosity of water at temperature $T$ and $r$ the particle radius. Ex vivo experiments were conducted at 20°C, giving $D \approx 0.4$ µm$^2$/s. Since the particle tracks were recorded at a limited frame rate due to experimental restrictions, we assume that thermal diffusion is the dominant source of noise and use this value of $D$ for our baseline model results shown in Fig. 3 and 6. The close match between the measured clearance directionalites and those predicted by our model serves as a sanity check of this choice. Note that since only uncorrelated white noise is considered, the value of $D$ has no impact to the predicted clearance distance per beat; see Supplementary Fig. 15A.

For *in vitro* experiments conducted at 37°C, a similar calculation shows that the thermal diffusivity $D \approx 0.7$ µm$^2$/s. However, this value of D together with the measured tissue- and cell- level structural parameters predict significantly higher clearance directionalities than the measured values. We believe this is caused by an additional source of noise, possibly due to immature ciliary beat patterns. Ciliary beat patterns can take longer to fully mature *in vitro* than the 28 days used in our study [24]. There could also be additional activity of cilia *in vitro* since these samples were never refrigerated during transport like their*ex vivo* counterparts. Therefore, we choose to compensate this undetermined source of noise by estimating an effective diffusion constant $D_{\text{eff}}$ based on the measured particle trajectories.

Specifically, for every time series of particle positions $\mathbf{p}(t)$ in $x-$ and $y-$coordinates, we compute the displacements $\Delta\mathbf{p}(\Delta t) = \mathbf{p}(t + \Delta t) - \mathbf{p}(t)$ for all time lags $\Delta t$ smaller than a single cilia beat period. Then for every such time lag, we can compute the covariance matrix $M$ for $\Delta\mathbf{p}$, and obtain an estimate for the effective diffusion coefficient $D_{\text{eff}}(\Delta t)$ from its diagonal entries via $D_{\text{eff}} = \mathrm{Trace}(M)/(4\Delta t)$. This is possible because cilia are expected to generate bulk transport at time scales at least as large as a single beat cycle due to differential drag between effective and recovery strokes. Therefore, at small enough time lags, we expect each component of $\Delta\mathbf{p}$ to be distributed like values drawn from independent normal distributions with possibly a small mean values accounting for bulk transport; for representative histograms, see Supplementary Fig. 15B. We verify this by hand and reject samples where $\Delta p$ at smallest recorded $\Delta t$ is clearly not normally distributed.

We show the final estimated $D_{\text{eff}}$ grouped by the different culture medium used in Supplementary Fig. 15C. Here each circle represents a result calculated using a single tracking movie / field-of-view. Indeed, we see values significantly larger than pure thermal diffusivity, and there exists a condition-specific variation. In Fig. 5, we use the minimum, maximum (black whiskers in Supplementary Fig. 15C), and median value (red lines in Supplementary Fig. 15C) of our estimated $D_{\text{eff}}$ when testing our model.

### 2 Limitations of experimental and computational data

#### 2.1 Experimental

The ciliary beat and clearance recordings were taken at room temperature ( ca. 20 ℃) in the case of *ex vivo* tissues, and at 37 ℃ in the case of *in vitro* tissues due to experimental constraints. Resulting differences in ciliary beat frequency was accounted for by normalizing clearance speed to clearance per beat. Changes in diffusion coefficients were implemented in our computational models. Nonetheless, other parameters relevant to ciliary beat could be affected by temperature.

Due to the limited availability of healthy human lungs for scientific research, the *ex vivo* data from 12 out of a total of 14 human donors (see Supplementary Table 1) are used in our model and may not represent the full span of natural variability. Some of the analyzed fixed bronchial rings were peritumoral tissue. The fact that ciliation levels were nonetheless robustly high provides confidence that this is a key feature of the human airways. We also observed the expected gradients in the proximal-distal distribution of goblet and club cells, providing additional confidence in the samples. Therefore, the data presented here serve as a guide until additional studies refine the picture.

We chose to standardize our clearance directionality measurement by averaging values with a fixed window size of 80µm, much greater than the typical ciliated patch wavelength (∼30µm). This value is chosen because in the limit of homogeneously distributed ciliated patch and tracer particles, directionality should approach a constant for a sufficiently large window. However, if a given field-of-view contains only a few ciliated patches, any cilia-driven transport far away from cilia is likely slow, multi-directional and difficult to distinguish from spurious tracks produced by particles stuck to the cell surface. As a result, trajectories from non-ciliated areas may be filtered out by the tracking algorithm and are then missing from the data set, biasing the results to tracks near ciliated cells that exhibit higher directionality; see points with low cilia coverage for *in vitro* culture in BD and mAir media of Fig. 5c. Therefore additional optimization for sparsely ciliated tissue is warranted in the future.

#### 2.2 Computational

Although based on sound physical principles, we did not separately verify the assumption that period-averaged flow due to a single multi-ciliated cell can be modeled by a single static Stokeslet. This is because it is difficult to isolate and control individual ciliated cells in human airway tissues without significant perturbation or destruction of its surrounding tissue, in contrast to what is possible for single-celled swimmers [11], or tissue where cilia naturally form easily separable bundles [23]. Nevertheless, the fact that we were able to match tissue-level CPB and clearance directionality to the reported accuracy strongly suggests that we are capturing the dominant underlying physical mechanism.

Our model precludes the possibility of bimodal or multimodal beat angle distribution, which could be true for *in vitro* cell cultures where cilia beat have not yet reach global alignment. While it is possible to extract exact angle distributions for specific field-of views, flow tracking cannot be performed on the exact same field-of-views simultaneously, and therefore we do not think that using the exact angle distributions would significantly improve CPB and clearance directionality predictions compared to our statistical approach.

We took an empirical approach when modeling the additional diffusive noise associated with *in vitro* cell cultures. The media-dependency of observed effective diffusivity (Supplementary Fig. 15C) suggests that the noise could have a biological origin associated with different degrees of ciliary beat maturation, which warrants exploration in future studies. In the *ex vivo* samples, the frame rates of the bead tracking recordings were not sufficiently high to estimate the diffusion constants; however, using diffusion coefficients purely based on the ambient temperature resulted in convincing CPB and clearance directionality predictions, therefore, no additional analysis was required.

### 3 User Guide for `simpleMCC.m`

1. (line 28) Modify string variables `dataFolder`, `figFolder`, `movFolder` according to the local computing environment. They should point to the directory for output data, output figure, and output movies, respectively.

2. (line 32) Set boolean variables `makeMovie`, `saveMovie`, `saveData` to the desired value. By default, movie will be generated on screen (`makeMovie=true`), AVI movie will be compiled (`saveMovie=true`), and no data or figure will be saved (`saveData=false`).

3. (line 42) Choose simulation scenario by (un)commenting the appropriate `caseList = {'...'};` and `modList = {'...'};` lines. By default, an example simulation will run and results shown in Supplementary Fig. 7B will be simulated on screen.

   (a) (line 45) To regenerate validation simulations using on kinematic input parameters derived from *ex vivo* samples (Fig. 3e), uncomment/modify this line. Case name starts with `human` (human) or `rat` (rat) and ends in `avg`, `min`, or `max` for the mean, lower, or upper bound of the model predictions.

   (b) (line 48) To regenerate simulations based on kinematic parameters derived from *in vitro* samples (Fig. 5c top panels), uncomment/modify line 48-50. Case name prefix specify differentiation medium type (`bd`, `mair`, `pc`, `pcs`, `sagm`, see Methods section "Source and generation of human primary airway epithelial cell cultures"). Variable suffix `avg`, `min` or `max` indicates running the mean, lower, or upper bound of the model predictions. (A special case name `pc_D37` can be added to show the case of using mean kinematics derived from PneumaCult medium using diffusion coefficient based on passive tracers in 37°C water). To generate cases used for Fig. 5c bottom bar charts, uncomment line 63 to add the alteration scenario where select input parameters (modList suffix `h`, `ap`, `oop`, `diff` indicates cilia length, beat amplitude, beat OP, or diffusion coefficient, respectively) are replaced by their counterparts from human ex vivo benchmarks. (Another set of cases where input parameters except the specified ones are replaced by those from human ex vivo benchmarks are available by uncommenting line 64).

   (c) (line 57) To regenerate parametric studies shown in Supplementary Fig. 8 and 15, uncomment line 57 or 58. Case name `hVar`, `aVar`, `kVar`, `pVar`, or `diffVar` stands for variation in cilia length, beat amplitude, beat order parameter, crystalline order, or effective diffusion coefficient, respectively.

4. (line 80) Characteristic scales assumed by the simulation. The numerical method runs assuming the fluid viscosity, cilia beat frequency and cell spacing to be of unit value. Together these fixes the force, time, and length scale of the simulation.

   (a) (line 138) Dimensionless diffusion coefficient for *ex vivo* validation simulations are calculated from average cell spacing (L0=$10\mu$m), the viscosity of water, tracer particle radius (r=$0.5\mu$m), Boltzmann's constant, and measured cilia beat frequency (`CBF`).

   (b) (line 145) Dimensionless diffusion coefficient for *in vitro* simulations are calculated from empirical data (See Supplementary Fig. 15 and discussion) combined with cilia beat frequency measurements and cell size estimates.

5. (line 168) Kinematic input parameters derived from *ex vivo* and *in vitro* samples (See 1.1). These variables has suffix indicating cilia length (`h`), cilia beat amplitude (`ap`), cilia beat angle order parameter (`oop`), ciliation crystalline order parameter (`cop`), and prefix indicating data source (`human` for *ex vivo* human, `rat` for *ex vivo* rat, and abbreviations for various *in vitro* differentiation medium as in 3b).

   (a) (line 222) Convert cilia beat angle order parameter to input parameter for wrapped normal distribution (circular standard deviation with variable suffix `Ka`).

   (b) (line 238) Convert crystalline order parameter and patch wavelength ($\lambda$ for variable suffix `lam`) to randomization parameters in simulation (variable with suffix `pr` standing for positional randomization).

   (c) (line 263) Other parameters used in all simulations such as, number of cells simulated in periodic domain (`nCell = nX*nY`), cell lattice center randomization parameter (`randness`), Stokeslet regularization parameter (`reg`), dimensionless drag coefficient generated by each Stokeslet/ciliated cell (`dragCoeff`).

6. (line 270) Control parameters that can be changed depending on simulation objective include cilia coverage (`pCilia`), Monte Carlo trial number (`Ntrial`), number of simulated tracer particle (`Npar`), initial particle seed height away from cilia tip (`hpar`), allow particle motion in vertical direction (`track3D`), random walk time step (`dt`), and length of simulated track (`Nt`).

7. (line 658) Cell location generation and patch distribution function (See 1.2 and [14]).

8. (line 697) Main simulation loop (See 1.3). For each Monte Carlo trial and each set of input parameters, a Brownian particle simulation will run based on Euler Maruyama scheme. Generated particle tracks are analyzed based on mean displacement and Eulerian directionality measure computed from simulated Particle-Tracking-Velocimetry (PTV) velocity fields.

9. (line 922) Final results showing clearance per beat and mean clearance directionality as function of cilia coverage. Outputs will be saved to specified figure and data folders.

10. (line 957) Regularized Stokeslet algorithm based on [18] and other helper functions.
